# Modern speech recognition systems exhibit human-like word recognition but superhuman use of statistical dependencies between words

**DOI:** 10.64898/2026.11.17.767420

**Authors:** Annika Magaro, Gasser Elbanna, Erica Shook, Alex Kell, Mark R. Saddler, Josh H. McDermott

## Abstract

Advances in machine learning have yielded machine systems that approach human performance on real-world perception tasks. For speech recognition, it remains unclear whether such models’ behavior resembles that of humans. We addressed this question by measuring word recognition abilities in humans and models across 272 speech distortions, providing a detailed behavioral fingerprint. Three recent industry speech recognition models trained on hundreds of years of speech outperformed humans. However, in conditions where linguistic context was removed, they matched human performance well, suggesting that the models are superhuman in their ability to leverage statistical dependencies between words. Experiments manipulating model training data suggest that training data variability and realism contribute to the match to human perception. Overall, the results show that superhuman performance is now possible with systems that are optimized over enough data and suggest that accurate models of human speech recognition require optimization over appropriate tasks and amounts of speech.

## Introduction

Machine speech recognition is now accurate enough to be useful in daily life. In the past few years it has become common for humans to dictate text messages and emails to a smartphone or personal computer, and to rely on AI transcriptions of meetings. These advances have been fueled by artificial neural networks that learn to transcribe character strings or words from large datasets of annotated speech^1^. In parallel, the resulting systems have been shown to explain aspects of human brain responses to speech^2–4^, and these same machine learning methods have been used to build new models of various aspects of audition^5–12^.

In this paper we assess whether these systems can replicate human speech recognition behavior. Establishing the extent of the match between model and human speech recognition could be both practically and theoretically useful. An accurate model is a fundamental goal of perceptual science, and could form the basis of attempts to understand effects of hearing loss^13,14^, to derive improved prosthetics^10^, and to understand interactions between speech and language. Practically, the intelligibility predictions of such a model could be used to optimize speech signals or listening conditions for intelligibility. Fundamental discrepancies between deep learning models and humans could also have implications for the extent to which artificial neural networks trained with current methods are able to model perception.

Some reason to think that such systems might reproduce human-like speech behavior comes from the idea that humans are fairly well optimized to recognize speech. By this logic, a machine system optimized for the same problem might converge to similar behavior^15,16^. On the other hand, machine systems are trained with far more labeled data than humans plausibly receive^17,18^, or are optimized using loss functions or architectures whose biological plausibility is unclear, and these differences might be expected to be reflected in the resulting behavioral phenotype.

A few studies have compared performance of humans to that of speech recognition models tested on the same or similar experiments, both in the context of industry models^19–24^, and models built for scientific purposes that incorporate biological constraints^5,9,11^. These comparisons have revealed both nontrivial human-model similarities^5,9,11^ as well as apparent discrepancies^22,23^. However, many of these comparisons evaluated models on a small set of conditions, or involved systems that are no longer state-of-the-art, leaving it unclear whether current systems would provide a detailed match to human performance. Previous studies also left it unclear what factors influence a model’s match to human behavior. In particular, because the training data for industry systems are typically not publicly available, it is difficult to reproduce such systems outside the company that built them^25^, and to assess the role of the task, architecture, or training data characteristics in the model results.

We sought to answer two main questions. First, how well can artificial neural network models of speech recognition account for human speech recognition behavior? Second, what factors affect how well models match human behavior? To answer these questions, we first measured patterns of human speech intelligibility across a large battery of speech manipulations spanning both natural and artificial distortions. We then tested a set of speech recognition models on this battery, assessing the extent to which they qualitatively and quantitatively reproduced human performance.

We found that recent industry speech recognition systems mirrored the pattern of human behavioral trends but in absolute terms substantially outperformed humans. This superhuman performance was largely eliminated when excerpts of sentences were replaced with random word strings or isolated words, suggesting a superhuman ability to leverage linguistic context to predict words. This finding is consistent with recent findings that machine language models are better

than humans at next-word prediction^26–29^, but shows that this advantage has real-world consequences for understanding spoken communication. To understand the factors underlying the match to human word recognition, we trained our own in-house models (neural networks optimized to recognize words from excerpts of speech without the text decoders found in industry speech recognition systems). Manipulations of the training data showed that both the variability and realism of training data contributed to the extent of human-model performance match. Overall, the results suggest that we have entered an era where machine speech recognition systems have started to diverge from humans by outperforming them in many conditions. The results provide a cautionary note for using industry models as models of human perception, and indicate that models of human perception may require intentional engineering incorporating ecological constraints. The results also provide a comprehensive fingerprint of human speech recognition that could be used to benchmark future models.

## Results

To evaluate speech recognition abilities, we presented humans and models with speech excerpts and scored their transcriptions (Fig. 1a). To assess the extent to which model recognition matched or diverged from human recognition across a range of conditions, we processed the speech audio with 37 different types of distortions (Fig. 1b). These included both distortions present in the natural world (e.g. reverberation, whispering, background noise), distortions that are sometimes induced by modern electronic audio processing (e.g. clipping, filtering) and artificial distortions sometimes used in speech perception research (e.g. sine wave speech^30^, noise vocoding^31^, modulation filtering^32^). We used representative examples of as many distortion types as we could think of, including those tested in prior human-model speech recognition comparisons^5,9,22,23^. For most of the distortion types, we measured human and model performance (proportion correct) across different magnitudes of that distortion, for a total of 272 distortions.

**Figure 1.**
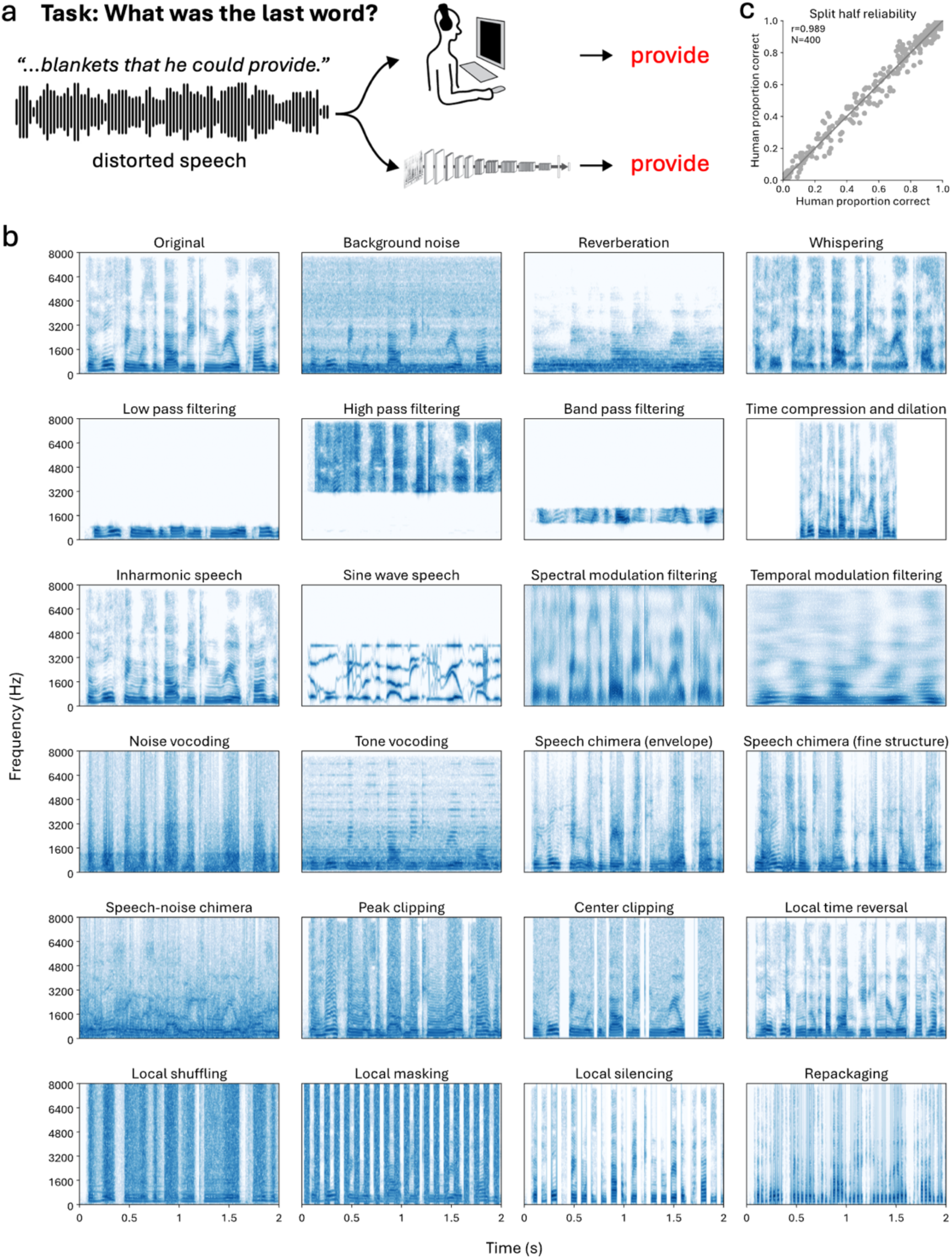
Experiment task and conditions. **a**. Example experimental task. Participants heard an audio clip and transcribed words (in this example, participants transcribed the last word they heard). Participants typed their responses. Models performed the same task on the same stimuli (they transcribed the whole excerpt but for this example experiment would be graded on the last word). **b**. Speech distortions. Panels show spectrograms of examples of each type of distortion included in experiments. For most distortions we tested several variants (e.g. different types of background noise, different filter cutoffs etc.). Due to space constraints, some distortion sub-conditions were excluded from this figure. **c**. Split-half reliability of human performance across distortion conditions for the task shown in panel a.

**Table 1:** Distortions included in experiment battery.

| Distortion Type | Distortion Settings | Description |
|---|---|---|
| Background noise | Five types of noise; SNRs of -9, -6, -3, 0, 3 dB | Speech was superimposed on music, auditory scenes, 8-talker babble, speech-shaped noise, or amplitude-modulated speech-shaped noise. |
| Background textures | 43 sound textures; fixed SNR of -3 dB | Speech was superimposed on synthetic sound textures matched to the statistics of one of 43 real-world texture recordings<sup>9,33</sup>. |
| Reverberation | Direct-to-reverberation ratios of 20 or 50 dB, RT60s of 200, 400, 800, and 1600 ms, applied to clean speech or speech in babble noise | Speech was convolved with synthetic room impulse responses, derived from the statistics of real-world room impulse responses<sup>34</sup>. |
| Whispering | Single condition | Speech was resynthesized with noise excitation intended to simulate whispering<sup>35</sup>. |
| Low-pass filtering | Cutoffs: 400, 800, 1600, 3200 Hz | Speech was convolved with a finite-impulse-response filter. |
| High-pass filtering | Cutoffs: 400, 800, 1600, 3200, 6400 Hz | Speech was convolved with a finite-impulse-response filter. |
| Band-pass filtering | Center frequency of 1500 Hz; bandwidths of 5, 10, 20, 30, and 40 semitones | Speech was convolved with a finite-impulse-response filter. |
| Time compression and dilation | 0.5x, 0.75x, 0.875x, 1.0x, 1.25x, 1.5x, 2.0x | Speech was time stretched (changing the speech rate while preserving the f0 and vocal tract filtering) using PRAAT. |
| Inharmonic speech | Frequencies jittered by up to 0, 10, 30, or 50% of the f0 | Speech was resynthesized with inharmonic excitation, obtained by jittering the frequencies of individual harmonics and then recombining them with an estimate of the original time-varying spectrotemporal envelope<sup>35</sup>. |
| Sine-wave speech | Single condition | Speech replaced with four frequency-modulated sinusoids whose frequencies followed the estimated trajectories of the first four formants<sup>30</sup>. |
| Spectral modulation filtering | 0.5, 1, 2, 4, 8 cycles/kHz | A spectral lowpass filter was applied in the modulation domain, attenuating spectral modulations above a specified cutoff before reconstructing the filtered signal<sup>32</sup>. |
| Temporal modulation filtering | 3, 6, 12, 24 Hz | A temporal lowpass filter was applied in the modulation domain, attenuating temporal modulations above a specified cutoff before reconstructing the filtered signal<sup>32</sup>. |
| Noise vocoding | 1, 2, 4, 8, 16, 32 channels | Speech was decomposed into ERB-spaced frequency channels. The subband from each channel was replaced by a noise signal with the same envelope. The modulated noise signals were summed to produce the final vocoded waveform<sup>31</sup>. |
| Tone vocoding | Vocoded above subband 0, 8, 16, 24, and 32 (out of 32 total); superimposed on stationary or modulated noise; SNRs of -6, -4, -2, 0, 2, 4, and 6 dB (stationary) or -18, -15, -12, -9, -6, -3, and 0 dB (modulated) | Speech was decomposed into subbands. Subbands above the cutoff were replaced with a tone at the center frequency of the subband modulated by the subband envelope, and then recombined. The resulting signal was superimposed on speech-shaped noise. |
| Chimeras | Speech-speech and speech-noise chimeras; 1, 2, 4, 8, and 16 subbands | Speech and/or noise signals were decomposed into subband envelopes and fine structure. The fine structure of one signal was combined with the envelopes of the other signal<sup>36</sup>. |
| Peak clipping | Threshold (percentile): 0, 0.25, 0.5, 0.75, 0.9, 0.98 | All samples in the signal that were above the threshold for that signal were clipped to that threshold<sup>37</sup>. |
| Center clipping | Threshold (percentile): 0, 0.25, 0.5, 0.75, 0.9, 0.98 | All samples in the signal that were below the threshold for that signal were set to zero<sup>37</sup>. |
| Local time reversal | Segment lengths: 10, 20, 30, 40, 50, 100, 150, 200 ms | Speech clips were divided into adjacent non-overlapping segments. Each segment was then time-reversed, and the reversed segments were concatenated to yield the final signal<sup>38</sup>. |
| Local shuffling | Segment lengths: 1/8, 1/4, 1/2, 1, 2, 4 ms | Waveform samples within segments of the specified length were randomly reordered<sup>39</sup>. |
| Local masking | SNRs of -18, -9, 0, 9 dB crossed with segment lengths of 1, 3, 10, 30, 100 ms | Speech was masked with a square-wave modulated Gaussian white noise<sup>40</sup>, whose period was equal to the segment length. |
| Local silencing | Segment lengths of 1, 3, 10, 30, or 100 milliseconds crossed with silenced proportions of 1/4, 1/2, or 3/4 | Speech was multiplied by a rectangular wave envelope, silencing a fraction of the waveform in every segment of the specified length<sup>40</sup>. |
| Repackaging | Silent intervals of 0, 10, 20, 40, 80, or 160 ms | Speech was time-compressed by a factor of 2. Some amount of silence was then inserted every 40 ms<sup>41</sup>. |

We designed the experiments with the intent of eliminating uninteresting reasons for human-model differences in performance. First, to eliminate potential effects of rarely encountered words, we used sentences that contained a word from a 230-word vocabulary of common English words, and then for most analyses scored only that word. Second, to eliminate any possibility of using speech material present in model training data (which nowadays tends to combine many different sources of publicly available recorded speech), we avoided existing speech corpora when generating stimuli. Instead, we generated sentences and rendered them using speech synthesis and voice cloning (see Methods; similar results were obtained in pilot experiments with stimuli based on recorded speech; see Supplementary Figures 1-2).

### Comprehensive benchmark of human speech recognition

Human performance varied across the distortion conditions in ways that were consistent with prior results. For example, jittering the frequencies of constituent harmonics did not compromise intelligibility, while drastically reducing spectral resolution by noise vocoding speech with few channels compromised intelligibility significantly. See Supplementary Figure 3 for plots of results for individual distortions for the experiment where humans transcribed the last word of the speech excerpt. The variation in human performance was highly reliable across splits of human participants. Fig. 1c shows this variation for the experiment in which humans transcribed the last word of the speech excerpt (split-half reliability: r = 0.989, p<.001). See Supplementary Figure 4 for split-half reliability of all experiments.

### Industry speech recognition systems

We tested five industry speech recognition models whose architectures and trained weights were publicly available. Three of these models were recently developed state-of-the-art systems (Whisper-large-v3^42^, Parakeet-TDT-0.6B-v2^43^, and SeamlessM4T-v2^44^; the latter two are henceforth referred to as Parakeet and Seamless for brevity) and two of them were earlier versions of one of these systems (Whisper-large-v1, Whisper-tiny-v1). Parakeet was the top-ranked model on the Open ASR leaderboard during time of testing^45^. The two earlier models were included to help assess whether the human-similarity of these models is consistent across different scales of model architectures and training data. Models transcribed the entire excerpt but were scored using only the word on which humans were evaluated. We initially present results for the three most recent systems (Whisper-large-v3, Parakeet, and Seamless).

**Table 2:** Listing of models, number of parameters, and amount of training data.

| Model name | Architecture | Number of parameters (million) | Input representation | Task objective | Training hours (million) |
|---|---|---|---|---|---|
| Whisper-tiny-v1 (OpenAI) | Transformer-based encoder-decoder model (4 encoder + 4 decoder layers) | 39 | 80-channel Mel spectrogram | seq2seq next token prediction | ~0.68 |
| Whisper-large-v1 (OpenAI) | Transformer-based encoder-decoder model (32 encoder + 32 decoder layers) | 1550 | 80-channel Mel spectrogram | seq2seq next token prediction | ~0.68 |
| Whisper-large-v3 (OpenAI) | Transformer-based encoder-decoder model (32 encoder + 32 decoder layers) | 1550 | 128-channel Mel spectrogram | seq2seq next token prediction | ~5 (~1 labeled + ~4 pseudo-labeled) |
| SeamlessM4T-v2 (Meta) | Speech encoder: wav2vec-BERT 2.0 24 Conformer blocks<br>Text decoder: 24 transformer layers | 2326 (669 for encoder) | waveform | self-supervised pretraining + supervised finetuning | ~4.5 (pretraining) + ~0.044 (finetuning) |
| Parakeet-TDT-0.6B-v2 (NVIDIA) | 24 FastConformer encoder layers + TDT decoder | ~1100 (~600 for encoder) | waveform | self-supervised pretraining + supervised finetuning | ~0.12 (pretraining) + ~0.0005 (finetuning) |

### Experiment 1: 2-second coherent sentence excerpts (last word)

We first compared models vs. humans on excerpts of coherent sentences in which the target word was at the end. To eliminate potential effects of memory limits in humans, in this experiment we asked humans to only transcribe the last word in the excerpt. Here and elsewhere, as an overall measure of human-model similarity, we compared model and human performance across the 277 experiment conditions (five of the stimulus conditions yielded two separate experimental conditions as they were scored in two ways; see Methods). We quantified the similarity using Pearson correlation coefficient (R) as well as the average difference in performance across conditions (Δ, where Δ>0 indicates a model over-performed humans and Δ<0 indicates a model under-performed humans across the battery). We evaluated the statistical significance of the human-model performance difference via a Wilcoxon signed-rank test. As shown in Figure 2a, all 3 models exhibited performance trends that were strongly correlated to those of humans (Whisper-large-v3: r=0.94, r^2=0.88, 95% CI [0.85, 0.91]; Parakeet: r=0.96, r^2=0.91, 95% CI [0.89, 0.93]; Seamless: r=0.95, r^2=0.90, 95% CI [0.87, 0.93]; all p<.001), but all 3 models tended to be superhuman in absolute terms (p<.001 for Whisper-large-v3 and Parakeet; p=.002 for Seamless). See Supplementary Figure 5 for results color-coded by distortion type.

### Experiment 2: single words

As an additional test, we ran an experiment in which the stimuli were single words. As shown in Figure 2b, model performance for single words was reduced relative to that for humans (Δ was significantly lower for all three models, p<.001; via bootstrapping over the 277 conditions, see Methods for details; all comparisons of model-human performance differences between experiments use this test). This effect was particularly pronounced for Seamless, which also tended to hallucinate extended strings of words despite the presence of a single word in the stimulus. This result is consistent with the idea that the models might be strongly benefitting from linguistic context but could also be explained by isolated spoken words being out-of-distribution for the models.

### Experiment 3: 2-second random word sequences (last word)

To further investigate the effect of linguistic context on human and model performance, we pseudo-randomly scrambled the non-target words of the stimuli from Experiment 1 so that they no longer formed coherent sentences (see Methods). As in Experiment 1, audio was cropped to 2 seconds, and humans transcribed the last word while models transcribed the entire excerpt but were graded based on only the last word. As shown in Figure 2c, all 3 models matched human performance well in this condition, with Whisper slightly outperforming humans (Δ=0.07, p<.001) and Seamless slightly underperforming humans (Δ=-0.06, p<.001). The model-human performance difference was reduced compared to Experiment 1 for all 3 models (p<.001 for all 3 models, via bootstrap over the 277 conditions), consistent with the model-human performance difference in Experiment 1 being driven by a superhuman ability to leverage linguistic context. In addition, the model-human performance difference was increased compared to Experiment 2 for all 3 models (p<.001 for all 3 models), indicating that the models perform particularly poorly for single words (plausibly because they are optimized for continuous speech). As shown in Supplementary Figures 6-9, humans did not have significantly different performance in Experiment 2 and 3, while all models did.

### Experiment 4: 2-second coherent excerpts (first word)

As a further test of the role of linguistic context, we ran an experiment in which the target word was the first word of the speech excerpt. The rationale was that the industry models we tested are acausal, such that they should be able to use the words both before and after the target word to constrain their judgments of the target word. By contrast, human language processing is plausibly more sequential, relying more on the past than the future^46,47^. It thus seemed that models might be more superhuman when the first word of the excerpt was scored rather than the last word. As shown in Figure 2d, the model-human performance difference increased for all 3 models when evaluated based on the first word (p<.001 for all 3 models), and all 3 models significantly out-performed humans (p<.001 for all 3 models). This result is further consistent with these models having a superhuman ability to use linguistic context when recognizing words.

### Experiment 5: 2-second coherent sentence excerpts (middle word)

To rule out the possibility that the model performance advantage was specific to words at the beginning or end of a speech excerpt, we ran an experiment in which the target word was located in the middle of the speech excerpt (i.e., overlapping the one-second mark of the excerpt). To obtain transcriptions of the middle word, we asked human participants to transcribe all the words in the excerpt and scored the middle word (Experiment 5).

As shown in Figure 2e, models again significantly outperformed humans (p<.001 for all 3 models). In addition, the model-human performance difference was larger than that for the last-word experiment (p<.001 for all 3 models), as might be expected given the models’ acausal processing.

To rule out the possibility that human performance on the full-transcription task of Experiment 5 was limited by working memory demands (from having to hold all the words in memory while typing them), we directly compared performance when participants had to transcribe all the words of an excerpt to when they just had to transcribe one word. For this purpose we conducted a new experiment (Experiment 1b) using the same stimuli as in Experiment 1. Participants transcribed all the words in the excerpt but we scored only the last word. Human performance was similar in these two experiments (1 and 1b; p=.102, Wilcoxon test; see Supplementary Figure 10 for Experiment 1b results). This result indicates that memory limitations did not substantially constrain performance in this setting (with two-second excerpts of coherent sentences).

**Figure 2.**
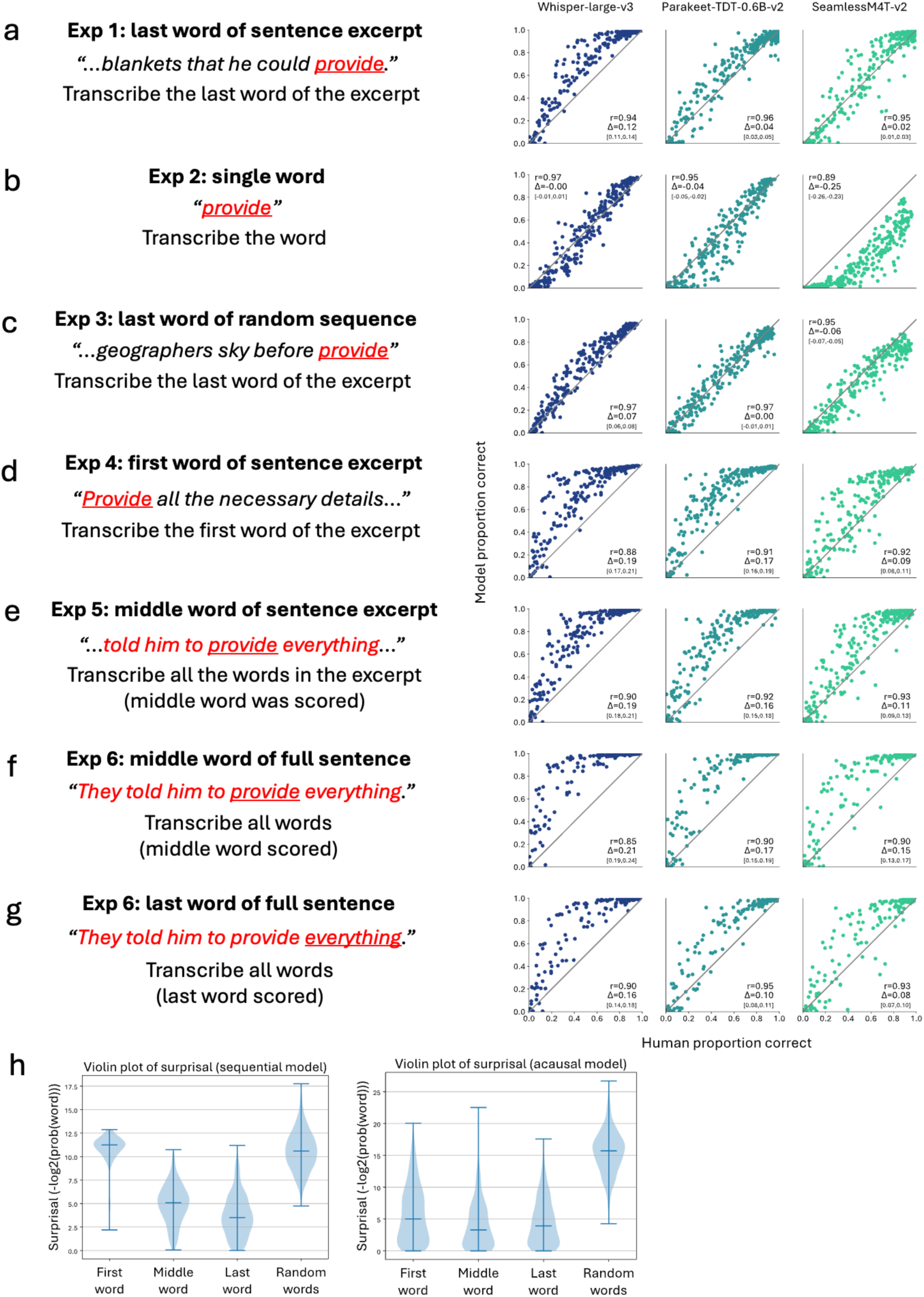
Summary of results for industry models and humans. **a-g**. Comparison of human and model speech intelligibility for industry models for Experiment 1-6. Red font denotes response transcript; underlined word was the word scored from the response transcript. Each dot plots intelligibility of one of the 277 distortion conditions. Average model-human difference is denoted by Δ, with 95% confidence intervals bootstrapped 1000 times in brackets below. **h.** Violin plots of surprisal analysis for sequential model (GPT-2) and bidirectional model (BERT).

### Experiment 6: Full sentences

To control for the possibility that presenting excerpts of sentences could be differentially hurting performance for humans and/or models, we ran an additional experiment in which humans and models transcribed all the words of entire sentences (i.e., not clipped to 2 seconds; durations ranged from 2 to 3.5 s). Stimuli were generated from the sentences used for Experiment 5, and we scored performance in three ways: first, based only on the middle word, second, based only on the last word, and third, based on the fraction of all words transcribed correctly. As shown in Fig. 2f-g and Supplementary Figure 10, the model-human performance difference persisted for full sentences and was somewhat larger than in the previous experiments using 2s excerpts. The increase in superhuman performance for the middle vs. the last word was again apparent (p<.001 for all 3 models). Thus, the superhuman performance is not specific to excerpts of speech and persists for full sentences.

### Surprisal analysis

To further substantiate potential differences in predictability of words at different positions in our stimuli and the possible interaction with the directionality of linguistic analysis, we calculated surprisal of the target word for the first, middle and last word stimuli using both a sequential surprisal model (GPT-2^48^) and an acausal surprisal model (BERT^49^). Intuitively, surprisal measures how surprising a word is in the context of the rest of the sentence (quantified as the negative log-probability of a word given the context). As shown in Fig. 2h, when evaluated with a sequential model, surprisal values were substantially lower for the last word than for the first and middle words, on average (i.e., the last word was more predictable than the first or middle word given the other words in the excerpt, p<.001). By contrast, when evaluated with an acausal model, surprisal values were similar for words at different positions, though there were modest differences (the middle word had slightly lower surprisal values than the last word, p=.03, and the first word had slightly higher acausal surprisal values than the last word, p<.001). These effects parallel the recognizability of words at different positions for humans and models (humans were better for the last word whereas models performed comparably across word positions; Experiments 1 vs. 4 and 5). The results are thus further consistent with the idea that human-model differences result from differences in the ability to use linguistic context, with humans mostly relying on past words to make word predictions, such that the models outperform humans by a larger degree when evaluated on words in the middle of sentences.

### Human-human and model-model comparisons

To further illuminate the effect of the different experimental conditions on humans and models, Figure 3 directly compares performance between the different experiments. The first two rows compare the two conditions without linguistic context (Experiments 2 and 3) to Experiment 1 (coherent sentence excerpts). Humans show a modest benefit of linguistic context, as expected from many prior studies of degraded speech^50–52^. However, this benefit is larger for the models. These comparisons also reveal that humans perform better on the last word (Experiment 1) than the first or middle word (Experiments 4 and 5; third and fourth rows of Figure 3), whereas models show little difference. These effects support the idea that models benefit from the bidirectional use of context, whereas humans do so to a lesser extent. The models also show better performance for full sentences compared to two-second excerpts (last two rows of Figure 3). This latter result is consistent with the models benefitting from the increase in linguistic context in full sentences, but could also reflect the fact that full sentences are presumably more similar to the bulk of the model training data. See Supplementary Figures 6-9 for all human-human and model-model comparisons. See Supplementary Figure 11 for results on only the words common to all experiments.

**Figure 3.**
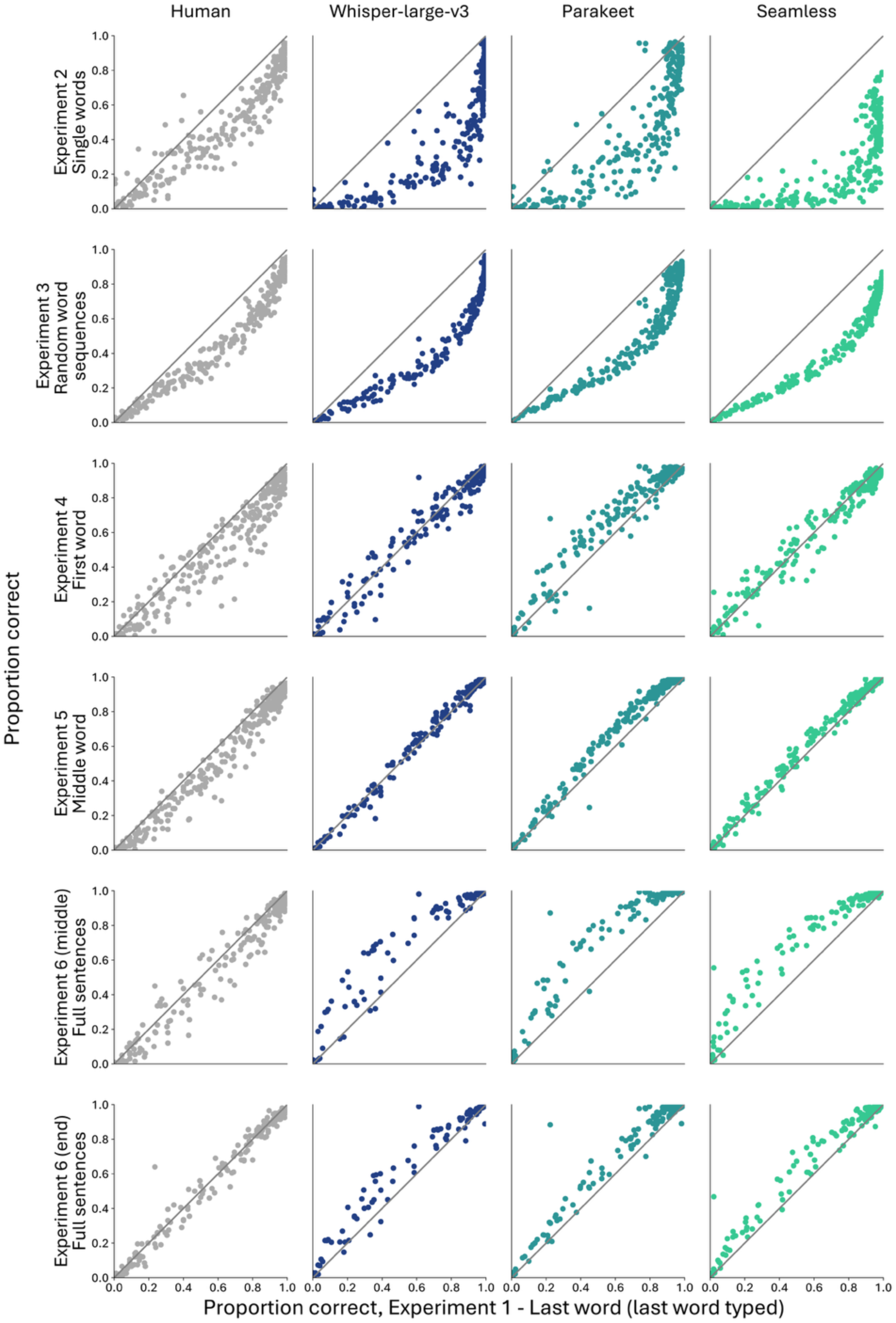
Comparison of performance by humans and models across experiments. Each dot plots intelligibility of one of the 277 distortion conditions. Data is the same data shown in Figure 2, plotted differently to expose effects within humans and models. x axis of each graph plots performance in Experiment 1. y axis plots performance in one of the other experiments (a different experiment for each row of the figure). The first two rows highlight the effect of linguistic context, comparing performance for coherent sentence excerpts to that for single words and random word sequences. Humans show a benefit from linguistic context, as expected, but this effect is more pronounced in the industry models. The third and fourth row compare performance for the last word to that for the first and middle word, respectively. The last two rows compare performance for sentence excerpts to that for full sentences.

### Overall match to human performance

Figure 4 shows individual results graphs for each type of distortion for Experiment 5, comparing humans to Whisper-large-v3 (see Supplementary Figures 12-20 for analogous results figures for the other experiments and models; we chose to show one model and one Experiment here for clarity of presentation). Nearly every panel exhibits the two main effects evident in the aggregate results in Figure 2. First, Whisper-large-v3 tends to reproduce the qualitative pattern of performance seen in humans across the variations of each distortion. Second, Whisper-large-v3 tends to produce overall higher performance than humans. The main exceptions involve instances of ceiling effects (where the model performs close to 100% across conditions that produce performance variation in humans). The one qualitative discrepancy occurs for one of the local masking manipulations (SNR=-9), for which Whisper exhibits a more pronounced “V” pattern of performance than humans (row 5, column 1).

The extent of superhuman performance varied somewhat across distortions. One striking specific example occurred for sine-wave speech^30^, a manipulation in which speech is replaced with frequency-modulated sinusoids that trace out the formants of the original speech (row 2, column 5, left set of bars). As expected, we found that humans could recognize a fraction of stimuli, getting around 40% correct. The fact that human recognition is above-chance is itself remarkable, as has often been noted^30^, as the stimulus retains little of the superficial acoustic structure of speech. However, Whisper-large-v3 recognized over 90% of words correct.

**Figure 4.**
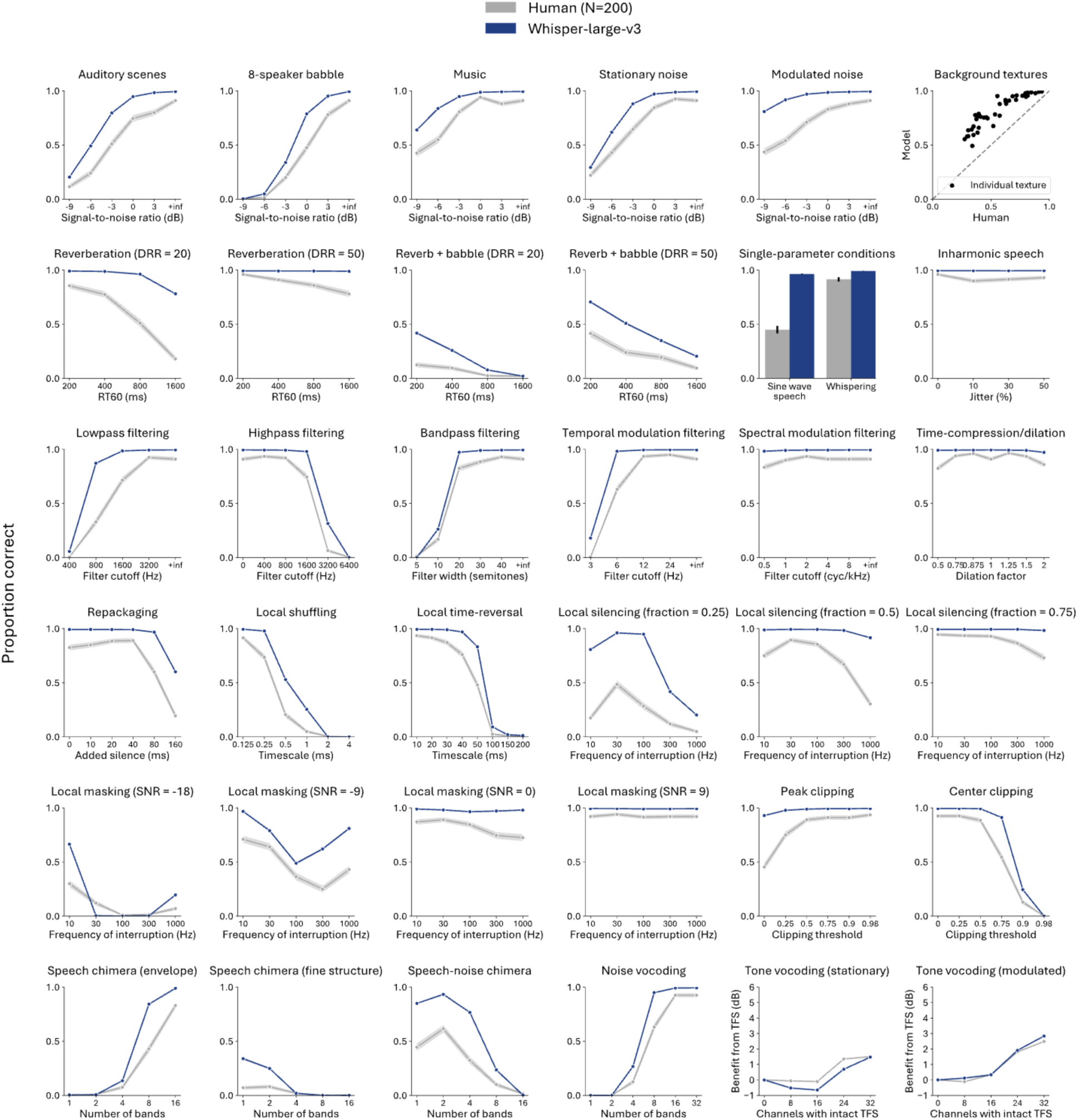
Comparison of Whisper-large-v3 and human (N=200) performance for each type of distortion for Experiment 5 (middle word task). Error bars plot SEM, obtained via bootstrap.

### Model properties influencing superhuman performance

To clarify the factors underlying superhuman speech recognition, we also tested two earlier versions of the Whisper model family (Whisper-large-v1 and Whisper-tiny-v1). Whisper-large-v1 has a similar architecture to Whisper-large-v3 but a training set that is around 8x smaller. Whisper-tiny-v1, by contrast, has an architecture that is around 40x smaller (but is trained using the same training data as Whisper-large-v1, around 8x smaller than Whisper-large-v3)^42^.

We found that Whisper-large-v1 performed similarly to Whisper-large-v3, though with somewhat less superhuman performance (s=0.11 vs. s=0.12, significantly different from each other; p<.001; Supplementary Fig. 21). However, Whisper-tiny-v1 dramatically underperformed Whisper-large-v1, and substantially underperformed humans (Δ=-0.18; p<.001). These comparisons obviously do not include comprehensive variation of architectures and training data, but suggest that large model architectures contribute to the superhuman ability to model word dependencies. We note also that even these earlier versions of Whisper were trained on enormous amounts of audio (if played continuously, the training data would amount to over 77 years of speech). See Supplementary Figures 21-32 for results analogous to Figures 2a-g, 3, and 4 for the earlier versions of the Whisper model family.

### In-house models

Because the training data for industry models is typically not publicly available, and because training such models requires considerable computing resources, it is difficult to perform architecture and/or training data manipulations (as might be useful to understand the basis of any similarity to humans^6,7^). To manipulate these factors more effectively, we built our own in-house models.

We tested two types of architectures: purely convolutional architectures (CNNs) and Conformer^53^ architectures (CF) (Fig. 5a&b). The CNNs were derived from previous searches conducted in our lab for architectures that performed well on word recognition tasks^9^, and differed from each other by the addition or subtraction of model stages and/or changes on filter kernel sizes or pooling ratios. There were 9 convolutional architectures, and results for convolutional models were generated by averaging the results for the 9 architectures.

We trained the conformer architecture after finding in parallel work^14^ that it performed well on the word recognition task. The conformer architecture consisted of 4 convolutional layers followed by 4 convolution-augmented transformer layers. Figure 5c summarizes the in-house model architectures.

We trained the in-house models to perform word recognition using a dataset of speech superimposed on background noise (around 1100 hours of speech). Speech excerpts were taken from the Common Voice dataset^54^. Background noises were taken from the AudioSet dataset^55^. We attempted to make the dataset as variable as possible subject to the constraints of using natural recorded speech and real-world background recordings, so we included as many unique talkers as we could (tens of thousands). The models were trained to recognize the word at the middle of a 2-second clip (i.e., overlapping the 1s mark), motivated by the use of this task in previous models that accounted for aspects of human word recognition^5,9,11,56,57^. The evaluation experiments with distorted speech used the stimuli from Experiment 5 (in which the target word was positioned at the middle of the excerpt). We then tested various manipulations of the training data to see how they affected the match to human performance. These manipulations are described in Figure 5d.

**Figure 5.**
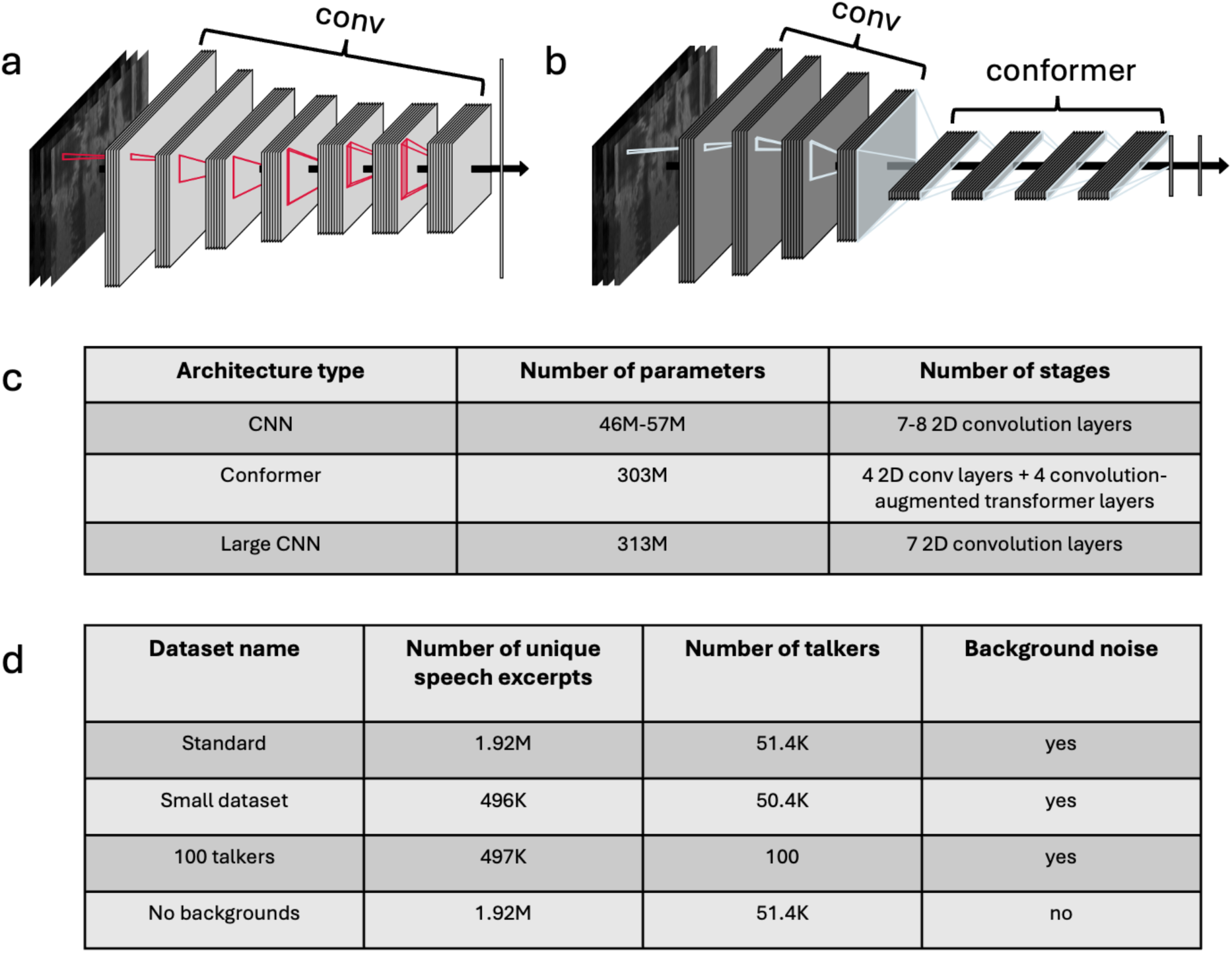
In-house model architectures and description. **a**. CNN architecture schematic. **b**. Conformer architecture schematic. **c**. Description of model architectures for 3 main architectures. **d**. Description of dataset creation and manipulations.

### In-house models modestly match human performance on Experiment 5

We refer to the CNN and Conformer models trained on the dataset described above as the Standard CNN and Standard Conformer, respectively. We found that both models matched the pattern of human performance well, though not as well as the industry models (CNN: r=0.90, p<0.001, r^2=0.81, 95% CI [0.74, 0.87]; CF: r=0.91, p<.001, r^2=0.82, 95% CI [0.77, 0.87]). The Standard Conformer tended to be slightly superhuman overall (Δ=0.08, p<.001; Fig. 6a). See Supplementary Fig. 33 for results for a CNN model whose architecture was matched in size to that of the Conformer (the performance difference persisted), and Supplementary Figures 34-37 for results for individual distortion types. See below for discussion of conditions where models substantially underperformed humans.

### Model with reduced training set variability shows worse match to humans

To first assess whether human-model similarity depended in part on the richness of the model’s training data, we trained models on a training set containing less speech variability, achieved by reducing the number of unique talkers to only 100. As shown in Fig. 6a, the resulting 100-talker models exhibited a worse match to humans (CNN: r=0.85, p<.001, r^2=0.73, 95% CI [0.65, 0.81]; CF: r=0.86, p<.001, r^2=0.75, 95% CI [0.67, 0.82]; significantly lower than the human-model similarity of the corresponding Standard model (p<.001 for both CNN and CF, assessed via bootstrap over the 277 conditions). The decreased similarity was driven in part by decreased robustness to highly distorted speech.

Reducing the number of talkers forced us to cut the total dataset size to approximately 25% of the size of the original dataset. To eliminate this confound, we trained an additional CNN and Conformer model on a dataset with the same number of unique talkers as the original dataset but with a total dataset size matched to that of the 100-talker dataset. As shown in Fig. 6a, each Small-dataset model exhibited a modestly better match to humans than the corresponding 100-talker model (p<.001 for both CNN and CF), and only a slightly worse match to humans than the corresponding Standard model (CNN: p<.001, CF: p=.016). These differences in human-model similarity were mirrored by differences in overall robustness (Standard better than Small-dataset, p<.001; Small-dataset better than 100 talker, p<.001). This result suggests that the variability in the training data (along with the overall amount of unique examples) contributes to human-model similarity (largely via their effect on robustness), though the effects of the specific manipulations used here were modest.

### Model trained without background noise shows worse match to humans

As a second manipulation of training data richness, we trained models without any background noise (No Backgrounds). The training data otherwise consisted of the same speech excerpts as were used for the standard models. The resulting models were less robust overall than the Standard models (Fig. 6a), even on conditions that didn’t involve background noise, yielding a worse match with humans (CNN: r=0.74, p<.001, r^2=0.54, 95% CI [0.48, 0.60]; CF: r=0.84, p<.001, r^2=0.70, 95% CI [0.65, 0.75]; significantly lower than the human-model similarity of the corresponding Standard model: p<.001 (CNN and CF), assessed via bootstrap over the 277 conditions).

These effects of the training data were evident in individual experiment results. For instance, the 100-talker model performed worse than the standard model (and worse than humans) on sine-wave speech and on speech with moderate amounts of local silencing (Fig. 6b). The No backgrounds model unsurprisingly performed worse for speech in different types of background noise but was also less robust in reverberation (Fig. 6c).

**Figure 6.**
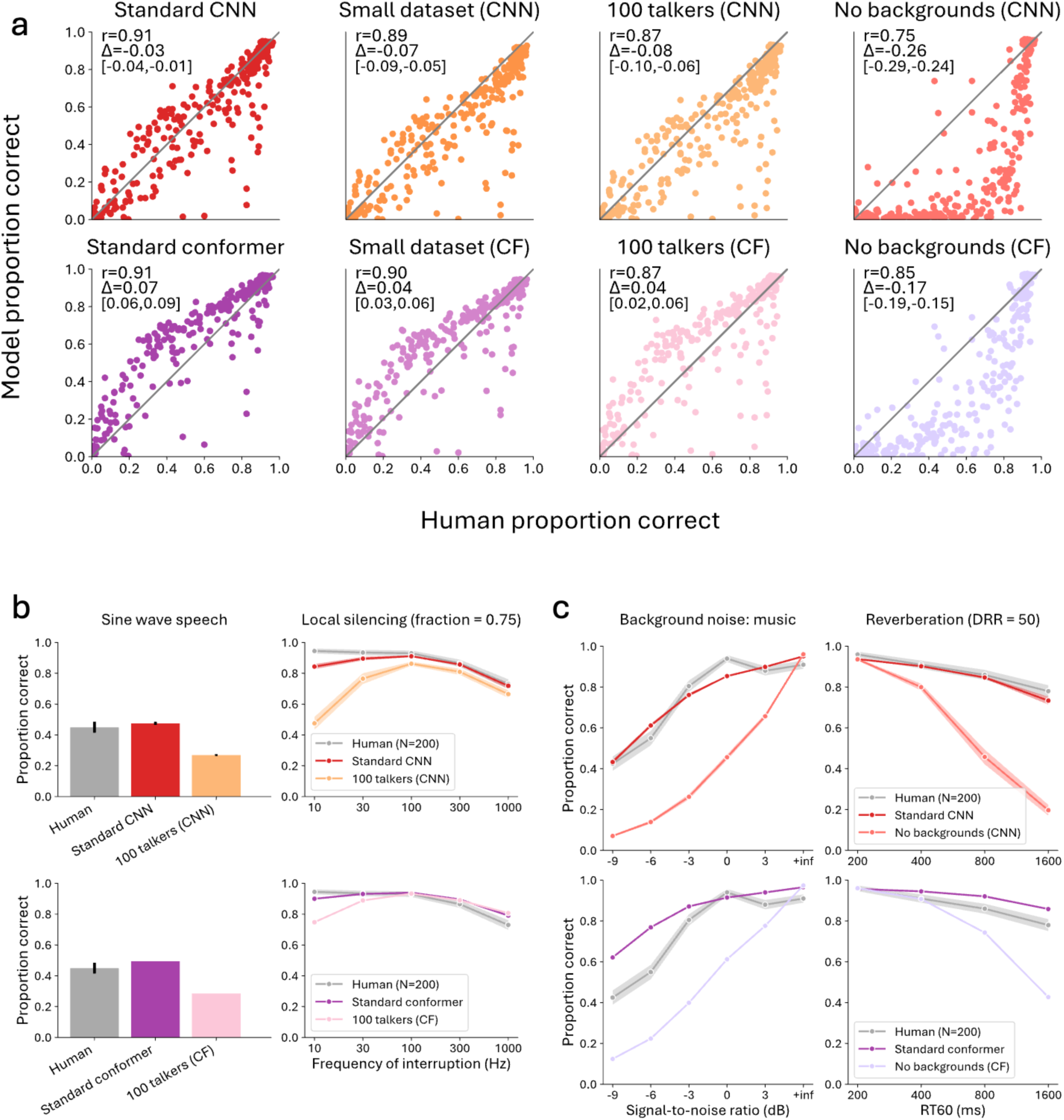
Results of in-house model training data manipulations. **a**. Summary results for 8 in-house models. Each dot plots intelligibility of one of the 277 distortion conditions. **b**. Comparison of human and model speech intelligibility on conditions where 100 talker models have worse match to humans than the Standard models. **c**. Comparison of human and model speech intelligibility for conditions where No backgrounds models have worse match to humans than the Standard models.

### Optimizing on discrepant conditions yields better match to humans on those conditions

The Standard in-house models produced better quantitative matches to human speech intelligibility than the other in-house models we tested but nonetheless exhibited some substantial discrepancies with humans. To better understand the source of these discrepancies, we trained an additional in-house CNN and Conformer model with the most discrepant conditions explicitly included in the training data (by processing some of the training speech excerpts with the distortion manipulations). These Augmented models were intended to address whether the poor model performance on some conditions reflected a limitation of the model architecture. The included distortions, when evaluated as part of the test battery, elicited Standard model performance that was substantially worse than humans, and that in some cases deviated from the pattern of performance seen across different variants of the distortion (e.g. filter cutoff, or silencing rate; Fig. 7). The distortions comprised time-compression and dilation, bandpass and highpass filtering with a wide range of cutoffs, and local silencing using a range of rates. Each distortion augmentation was applied independently to a modest fraction of training examples (Fig. 7a; see Methods).

As shown in Fig. 7b, training on the discrepant conditions substantially reduced the discrepancies with humans for all the conditions (by improving the model’s performance). In particular, for the audio filtering, time compression, and repackaging distortions, the models matched human performance almost exactly (Fig. 7c). However, for the local silencing manipulation, the models became strikingly superhuman (Fig. 7d). These results indicate that recognition performance is not fundamentally limited by the model architecture, in that explicitly training on difficult conditions boosted performance. However, they do not rule out the possibility that the model architecture limits the ability to generalize from naturally occurring speech to the unnatural forms of distortion tested here. See Supplementary Figure 38 for all in-house model vs. in-house model comparisons.

**Figure 7.**
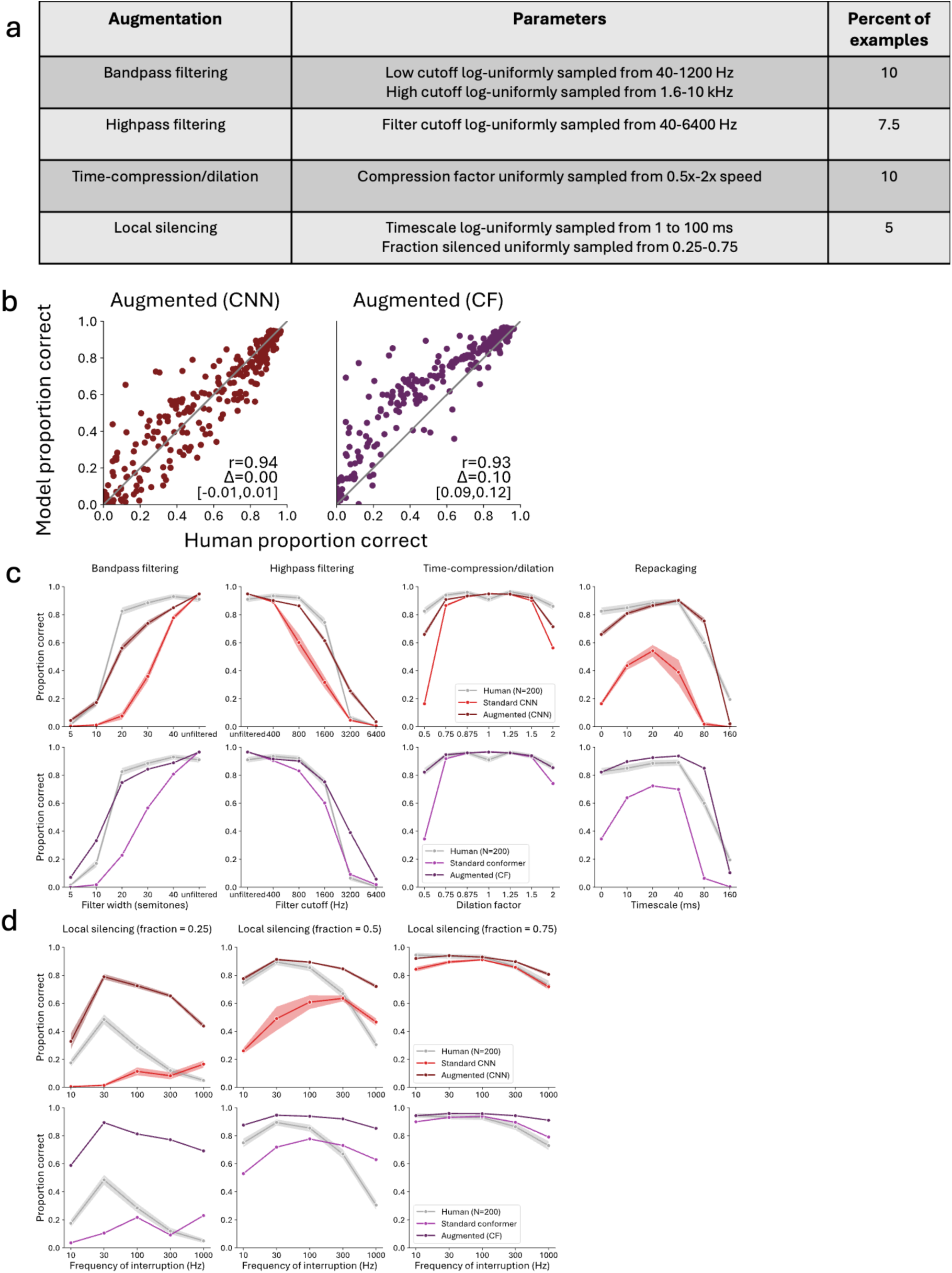
Augmented model results. **a**. Description of augmentations applied to the training set. **b**. Summary scatter plots of human vs. Augmented model performance. **c**. Performance of Augmented models vs. Standard models vs. humans on four experiment conditions for which the augmentations produced notable improvements in performance. **d.** Performance of Augmented models vs. Standard models vs. humans on local silencing (for which the augmentations caused models to substantially outperform humans).

## Discussion

We conducted a large-scale comparison of human and machine word recognition abilities. We measured the intelligibility of speech across 272 types of distortion, some of which greatly compromised human intelligibility and some of which did not, providing a “fingerprint” of human speech recognition. We then made analogous measurements of machine speech recognition models, testing them on the same stimuli and task for which humans were evaluated. We tested recent industry models as well as a set of in-house models for which we could manipulate the training data and architecture. Although industry models tended to mirror the qualitative pattern of human performance, they also tended to exhibit superhuman performance. Experiments manipulating the presence of linguistic context suggested that this advantage relative to humans primarily reflects a superhuman ability to use statistical dependencies between words, rather than superhuman abilities to discriminate individual words. Current industry models thus appear to capture aspects of human word recognition but diverge from humans in how language influences word recognition. This conclusion is consistent with recent work comparing humans to language models that operate on text. However, the results go beyond text-based human-model comparisons by showing that machine linguistic advantages have substantial consequences for spoken communication in challenging conditions. To explore the factors influencing model-human alignment, we built a set of in-house models and varied the training data and architecture. Reducing the variability (the number of talkers) or realism (the presence of background noise) of training data yielded models with worse matches to humans, supporting the hypothesis that the human behavioral phenotype is shaped by the demands of real-world speech recognition, and that the high alignment of industry models to humans (once linguistic factors are discounted) could be driven by the models having been optimized over vast amounts of data.

### Similarities and divergence of industry speech systems compared to humans

The industry speech recognition systems we tested showed strong alignment with humans once contextual information from surrounding words was removed (r=0.97 for Whisper-large-v3 and Parakeet, only just below the noise ceiling), along with striking overperformance in conditions where surrounding words could be used to help infer the target word. Both of these effects plausibly reflect the amount of training data such systems are optimized for, which has steadily increased over previous decades^42,58^. The most recent models we tested were exposed to hundreds of years of speech (Fig. 2a), with even the older models having been trained on 77 years. It is difficult to estimate the effective “training set” that shapes a human’s speech recognition abilities (through some combination of evolution and development). However, much of the robustness of human speech recognition may be general-purpose. For instance, noise invariance is plausibly not specific to speech, and rather may reflect a general ability to separate stationary “background” signals from less stationary signals^59–62^. The effective training set that underlies such general-purpose recognition robustness in humans could potentially be quite large, reflecting the evolutionary history of human perceptual systems long before the arrival of spoken language. It could be that the large-data regimes in which current industry systems are trained are not unreasonably larger than the scale of the data that shaped this general-purpose robustness. By contrast, the effective training set that underlies speech and language abilities in humans could be considerably smaller than what the most recent machine systems are being trained on^17,18^, allowing the models to be much better than humans at predicting likely word sequences. Thus, both the alignment to humans and the superhuman ability to leverage word dependencies could reflect the scale of the training data.

The results also revealed clear limitations of the industry models we tested. For instance, some of the models performed much worse than humans when tested on isolated words. This is plausibly due to these models having been optimized for continuous speech.

The results provide a cautionary note regarding the use of industry models as models of perception. Building deep learning models is resource-intensive, and one might suppose that scientists would be well served by leveraging the efforts in large companies, adopting industry systems as candidate models of human perception and/or sensory systems. However, industry systems are not built to replicate what happens in the brain, and the present results provide some evidence that in the domain of spoken language they have started to diverge from human levels of performance. At the same time, the results provide clues that the large-scale training typical of industry settings may be helpful in reproducing aspects of human abilities.

### Relation of human-model similarity to training conditions

The underlying causes of the human-model similarity seen for industry systems remains speculative due to uncertainty over the model training data, which was the main motivation for experiments with in-house models. The Standard in-house models tended to match humans when tested on natural sources of distortion, largely reproducing the intelligibility variation produced by different types of background noise, reverberation, and lowpass filtering (Supplementary Figures 34-35). These similarities are plausibly explained by the model having been optimized for what we intended to be natural conditions. If humans are also fairly well optimized for such conditions, both humans and models might be performing about as well as is possible in such conditions.

The Standard in-house model also exhibited a considerable amount of human-like generalization to speech distortions that were not present in the model’s training data. For each of local time reversal, noise and tone vocoding, modulation filtering, sine wave speech, and inharmonic speech, the model was robust where humans are robust, and degraded where humans degrade, to about the same extent as humans (Supplementary Figures 34-35). These results are plausibly behavioral signatures of the cues that are used by a system optimized for word recognition. In particular, it doesn’t seem necessary to appeal to biological constraints to account for these phenomena in humans, in that they emerge in a model without such constraints. The industry models were also robust to these distortions, and tended to replicate the overall qualitative effect of the distortions on recognition, but it is difficult to know whether the distortions were present in the industry training data to some extent.

### Residual human-model discrepancies

Even the in-house models that were most similar to humans showed discrepant performance on a fair number of conditions. As of now it is unclear how to understand the remaining gap in human-model similarity. Our hypothesis leading into this work was that the phenotype of human speech recognition is mostly determined by the pressure to communicate effectively in real-world conditions, which over evolution and development might optimize us for real-world speech recognition. Under this hypothesis, discrepancies between task-optimized models and humans might be resolved by optimizing models in conditions even more realistic than those used here.

It also seemed possible that the model architecture might be constraining performance, since there were modest effects of the model architecture (CNN vs. conformer). However, explicitly training either architecture on the discrepant distortions improved performance on those distortions, showing that it is possible for these architectures to perform at or above the level of humans. It remains possible that the architecture in some way limits generalization to these types of distortions. It is also possible that discrepancies reflect the fact that our in-house task of recognizing individual words is impoverished relative to the tasks that shape human speech recognition, and that optimizing for continuous recognition, or for the recognition of sub-word units in addition to words, would elicit models with higher similarity to humans.

The most pronounced discrepancies were evident for bandpass and highpass audio filtering and different types of local silencing. Both of these types of distortion are rare in everyday listening conditions (at least for the extent of filtering that produced variation in performance), and yet humans exhibit some degree of generalization to them, whereas the model did not. One might thus think that these effects provide a useful signature of some property of the human auditory system. However, as of now we have no explanation for why these particular distortions produce discrepancies when many other types of unnatural distortion do not.

### Relation to prior work

Our work builds on a recent tradition of comparisons between human and machine speech recognition. Automatic speech recognition researchers have historically used human performance as a benchmark to aim for, and even with rapid advance in machine speech recognition during the past 15 years, machine systems were found to underperform humans^20,21^. Very recently some industry systems have been found to perform comparably to humans^63^. These comparisons typically involved continuous annotation of speech in quiet conditions, and more recently, speech in noise. As deep learning began to be adopted to build candidate models of human perception, attention shifted to comparing models and humans in other challenging real-world conditions, as human robustness was regarded as a critical trait for models to explain. Kell et al. (2018)^5^ built a model in the style of the in-house models in the present paper (with a fixed cochlear stage followed by a CNN optimized for word recognition) and measured its word recognition performance in different types of noise, finding that the model’s performance across noise conditions was similar to that of humans. A more expansive series of experiments testing robustness to some of the speech manipulations used in the present paper was included in Kell’s PhD thesis and Shook’s Master’s thesis. More recent work found convergence of model and human performance across a large set of natural noise sources^9^.

Another strand of recent work has sought to test industry speech recognition systems on experimental speech manipulations from the human perception literature. Two such attempts using models built roughly ten years ago found significant discrepancies with human behavior^22,23^, the reasons for which remained unclear. More recent work tested state-of-the-art industry systems on a standard clinical assessment of speech recognition in babble noise^24^. The performance of the tested models ranged from on par with humans to slightly worse. Our results are not inconsistent with this work: in the babble noise condition we found the industry systems we tested to perform on par with humans, particularly at lower SNRs. Superhuman performance was most evident in other types of distortion. Overall, we built on these two strands of work by comparing humans and models on a more extensive set of conditions, by using recent industry systems, and by conducting experiments on in-house models to understand the factors contributing to human-model behavioral alignment.

Our work dovetails with evidence that recent language models outperform humans on word prediction tasks^26–29^. Specifically, such models exhibit higher top-1 accuracy than humans when predicting the next word in a sequence^27^. Estimates of human perplexity (a measure of the match between the true and predicted probability distribution of possible next words) are also higher than those of models, suggesting worse predictions^27^. Moreover, comparisons of model surprisal estimates to human reading times suggests that models estimate lower surprisal than humans^26,28^, such that newer, larger language models produce worse predictions of human reading times than some smaller models. Although there are many differences between these language models and speech recognition systems, both types of models use transformer stages and are trained on large amounts of data to predict text. Our work complements these human-model language comparisons by revealing linguistic prediction effects indirectly, showing that superhuman word prediction abilities have a real-world consequence in yielding dramatic improvements in speech recognition in conditions where speech is degraded.

### Limitations

Our tests were limited to a set of words that were both relatively high-frequency and that were of medium length (at least 4 characters). Stimuli were also generated from synthetic speech, and were relatively short (up to 3.5 seconds maximum). Additionally, our model-human comparisons were limited to accuracy, omitting other measures of human behavior like effort or confidence. Moreover, although we assembled a set of distortions that was large compared to previous studies, the distortions we chose were somewhat ad hoc – we tested as many manipulations as we could think of and/or find in prior literature. Overall, our experiments were sufficient to elicit a wide range of performance in humans and models and thus to permit a well-powered comparison, but they inevitably do not provide an exhaustive characterization of speech recognition. It is possible that tests based on recorded conversational speech, and including lower frequency words, would show some residual cases in which humans outperform machine speech recognition. The results nonetheless show that superhuman performance occurs in some regimes across a large set of distortion types.

Although the results suggest that industry speech recognition systems are in some way better than humans at estimating and applying conditional word probabilities, there are several potential explanations for this human-model difference. One possibility is that the models are exhibiting the same effect seen in recent large language models, yielding more accurate word predictions given surrounding words, irrespective of the conditions. A second possibility is that the models are specifically better at estimating word probabilities in conditions of high ambiguity (e.g. when speech is distorted). The ambiguity introduced by distortion plausibly requires probabilities to be estimated for larger numbers of possible word sequences than are required for clean speech. Humans might be limited in how many such sequences can be maintained in memory. Our experiments do not distinguish these two possibilities.

The in-house models were also limited in several respects. The models were trained on an 800-word vocabulary, smaller than that used by humans and word-based industry models. This training task could limit how well the models are likely to match humans. The training data for the in-house models was also restricted to recordings of people reading, which limited the naturalism and variability of the speech training data. The industry models by comparison perform what is arguably a more human-relevant task (word recognition with a human-scale vocabulary), but their training data are not publicly available or described at length, and so it is impractical for external researchers to explore the factors that underlie their performance characteristics. The scale of industry training data, and of the model architectures, stands out as a possible cause of both the superhuman performance we observed, and of the tight alignment with humans once word dependencies were eliminated, but we were not able to test this in a controlled experiment. It is also difficult to know whether the industry models were explicitly trained on some of the distortions in our battery. In at least one case (Whisper-large-v3), the model was reported by the authors to have been trained on filtered speech (the SpecAugment augmentation), but it is plausible that other augmentations were not reported.

### Future directions

None of the models we tested fully account for human speech intelligibility, illustrating the need for some other approach. One path that seems promising is to replicate the scale of industry system training but with a training objective that does not involve speech-specific supervision^64^, augmented by speech-specific training on a scale that is more human-realistic. Such models will additionally be essential for understanding the development of speech perception^65^, and so seem an important goal for independent reasons. It may also be important to build a dataset based on conversational speech to better reflect the distribution of speech that humans encounter, and/or to optimize models for continuous speech recognition (either using sub-word units, or a substantially larger vocabulary).

Apart from having a front-end modeled after the auditory periphery, the model architectures we tested were not biologically constrained. It is an open question as to whether the implementation constraints of biology imprint themselves on speech perception to a substantial extent^66^. Incorporating other features of biological auditory systems – such as recurrence, feedback, or receptive field scaling that resembles the hierarchy of the central auditory system – could provide one test of whether architectures that hew more closely to the brain better account for behavior.

It is also possible that aspects of human speech perception reflect a fundamentally different computational motif. For instance, human perception might be aided by an internal generative model of how speech is produced^67^. Such an internal model could support the ability to imagine speech and might enable generalization to some of the unnatural forms of distortion for which the models tested here underperformed humans. Humans are also known to improve at recognition with repeated exposure to particular types of distorted speech^68,69^. Such effects clearly require a recognition system that is less static than the models we considered here, and potentially a different way of learning from small numbers of examples.

## Methods

### Speech distortion stimuli

#### Speech source clips

Speech excerpts were generated from a vocabulary of 230 common words. Claude Sonnet 4.6 was instructed to generate 4 sentences per vocabulary word, where the target word was either at the end or in the middle of the sentence. Some of the words in the vocabulary did not make sense at the end of a sentence (as judged by the authors) and were excluded from the end-word experiments. As a result, there were 187 x 4 = 732 unique “last word” sentences, 214 x 4 = 856 unique “first word” sentences, and 230 x 4 = 920 unique “middle word” sentences. From these sentences, audio clips were synthesized using the MeloTTS speech synthesizer, and then cloned to 4 voices, 2 male and 2 female, using SeedVC, for a total of 732 x 4 = 2928 last word stimuli, and 3680 middle word stimuli.

For all experiments except the single word experiment, distortions were applied to the entire excerpt, after which a ramp was applied to the beginning and end of the audio. For the single word experiment we determined whether the distortion signal processing would interact poorly with zero-padding. In cases where this was the case, we applied the distortion to the audio signal for the single word without zero-padding, applied the ramp, and then zero-padded to two seconds afterwards.

#### Background noise

We simulated everyday background noise by combining original source clips with five different types of noise: instrumental music, auditory scenes, 8-talker babble, speech-shaped noise, and amplitude-modulated speech-shaped noise. This experiment included some of the background noise conditions from the experiments run by Kell et al. (2018) and Saddler and McDermott (2024). Auditory scene clips were sourced from from the 2013 IEEE AASP Challenge on Detection and Classification of Acoustic Scenes^70^. 8-talker babble was generated by combining 8 speech clips from the Common Voice dataset that were not included in the training. Music was taken from the MUSDB18 dataset^71^. Speech-shaped noise was created by estimating the average amplitude spectrum for the clip’s speaker from all of that speaker’s exemplars in the corpus, and then replacing the Fourier amplitudes of a white noise sample with this average spectrum. We included stationary noise conditions that used the speech-shaped noise as is, and modulated noise conditions that applied amplitude modulation to the speech-shaped noise to match the envelope of a randomly chosen other speech signal. We measured performance for each type of noise at a range of signal-to-noise ratios (SNRs): -9, -6, -3, 0, 3 dB SNR.

#### Background textures

Speech was imposed on 43 different types of synthetic textures at -3 dB SNR. There were 376 unique exemplars of each auditory texture, each generated by performing gradient descent on a unique sample of white noise to match the statistics of each of 43 recorded real-world textures. Synthesis was performed using the McDermott and Simoncelli texture synthesis algorithm^33^.

#### Reverberation

In everyday listening environments, sound reflects off walls and other surfaces. The ear thus receives many delayed and filtered copies of the sound emanating from a source in the world, substantially distorting the original signal. The effects of this reverberation are well described by the convolution of a source signal with the impulse response of the auditory environment for a particular pairing of source and listener locations. To simulate reverberation, we convolved each speech stimulus with synthetically generated impulse responses.

The impulse responses were generated to match the statistics and frequency-dependent properties of real-world impulse responses collected from a survey by Traer and McDermott (2016)^34^. We generated impulse responses with a direct-to-reverberant ratio (DRR) of either 20 or 50 dB, and an RT60 of 200, 400, 800, and 1600 ms. Synthetic impulse responses were generated by imposing exponential decay on the subbands of a noise sample, with decay constants obtained by interpolating the decay rates measured in the Traer and McDermott real-world impulse response. The interpolation yielded decay rates for each RT60 value. The DRR was set by scaling the decaying noise (the “tail” of the impulse response) relative to an initial impulse at the start of the impulse response. Impulse responses were synthesized using a Python script written by James Traer.

To add realism and obtain a wider range of human and model performance, in addition to applying reverberation to clean speech, we also applied it to speech in babble (simulating a conversation in a crowded reverberant room). Each babble clip was generated by combining 8 clips from the WSJ/TIMIT dataset. We added the speech and babble at an SNR of 0 dB, and then applied reverberation.

#### Whispered speech

Simulated whispered speech was generated using STRAIGHT as described in more detail below for the generation of inharmonic speech, replacing the harmonic source with high-pass filtered Gaussian noise to mimic the unvoiced breath in whispering^35^. The speech signal was decomposed into its time-varying spectral envelope, periodic excitation, and aperiodic excitation and was then resynthesized while omitting the periodic excitation, filtering the aperiodic excitation to make it more closely resemble whispering.

#### Audio filtering

Filters were generated with the scipy function scipy.signal.firwin with num_taps set to 2001. The filter was applied using scipy.lfilter. Lowpass speech was generated with octave-spaced filter cutoffs ranging from 400 Hz to 3200 Hz. Highpass speech was generated with octave-spaced filter cutoffs ranging from 400 Hz to 6400 Hz. Bandpass filtered speech was generated with a center frequency of 1500 Hz and filter widths of 5, 10, 20, 30, and 40 semitones.

#### Time compression and dilation

We temporally compressed or dilated speech excerpts by time-stretching (as to preserve the f0 contour and vocal tract filtering) using Praat^72^. We measured human and network performance on seven different speeds, ranging from twice as fast to twice as slow: 0.5x, 0.75x, 0.875x, 1.0x, 1.25x, 1.5x, 2.0x.

#### Inharmonic speech

To generate inharmonic speech, we used a previously published method^35^. The source and filter were first estimated from a natural speech source signal. The source was then modified to be inharmonic by independently jittering the frequency of each constituent harmonic, and was then combined with the original filter. As in the original publication (Popham et al., 2018)^35^, we performed this manipulation using a variant of the STRAIGHT analysis algorithm^73^ that models periodic excitation as the sum of frequency-modulated sinusoids^74^. The frequencies of these sinusoids were jittered by a random proportion of the fundamental frequency (f0), with the proportion drawn uniformly from between *-p* and *p*, where *p* defined the condition. Jitters were sampled repeatedly until they satisfied the constraint that adjacent frequencies remained at least 30 Hz apart, to avoid salient beating between adjacent harmonics that might differ across different degrees of inharmonicity. The resulting sinusoids were then recombined with the aperiodic component of the speech excitation, and filtered to impose the original spectral envelope. The degree of inharmonicity was varied across conditions, with *p* set equal to 0%, 10%, 30%, or 50%.

#### Sine-wave speech

Sine-wave speech^30^ was generated by measuring the time-varying trajectories of the first four formants of a speech signal using an 8^th^ order linear predictive coding model, generating frequency-modulated sinusoids that followed these trajectories (one for each formant), and summing these frequency-modulated sinusoids. Stimuli were generated using MATLAB code written by Dan Ellis.

#### Modulation filtering

To generate spectrally and temporally modulation-filtered speech, we followed the method described in Elliott and Theunissen (2009)^32^, using code provided by the authors. First, the signal was transformed into a spectrogram. Second, the 2D Fourier transform of the spectrogram was taken to obtain the modulation power spectrum, and the amplitude of every Fourier component above the modulation filter cutoff (3, 6, 12, or 24 Hz for temporal modulations; 0.5, 1, 2, 4, or 8 cycles/kHz for spectral modulations) was set to 0. Third, the inverse 2D Fourier transform was taken to yield a spectrogram, and finally, an iterative spectrogram inversion process was applied to transform this back into an audio signal^75^.

#### Noise vocoding

To generate noise-vocoded speech, the speech signal is filtered into frequency subbands, the amplitude envelopes of which are extracted and applied to a noise signal. The noise-vocoded signal is generated by summing the new subbands together. Noise vocoding substantially alters the spectral structure of the source signal, but largely preserves the temporal envelope cues, and was originally motivated as an attempt to simulate the kind of signal processing that occurs in a cochlear implant^31^.

In our noise-vocoding algorithm, waveforms were first passed through a bank of 320 bandpass filters. Filters were zero phase and were the positive portion of a single period of a cosine function, with center frequencies ranging from 20 Hz to 10 kHz, and adjacent filters overlapping by 50% so that the filters perfectly tiled the frequency spectrum^33^. Filters were evenly spaced on an Equivalent Rectangular Bandwidth (ERB) scale^76^. These (very narrow) subbands were summed across frequency in various combinations to produce the desired number of channels (i.e., 1, 2, 4, 8, 16, and 32 bands). For instance, to generate a single band, all of the subbands were summed, whereas to generate 4 bands adjacent sets of 80 filters were summed. The envelopes of these summed subbands were extracted by applying the Hilbert transform and taking the modulus of the analytic signal. These envelopes were imposed on the subbands of a Gaussian noise signal by multiplication with the noise subbands, after dividing the noise subbands by their original envelope. The resulting subbands were refiltered with a filterbank of either 1, 2, 4, 8, 16, or 32 filters (obtained by summing together subsets of the transfer functions of the filters used to obtain the original set of narrow subbands) and summed to generate the noise vocoded speech signal.

Our human participants’ performance with four bands is lower than that reported in the original paper on noise-vocoded speech^31^, likely because participants in the original experiments received 8-10 hours of training and received feedback, whereas our participants received neither training nor feedback in order to replicate the network’s testing conditions (the network was not trained on noise vocoded speech).

#### Tone vocoding

Speech clips were filtered into 32 frequency subbands using a bank of bandpass filters intended to mimic the frequency selectivity of the human cochlea. Filter center frequencies ranged from 100 to 10000 Hz, equally spaced on an ERB_N_ scale^76^. Subbands above the cutoff channel were tone-vocoded by point-wise multiplying the envelope of the subband with a pure tone with a frequency equal to the center frequency of the subband. These tone-vocoded subbands were then bandpass filtered again and summed with the other subbands to produce a modified speech signal. Stimuli were generated by tone-vocoding subbands above cutoffs of 0, 8, 16, 24, and 32 subbands.

Tone-vocoded stimuli were imposed on either stationary or amplitude-modulated speech-shaped noise^77^. Speech-shaped noise backgrounds were made individually for each speech signal by generating flat-spectrum noise with a long-term power spectrum matching that of the speech signal. Modulated noise was generated by imposing sine-wave modulation on speech-shaped noise at a modulation rate of 8 Hz, with a peak-to-valley ratio of 30 dB.

Experimental results were converted into estimates of a 50% speech-reception threshold so that the benefit of modulated vs. stationary noise for each cutoff channel could be quantitatively compared. For each cutoff, background noise type and SNR, we calculated the average proportion correct across all human participants or model architectures which we grouped into psychometric functions for each cutoff and noise type. We fit this data with a cumulative gaussian distribution function to estimate the SNR at which performance was 50%. Following the original publication^77^, the cost of tone-vocoding was summarized by subtracting the speech-reception threshold at a cutoff channel of 0 from the threshold of each higher cutoff value. Intuitively, this measures how much better humans or models are at recognizing speech when fewer channels are tone-vocoded and compares this effect between modulated and stationary noise.

#### Chimeras

A chimeric audio signal is created by combining the envelope of one sound with the fine structure of another^36^. Each sound is filtered into a given number of subbands, and each filtered subband is decomposed into its envelope and fine structure using the Hilbert transform. Two types of chimeric signals were used here: speech-speech chimeras and speech-noise chimeras. In a speech-speech chimera, the two source sounds are different clips of speech, so that the resulting chimera has conflicting speech information. When testing models/humans, a response was marked as correct for the fine structure condition and incorrect for the envelope condition if the scored word of the fine structure source sound was predicted, and correct for envelope and incorrect for fine structure if the scored word of the envelope source sound was predicted (and incorrect for both otherwise). In a speech-noise chimera, the envelope of noise is imposed on the fine structure of speech. The complementary chimera in which the envelope of speech is imposed on the fine structure of noise was not included because it is similar to noise-vocoded speech, which we tested in a separate manipulation. Speech-noise and speech-speech chimeras were generated with 1, 2, 4, 8, and 16 subbands. Because each speech-speech condition was scored both for the fine structure sound source and the envelope sound source, the five speech-speech conditions yielded 10 points on the results graphs (yielding 277 points in total from the 272 total conditions).

#### Clipping

There were two types of clipped speech; peak-clipped speech and center-clipped speech. Given a proportion between 0 and 1, a threshold amplitude value is chosen such that the chosen proportion of samples in the audio clip have values less than the threshold (e.g. if the chosen proportion is 1, the threshold would be equal to the maximum absolute waveform peak in the signal, and if the chosen proportion is 0, the threshold would be equal to 0). Peak-clipped speech was generated by setting every sample in a speech signal above that threshold to the threshold amplitude multiplied by the sign of the sample. If the proportion is equal to 1, the signal is unchanged, and if the proportion is equal to 0, the signal becomes the sign signal. Center-clipped speech was generated by setting every sample in a speech signal below that threshold to 0. If the proportion is equal to 1, the signal becomes all zeros^37^, and if the proportion is equal to 0, the signal is unchanged. Peak and center clipped speech was generated with thresholds of 0, 0.25, 0.5, 0.75, 0.9, and 0.98.

#### Local time-reversal

Time-reversed speech was generated by dividing a source speech signal into segments of a given length (10, 20, 30, 40, 50, 100, 150, or 200 milliseconds) and temporally reversing each segment of the waveform in place^38^. As in the original work, no smoothing was performed at boundaries of these segments.

#### Local shuffling

Shuffled speech was generated by splitting a speech signal into segments of a particular length (ranging from 1/8 ms to 4 ms by factors of 2), and then randomly reordering the samples within each segment^39^. This manipulation preserves some of the global structure of the signal while destroying the local structure.

#### Local silencing

To generate silenced speech, speech was divided into segments of a given length (1, 3, 10, 30, or 100 milliseconds), and a given proportion (1/4, 1/2, or 3/4) of the segment was silenced^40^.

#### Local masking

Masked speech was generated similarly to silenced speech, except instead of silencing a proportion of each segment, a proportion (always 1/2) of each segment was replaced with Gaussian noise such that the signal-to-noise ratio of each segment was -18, -9, 0, or 9 dB^40^.

#### Repackaging

Repackaging is a variant of local silencing generated by first time-compressing speech by a factor of 2, and then inserting silence (0, 10, 20, 40, 80, or 160 milliseconds in duration) every 40 milliseconds^41^. In the condition where 40 milliseconds of silence is added, the speech information in the resulting signal is presented at the same overall rate as the speech information in the original signal. Previous literature has suggested that this condition would be the most intelligible to humans^41^.

### Human psychophysics

#### Ethics statement

All participants provided informed consent and the Massachusetts Institute of Technology Committee on the Use of Humans as Experimental Subjects (COUHES) approved all experiments.

#### Overview

Human experiments were conducted online via the Prolific platform. At the beginning of the experiment, participants were asked to set their sound volume to a comfortable level using a peak-normalized calibration noise with rms level 0.3. They then performed a headphone check to test whether they were wearing headphones^78^. The stimuli were set to rms level 0.03. This was the level needed to avoid clipping, but was also judged by the experimenters to be comfortably audible when the volume was adjusted using the calibration noise. Pilot experiments indicated that human performance was robust to the exact sound presentation level. Many previous studies conducted in our laboratory have consistently demonstrated that online data can match the quality of data collected in traditional laboratory settings^5,79–85^ provided measures are taken to ensure standardized sound presentation, encourage participant compliance, and promote active engagement in the tasks.

For each main experiment, participants heard excerpts that contained a target word drawn from a vocabulary of 230 common words. The stimuli were generated as described above. 272 (or 196 in the case of Experiment 6, the full-sentence experiment) types of distortions were then applied to the stimuli. Participants were randomly assigned to distorted stimuli as described below.

For each experiment, 400 participants were run who passed the inclusion criterion (based on catch trials, described below). Each participant was assigned to half the distortion conditions, so that each pair of participants received the full set of distortions, and so each of the distortion conditions was presented on a total of 200 trials. Each participant heard each word no more than once, and the words were assigned such participant 1 heard words 1 through D/2, participant 2 heard words D/2+1 through D mod N, etc. where D is the number of distortions (272 or 196 depending on the experiment) and N is the number of total words (187, 230, or 214 depending on the experiment). In this way, each target word was presented the same number of times across participants, with the exception that some words were presented 1 trial fewer in total.

Catch trials were distributed throughout the experiment with the intention of assessing whether participants were paying attention and engaged in the task. On catch trials, humans were presented with a speech clip in which a single word was spoken very clearly. Participants who got more than 2 out of 15 catch trials wrong were excluded from the analysis, and then those conditions were rerun with new participants. Participants were compensated at a rate of approximately $12.00 per hour.

#### Sample size

We determined the target sample size using a power analysis based on several pilot experiments. The power analysis estimated the reliability of the pattern of intelligibility across distortions as a function of sample size. The pilot experiments differed from the actual experiments in presenting different speech material and subsets of the full set of distortions (described in more detail below in the Pilot Experiments section). Participants heard each target word once, randomly assigned to the subset of distortions that were tested (such that each distortion was presented multiple times; each distortion was presented about 10 times per participant). For the power analysis, we attempted to simulate the planned design of the main experiment by pooling across experiments while subsampling participants for each experiment, so that all distortions were represented. We could then calculate the average proportion correct for each distortion condition from this pooled data. We computed the split-half reliability of the resulting pattern of intelligibility across all distortions for 1000 random splits of participants of size N, with N ranging from 1 to 5 (this upper limit was half of the smallest sample size collected in the pilot experiments). We extrapolated from these results that a sample size of 20 participants (equivalent to 200 trials for each distortion, given that each participant heard each distortion 10 times on average) would be sufficient to reach a split-half reliability of 0.95. Because the planned design of the main experiment involved presenting each distortion once per pair of participants, the target of 200 trials per distortion translated to a target sample size of 400 participants.

#### Experiment 1 – coherent sentence excerpts (last word)

We collected data from 458 participants who passed the headphone check, 58 of which were excluded due to catch trials, for a total of 400 participants (182 female), mean age: 38, SD: 1 Stimuli were generated from the “last word” set comprised of 187 unique vocabulary words, for 187 x 4 sentences per word x 4 cloned voices = 2992 different total stimuli. Audio clips were cropped to 2 seconds. Participants were asked to transcribe the last word that they heard in the excerpt.

#### Experiment 2 – single words

We collected data from 451 participants who passed the headphone check, 51 of which were excluded due to catch trials, for a total of 400 participants (205 female), mean age: 42, SD: 13. For each of the 230 words, we synthesized 4 stimuli, one for each of the 4 cloned voices. 10 of the synthesized words were judged by the experimenters to have been pronounced abnormally, and were excluded, for a total of 220 x 4 cloned voices = 880 stimuli. Participants were asked to transcribe the word that they heard.

#### Experiment 3 – random word sequences (last word)

We collected data from 476 participants who passed the headphone check, 76 of which were excluded due to catch trials, for a total of 400 participants (191 female), mean age: 40, SD: 12. Stimuli were generated from the “last word” sentences, with words pseudo-randomized across stimuli as described below, synthesized, and then cropped to 2 seconds in length. Participants were asked to transcribe the last word that they heard in the excerpt.

Randomization algorithm: All of the non-target words were taken and put into a list W, sorted by how frequently they occurred (e.g. the first X words in the list were “the” where “the” occurred X times overall in the sentences). Then, for L equal to 1 until the length of the longest sentence in words, this process was repeated. The number of sentences with number of words greater than L was tallied for a total number N. The first N words were taken from the list W and randomly shuffled. Then one word each was assigned to each of the qualifying sentences. In this way, all of the words in W were assigned to the sentences such that each sentence has a mix of high, mid, and low frequency words. Then, the non-target words in each sentence were pseudo-randomly shuffled so that the same word did not appear twice in a row. This resulted in a set of pseudo-random word sequences where each word sequence ended with a target word.

#### Experiment 4 – coherent sentence excerpts (first word)

We collected data from 495 participants who passed the headphone check, 95 of which were excluded due to catch trials, for a total of 400 participants (218 female), mean age: 42, SD: 12. Stimuli were generated from the “first word” set comprised of 214 unique vocabulary words, for 214 x 4 sentences per word x 4 cloned voices = 3424 different total stimuli. Audio clips were cropped to 2 seconds. Participants were asked to transcribe the first word that they heard in the excerpt.

#### Experiment 5 – coherent sentence excerpts (middle word)

We collected data from 473 participants who passed the headphone check, 73 of which were excluded due to catch trials, for a total of 400 participants (190 female), mean age: 38, SD: 12. Stimuli were generated from the “middle word” sentences comprised of 230 unique vocabulary words, for 230 x 4 sentences per word x 4 cloned voices = 3680 total stimuli. So that the in-house models could be evaluated on these stimuli as well, we restricted the stimuli so that the target word overlapped the 1-second mark. For some of the stimuli, the target word appeared too early or late (these stimuli were excluded), leaving 3343 total stimuli. The stimuli were then randomly cropped to 2 seconds such that the target word overlapped the 1-second mark. Since it is difficult to determine from listening to an audio clip which word overlaps the 1-second mark, participants were asked to transcribe the entire excerpt instead of just the target word, with scoring restricted to the middle word.

#### Experiment 1b – coherent sentence excerpts (last word)

We collected data from 445 participants who passed the headphone check, 45 of which were excluded due to catch trials, for a total of 400 participants (205 female), mean age: 38, SD: 12. The stimuli were identical to those of Experiment 1. The only difference from Experiment 1 is that participants transcribed the entire excerpt instead of just the last word.

#### Experiment 6 – full sentences (middle word)

We collected data from 473 participants who passed the headphone check, 73 of which were excluded due to catch trials, for a total of 400 participants (210 female), mean age: 41, SD: 12. Stimuli were synthesized from the sentences generated for the middle word experiment (Experiment 5), with sentences less than 2 seconds or more than 3.5 seconds in length excluded, for a total of 3632 stimuli. Some of the distortions were implemented in a way that relied on stimuli being exactly 2 seconds in length and were omitted from this experiment (omitted conditions: 43 background textures, 5 background noise conditions, and reverberation in 8-speaker babble), yielding 196 total distortion conditions. Participants were asked to transcribe the entire sentence.

#### Pilot experiments

Pilot experiments used to estimate the sample size are shown in Supplementary Figures 1-2. Pilot experiment stimuli were excerpts of recordings from the Wall Street Journal and TIMIT corpora for which the middle word (overlapping the 1-second mark) was taken from the 230-word vocabulary described above. The pilot experiments differed from the main experiments in a few other ways. First, categories of distortions were assigned to different experiments with different sets of participants. The categories were audio filtering (lowpass, highpass, bandpass), modulation filtering (lowpass temporal and spectral modulation filtering), repackaging/silencing (repackaging and local silencing), masking (local masking), chimeras (speech-speech and speech-noise chimeras), clipping/shuffling (peak and center clipping and shuffling), spectral manipulations (sine wave speech, whispering, inharmonic speech, noise vocoding), reverberation (clean reverberation and reverberation in babble), time-compression/dilation, local time reversal, background noise (5 types of background noise x 5 SNRs), 43 textures, and tone vocoding. Since there were different numbers of conditions within each category, some conditions were heard more often than others. For each category, the conditions were divided among 230 trials so that each participant heard each word once. Tone vocoding was excluded from Supplementary Figures 1 and 2 due to humans and models being tested on different SNRs. Second, the experiment was analogous to Experiment 5 (the middle word experiment), except that humans only typed the middle word as opposed to all words in the excerpt. Third, humans saw the vocabulary on the screen and were only able to enter words from the vocabulary.

### Model evaluation and psychophysics

#### Industry speech recognition models

Industry models were tested on the same stimuli used in human experiments. The industry models performed continuous speech recognition on the speech clips. If the target word was found in their transcription, the model trial was marked as correct. Otherwise, it was marked as incorrect. In the case of the full sentence experiment, models were graded in the same way that humans were. Parakeet had an issue where it produced empty strings as output for some of the stimuli for the experiments that presented sentence excerpts where the beginning of the sentence was cut off (Experiment 1, 1b, 3, and 5). For those experiments, we excluded the source clips where Parakeet outputted empty strings on clean speech from Parakeet’s evaluation. Results where those source clips were also excluded from human evaluation can be found in Supplementary Figure 39.

#### In-house models

In-house models were tested on the same stimuli used in Experiment 5 since they were trained to perform middle word prediction. The word class with the highest probability out of the model’s 800-word vocabulary was taken as the model’s choice.

### Statistics

#### Aggregate measures of human-model similarity

We captured qualitative similarity between human and model performance using Pearson correlations. This measure is high if the relative performance differences between conditions is similar between humans and models, regardless of whether absolute performance is similar. Correlation was computed from the model and human accuracies across the 277 experimental conditions (pooled across experiments). We note that the experiments presented 272 stimulus conditions in total, but this translated to 277 experimental conditions for analysis because the speech-speech chimera conditions were scored twice, once for the fine-structure source and once for the envelope source.

#### Aggregate measure of superhumanness

We captured the amount of superhumanness with a metric Δ, which was the average difference between model and human proportion correct across the 277 experimental conditions. We computed 95% confidence intervals on Δ by bootstrapping 1000 times across the 277 conditions.

#### Statistical tests of differences between models

We used a two-sided Wilcoxon signed-rank test to determine statistical differences in performance in four settings: a) between a model and humans on the same experiment, b) between a model in one experiment vs. the same model in another experiment, c) between humans in one experiment vs. humans in another experiment, and d) between one model and another model on the same experiment. In each case these tests operated on arrays of performance (proportion correct) for each of the 277 conditions.

To determine confidence intervals for the model-human performance difference metric (s), we bootstrapped over the array of model-human differences and took the 95% confidence interval lower and upper bound.

To determine confidence intervals for r^2, we bootstrapped over the 277 conditions and took the 95% confidence interval.

To determine whether a model’s performance in one experiment was more superhuman than the model’s performance in another experiment, we bootstrapped over the 277 conditions. For each bootstrap sample we calculated the average model-human performance difference for the first model (Δ1) and the second model (Δ2). We estimated the p value for the difference of these differences as the proportion of sample for which Δ1-Δ2 were below zero.

We determined whether one model had a better match to humans (as measured by Pearson correlation) than another model in a similar way. We let Δ*r* be the difference between the Pearson correlation of model 1 with humans and the Pearson correlation of model 2 with humans. We then bootstrapped over the 277 conditions to get an array of differences Δ*r*’ between Pearson correlations for the bootstrap samples. We estimated the p value for Δ*r* as the proportion of sample for which Δ*r*’ was below zero.

#### Surprisal analysis

Distil-GPT2^86^ was used for the sequential surprisal analysis while BERT-base-uncased^49^ was used for the acausal surprisal model. Sequential and acausal surprisal was computed for the target word (the target word of the corresponding experiment) of the transcripts used for the stimuli for Experiments 1, 3, 4, and 5. Surprisal was only computed for the transcripts that contained target words common to all of these 4 experiments. For sequential surprisal, surprisal of each transcript was computed using a surprisal function built into Distil-GPT2, which yielded surprisal values for each word. For acausal surprisal, word surprisal was computed from each of the tokens within the target word by repeating the following process for each token. First, the token was masked. Then the log probabilities for each possible token in that position given the surrounding context were computed. Surprisal for the token was computed as the negative log base 2 probability of that token. Surprisal for the entire word was computed as the sum of the surprisals of the tokens. A two-sided Wilcoxon test was used to compute statistical significance of differences in surprisal between experiments.

### Model tasks

#### Industry model speech recognition task

Industry models were presented with speech audio and produced word transcripts. In cases where humans were evaluated based on a target word (one of the words contained in the speech stimulus), the models were marked correct based on whether the target word was present in their transcription.

#### In-house model word recognition task

All in-house models were evaluated on the task of recognizing the middle word of a 2-second speech excerpt. This task was chosen as it had been used in prior work^5,9,11,56,57^. The middle word was defined as the word overlapping the 1-second mark of the audio clip. The target words were drawn from a set of 800 words, chosen as those for which we could create a relatively balanced training set for the in-house models (i.e., in which each word class had roughly the same number of training examples). Given the goal of balancing training in this way, Zipf’s law places a limit on how many words can be included. We settled on 800 as a number that was large enough to form a nontrivial recognition task (i.e., of similar order of magnitude to standard object recognition tasks) but that allowed a balanced dataset to be constructed from speech corpora that were publicly available at the time of the work. We note that all experiments tested models and humans on a subset of 230 of these 800 words.

### In-house model architecture and training

#### Cochlear model

The first stage of the model was intended to approximate the human auditory periphery. This stage was identical to the simplified cochlear model used in an earlier publication from our lab^9^, and consisted of 3 operations: filtering with a bank of bandpass filters with tuning modeled on the human cochlea, half-wave rectification, and low-pass filtering to simulate the upper limit of phase locking.

The input to the model was a 20 kHz audio signal. This signal was passed through a finite impulse response approximation to a gammatone cochlear filter bank (50 filters, with center frequencies ranging from 60 to 8000 Hz, evenly spaced on an ERB_N_ scale^76^). Gammatone impulse responses were truncated at 50 ms. After half-wave rectification, a low-pass filter was applied (with a cutoff of 3000 Hz). The resulting signals were downsampled to 10 kHz. Pointwise sigmoid functions approximating the rate-level functions of high, medium, and low spontaneous rate fibers were then applied to generate simulated firing rates. These functions had spontaneous rates of 70, 4, and 0.1 spikes/s, thresholds of 0, 12, and 28 dB SPL, and 95% dynamic ranges of 20, 40, and 80 dB.

#### Model architecture (CNN)

The CNN model architectures included eight types of operations, each applied in a dedicated “layer”:

- Convolution (kernel size (x, y) number of filters z)
- ReLU activation
- Hanning pool (pool size (x, y) stride (w, z))
- Layer normalization
- Fully-connected flatten layer
- Fully-connected dense layer (512 units)
- Dropout layer (rate of 50%)
- Fully-connected classification layer (801 units for the 800 word classes and one null class)

A “block” of the network was composed of the first four layer types in order (convolution, ReLU, Hanning pooling and layer normalization) with parameters that varied depending on the depth. Each model consisted of the initial cochlear stage, and then 7-8 blocks, depending on the architecture, followed by a fully connected flatten layer, a fully connected dense layer with 512 units, a final ReLU and normalization layer, a dropout layer, and finally the classification layer (801 classes, representing the 800-word vocabulary plus a null class). See Supplementary Fig. 40 for full specification of the 9 architectures. The Small-dataset model was only trained on CNN architecture 1.

#### Model architecture (Conformer)

All of the conformer models used the same architecture^53^. They had 8 blocks total, with the first 4 blocks of the model being convolutional, while the last 4 blocks were conformer layers. The 4 convolutional blocks were composed of a convolutional layer, SiLU activation, average pooling, and batch normalization. The conformer layers were convolution-augmented transformer layers, each with 4 attention heads, a kernel size of 31, and feedforward dimension of 256. The final layers of the model were the same as for the CNN architecture, except SiLU was used instead of ReLU. The number of total parameters in this model was approximately 303M (See Supplementary Fig. 40 for specification of architecture).

#### Model architecture (parameter-matched CNN)

The parameter-matched CNN was a variant of the standard CNN architecture. To approximately match the number of parameters in the Conformer, several blocks in the second half of the CNN were widened (i.e., the number of channels was increased). This resulted in the fully connected layer having 2048 units instead of 512. In addition, the convolutional kernel of the final block was changed from 8x8 to 12x12. The number of total parameters in this model was approximately 313M (See Supplementary Fig. 40 for specification of architecture).

#### Model optimization

All in-house models were trained to 250K steps, which was equivalent to 10 epochs (an amount that we determined was sufficient for model performance to plateau). The models were optimized using the Adam optimizer with a learning rate of 1e-5. The loss was computed with sparse categorical cross-entropy loss. CNN and Conformer models were trained across 4 A100 GPUs, with a per-GPU batch size of 32. CNNs took ∼27 hours to train across the four GPUs. Validation performance was measured every 2500 steps. If the validation loss improved, then it was saved as a checkpoint. Otherwise, it was discarded. After every validation step, the model continued training from its last saved checkpoint (i.e. the checkpoint with the best validation loss up to that point). This was done to conserve disk space and prevent overfitting.

### In-house model training datasets

#### Overview

In-house models were trained to identify the middle word (i.e. the word overlapping the 1-second mark) in a 2-second speech clip from an 800-word vocabulary. For all in-house models, speech excerpts used for training were taken from the Common Voice dataset. Since the Common Voice dataset has many more male than female talkers, to balance the dataset across talkers, the 800-word vocabulary was chosen by finding the unique words spoken by the greatest number of female talkers. For models whose training sets included background noise, background noises were taken from the AudioSet dataset (a dataset of YouTube soundtracks that includes various naturalistic background sounds)^55^ and combined with foregrounds during training such that the same speech clip is presented with different noises during training (SNRs ranged from -10 to 10 dB and also included a +inf condition without background noise). Background AudioSet clips were excluded that might interfere with the word (i.e. those that had labels that meant they contained speech). Additionally, clips that were only labeled ‘Music’, and clips that had sampling rates less than 40 kHz and durations less than 9 seconds were excluded.

We trained several different versions of in-house models, each with different training data, in order to investigate the effect of the training diet on human-model similarity.

#### Standard dataset (CNN and Conformer)

Around 3.2 million 2-second speech excerpts were sampled from the set of Common Voice clips that included one of the words in the 800-word model vocabulary (4000 excerpts per word), subject to the constraint that there were as many unique speakers as possible. To do this, the speakers in the set of clips were weighted inversely by how many times they occurred in the dataset. Then, 4000 clips per word were sampled without replacement until it was impossible to do so, at which point they were sampled with replacement. This resulted in 3.2 million clips total, although there were only 2.5 million unique exemplars. For the validation set used to monitor performance during training, 160K 2-second clips were sampled in the same way (200 examples per word). Both the training set and validation set used backgrounds from AudioSet.

#### 100 talkers dataset (CNN and Conformer)

To reduce the variability of the talkers while also having as much training data as possible, only speech clips from the most frequently occurring 50 male and 50 female talkers were selected. This resulted in around 500K unique speech clips. Clips were sampled with replacement as needed to reach 3.2 million foreground training clips and 160K foreground validation clips. Background noises were the same as those in the Standard dataset.

#### Small dataset (same size as 100 talkers, but with many voices, CNN only)

To test whether the worse human-model match in the model trained on the 100-talkers dataset was due to the decreased number of unique foreground clips or the reduced speaker variability, we made an alternative dataset in which the number of unique talkers was as large as possible, but the number of unique foreground clips was approximately equal to that in the 100-talkers dataset. The number of unique foregrounds in the 100-talkers dataset was approximately 1/5 of the number for the standard model dataset. So, for the small dataset, the same sampling algorithm as for the standard model was performed, but with only up to 800 instances of each word. Then, from this set, clips were sampled with replacement up to 4000 instances of each word. The validation set was sampled in the same way as in the Standard dataset.

#### No backgrounds dataset (CNN and Conformer)

This dataset was identical to the Standard dataset, except that background noise was omitted.

#### Augmented dataset (CNN and Conformer)

To further explore the discrepancies between the in-house neural networks and humans, an additional model was trained using the same speech (from Common Voice) and background noise (from AudioSet) excerpts, but with augmentations applied to the speech. These augmentations were chosen based on the experimental conditions that produced the most pronounced discrepancies between the Standard models and humans. The question was whether the model architecture was capable of yielding human-like performance on these conditions if it was explicitly trained on them. The full set of augmentations for this training condition are as follows:

Bandpass filtering was applied to 10% of the dataset. We used low cutoffs log-uniformly sampled between 40 and 1200 Hz, and high cutoffs log-uniformly sampled between 1600 and 10000 Hz. We chose these cutoffs to encompass the range of bandpass filtering in our experiment.

Highpass filtering was applied to 7.5% of the dataset. We used filter cutoffs log-uniformly sampled between 40 and 6400 Hz. For all audio filtering manipulations, the order of the filter was chosen uniformly at random from the set [1, 2, 3, 4].

Time-compression and dilation were applied to 10% of the dataset. The amount of time-compression was uniformly sampled between 0.5x and 2x speed. We chose these limits as they were the limits in the compression and dilation experiment.

Local silencing was applied to 5% of the dataset. The reciprocal of the rate of silencing was uniformly sampled on a logarithmic scale between 1/1000 and 1/10 Hz^-1^, and the fraction of the clip being silenced was uniformly sampled between 0.25 and 0.75.

## Code and data availability

All code and data will be made available in a project repository upon publication.

## Author contributions

A.M. designed and ran the human experiments, trained the in-house CNN models, tested all in-house models, made the figures, and wrote the initial draft of the paper. G.E. helped design the experiments, synthesized the speech stimuli, and ran the industry models in the experiments.

E.S. and A.K. designed some of the experiments and ran pilot versions of these experiments on earlier versions of in-house models. M.R.S. created the architectures for the standard CNN and conformer models, trained the conformer models, and generated the training data for the in-house models. J.H.M. supervised the research, raised the funding for the research, and wrote the initial draft of the paper. All authors edited the paper and contributed to project planning.

## Acknowledgements

Research supported by NSF grant BCS-1634050 and National Institutes of Health grants R01 DC014739 and R01 DC017970. The authors thank the McDermott lab for helpful feedback on the manuscript.

## Supplementary figures

**Supplementary Figure 1:**
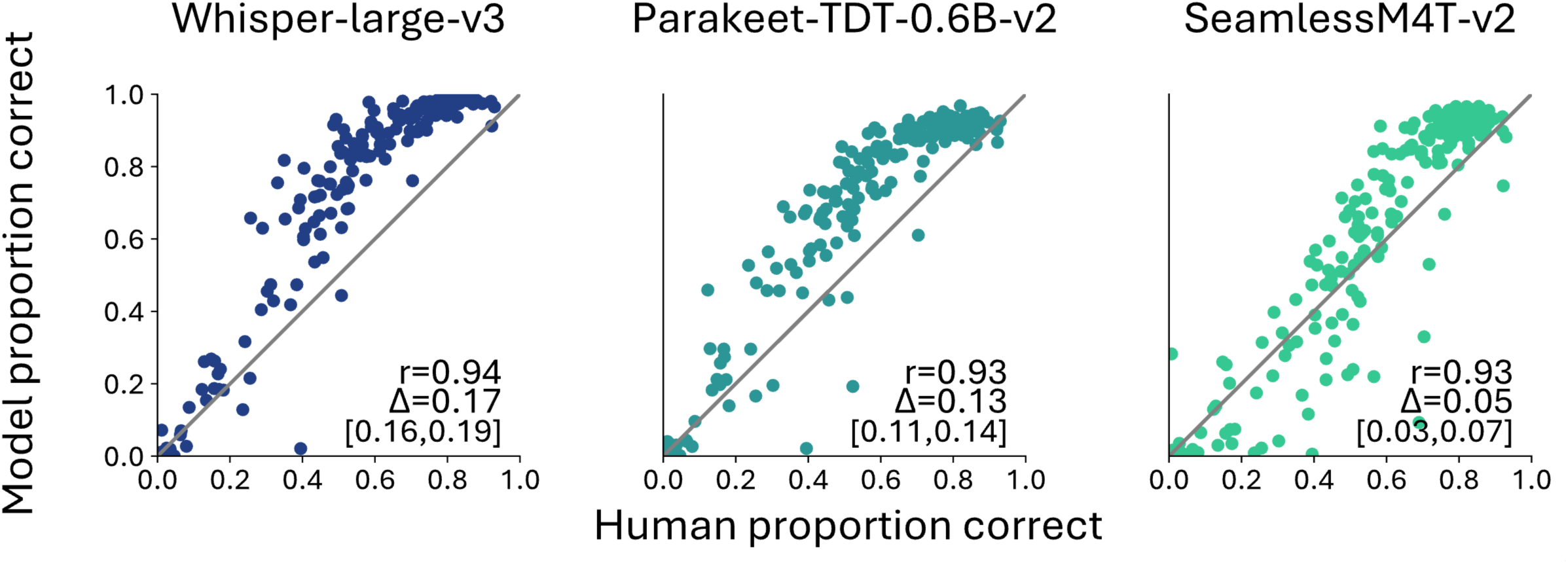
Scatter plots of performance for pilot experiment with recorded speech. Stimuli were generated from excerpts from WSJ and TIMIT speech corpora (see Methods).

**Supplementary Figure 2:**
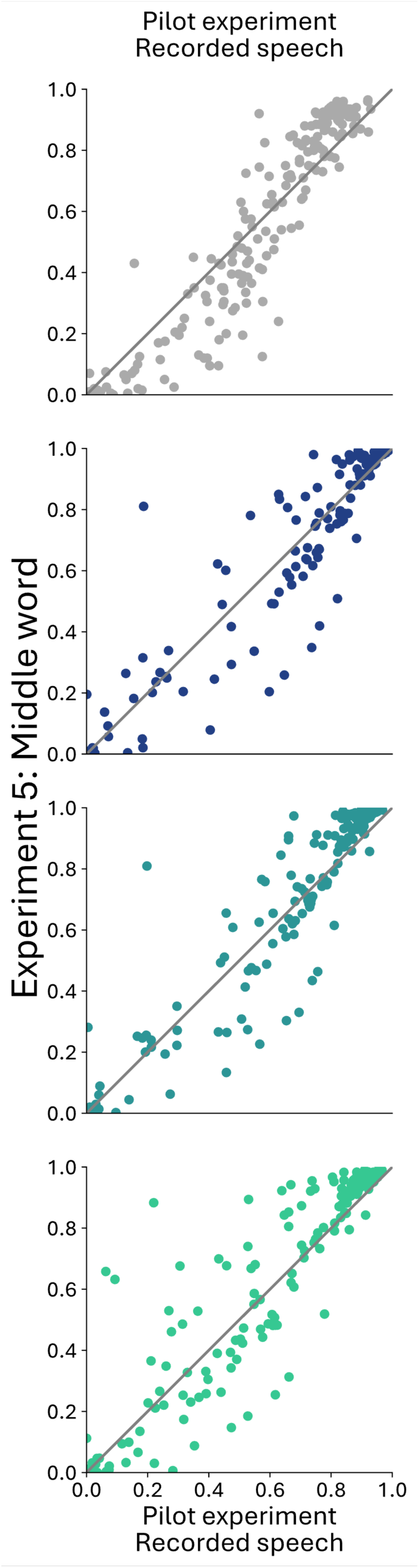
Comparison of performance on stimuli generated from recorded speech (Pilot experiment, see Methods) vs. stimuli generated from synthetic speech (Experiment 5) for humans and models.

**Supplementary Figure 3:**
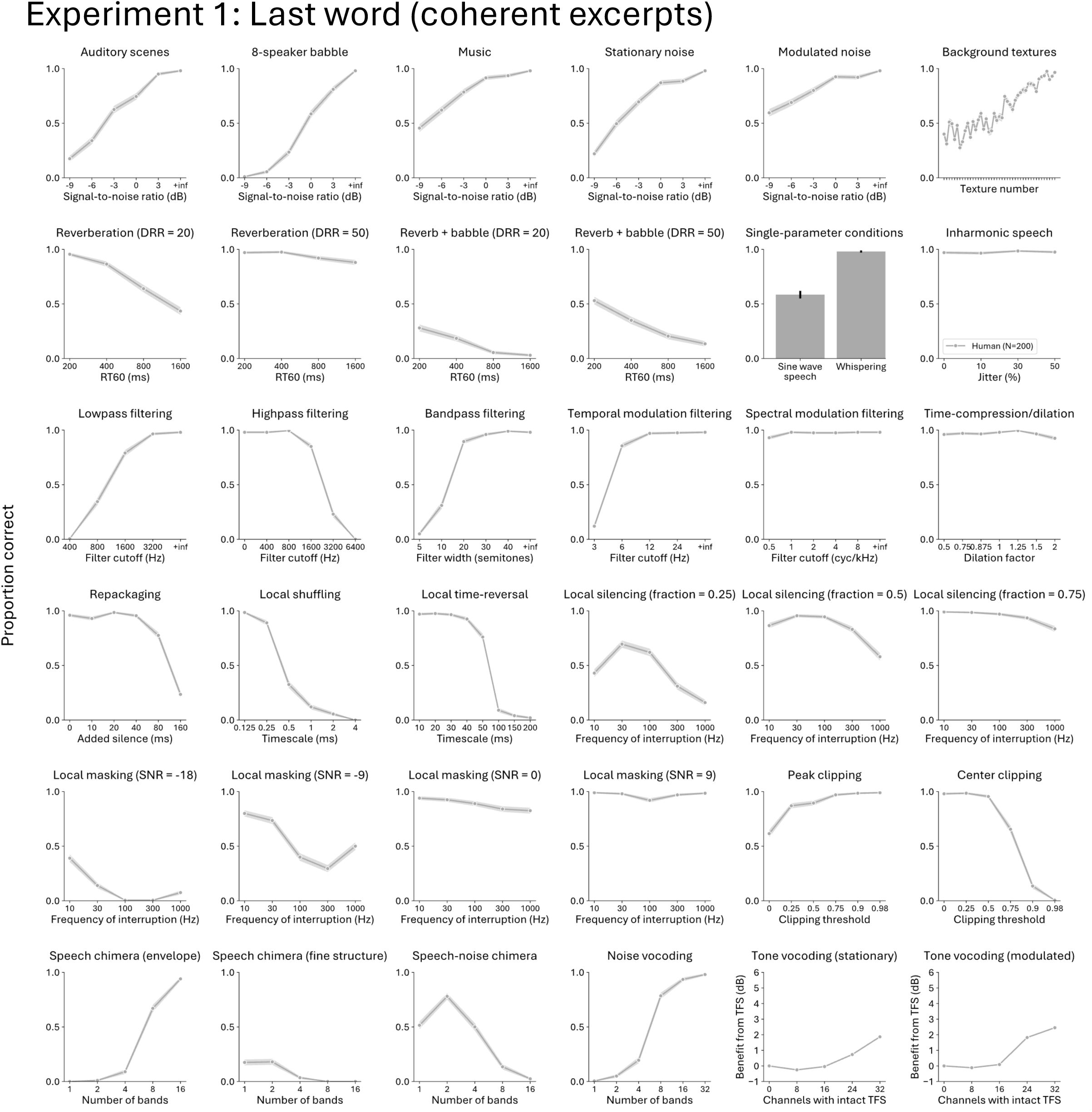
Human results on each distortion type for Experiment 1

**Supplementary Figure 4:**
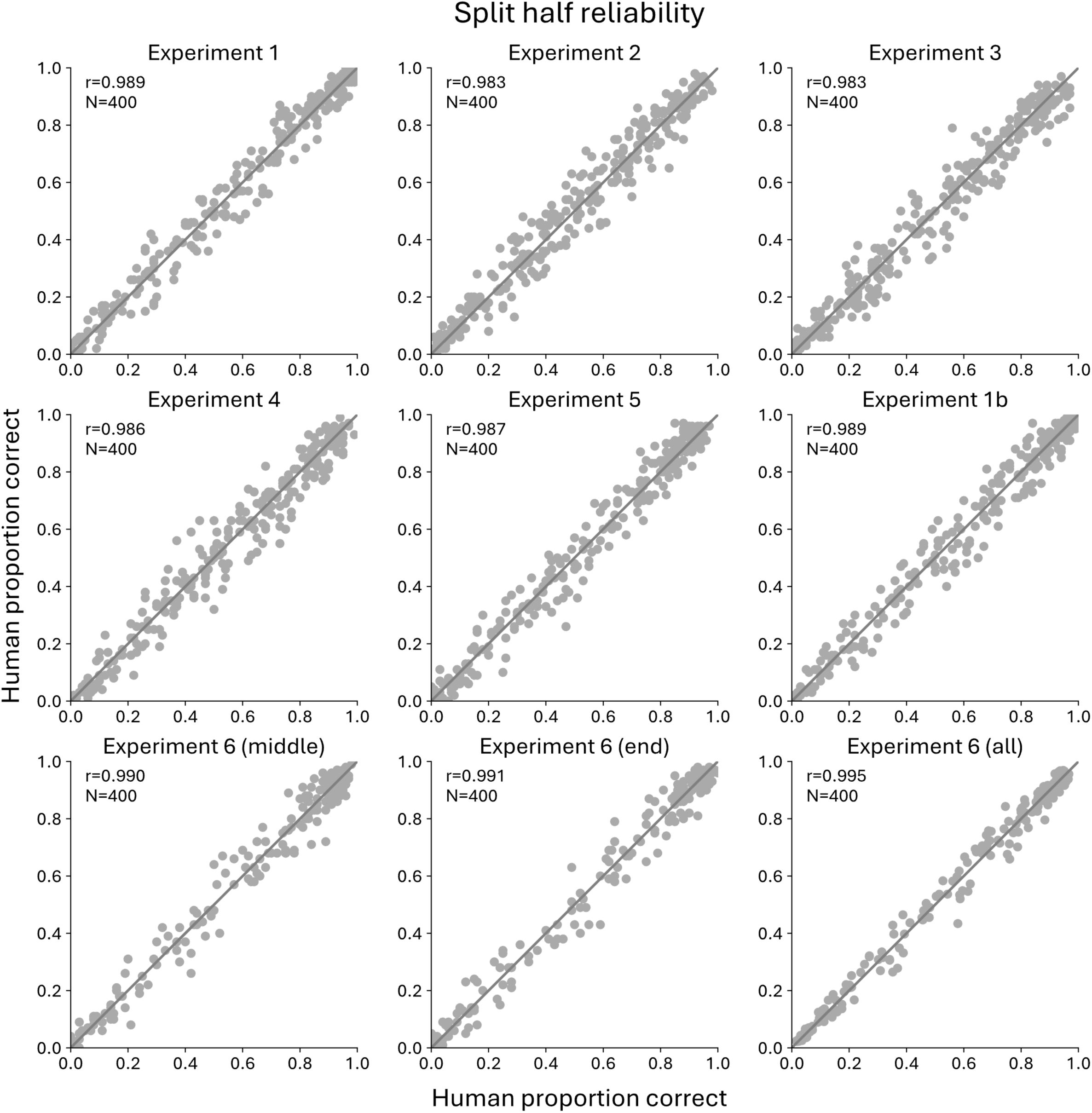
Split-half reliability of each experiment

**Supplementary Figure 5:**
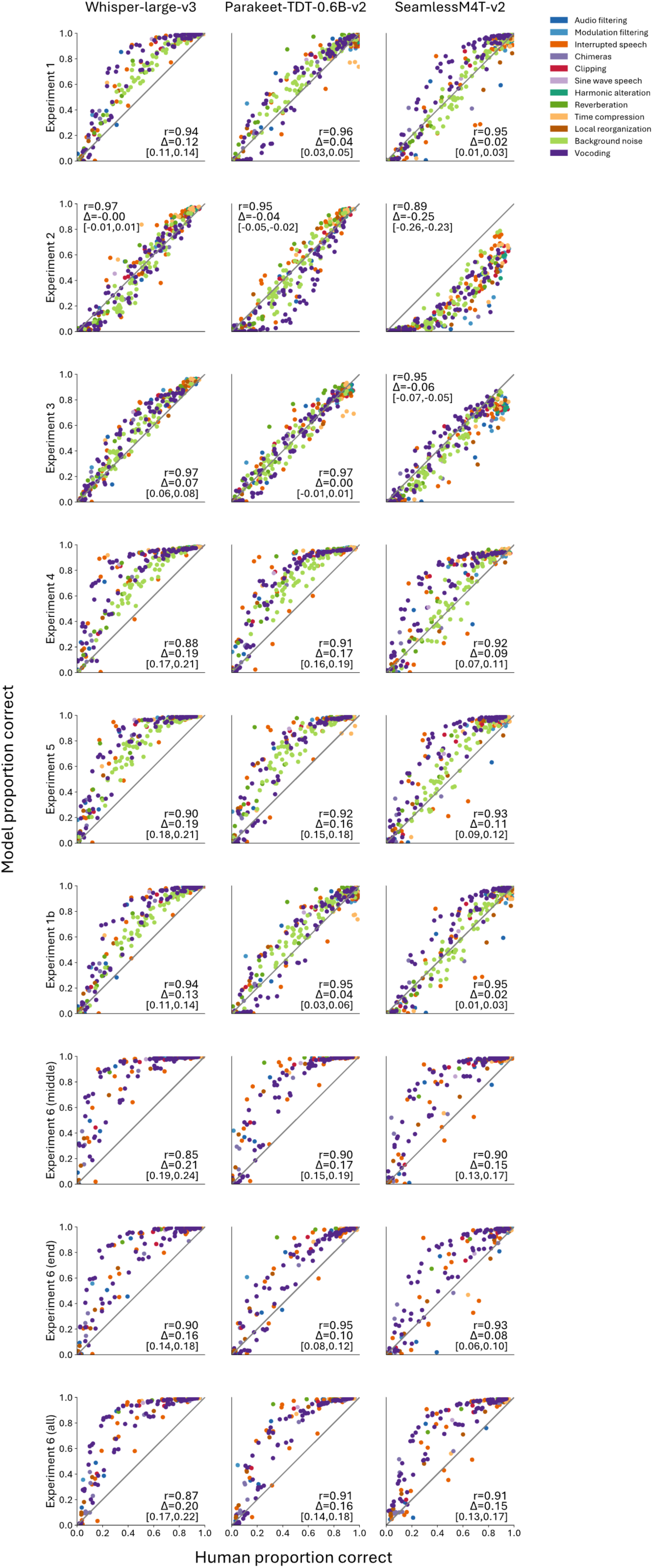
Color coded scatter plots of performance for all experiments

**Supplementary Figure 6:**
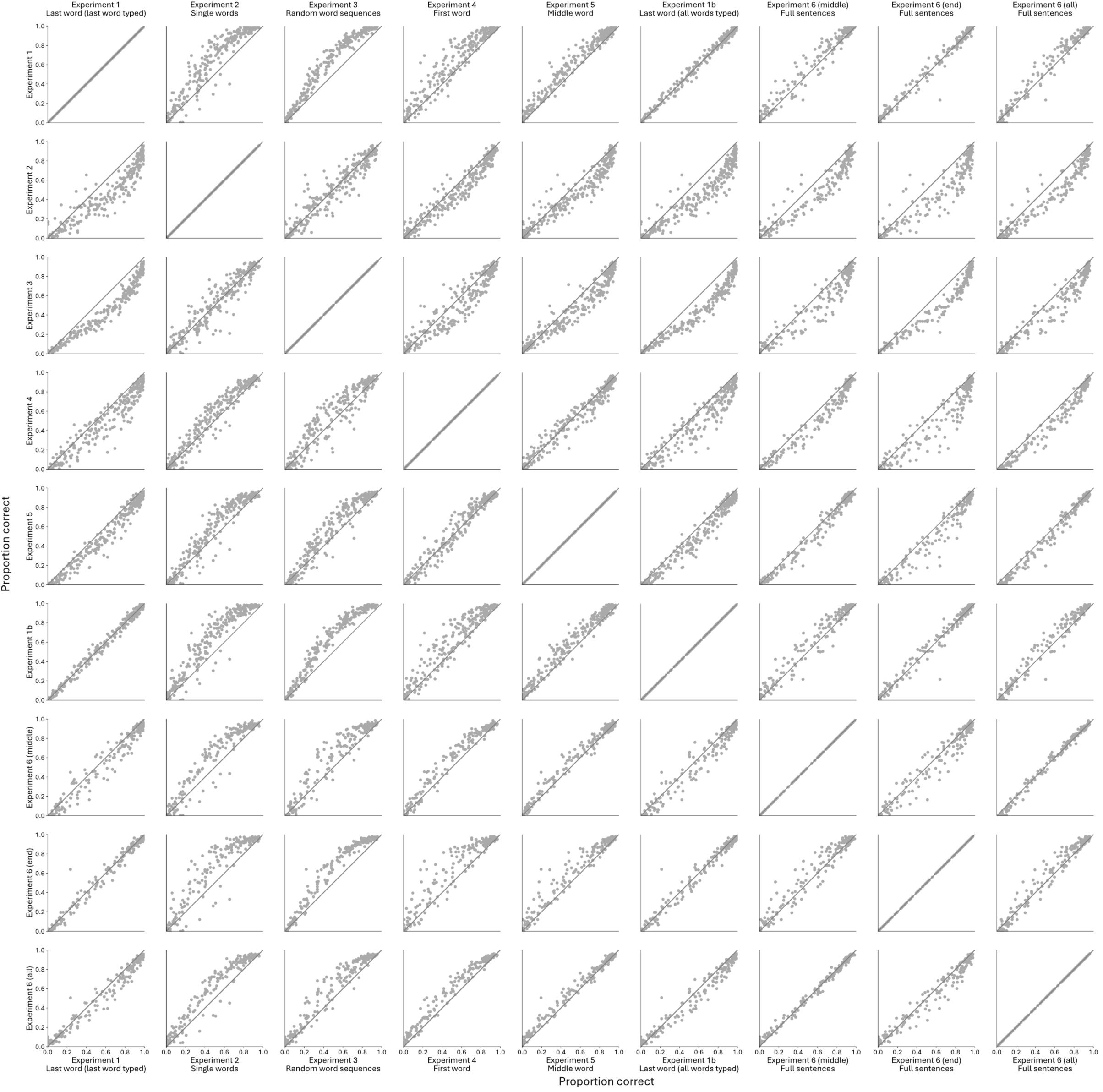
Human vs. human performance on all experiments

**Supplementary Figure 7:**
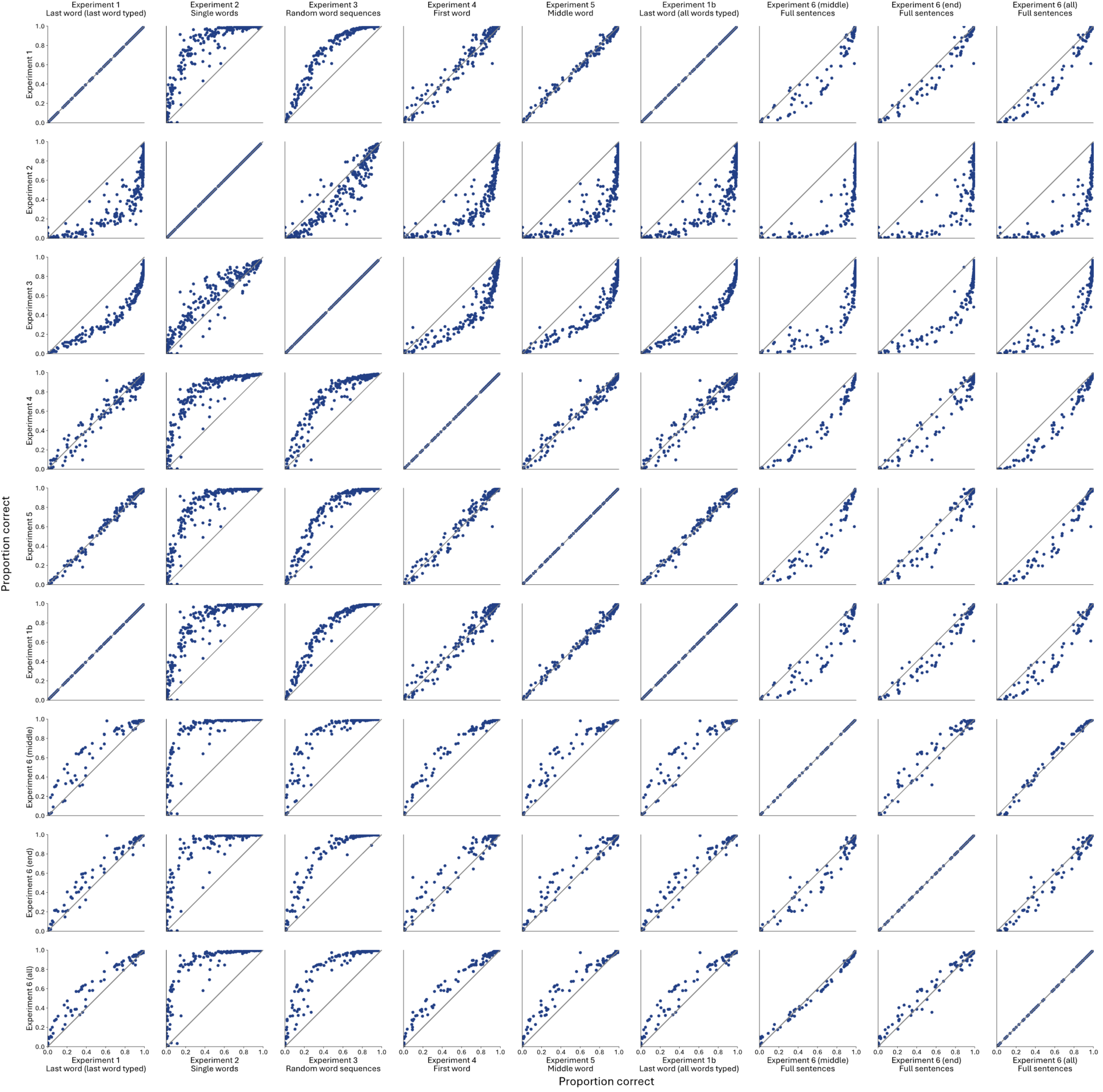
Whisper-large-v3 vs. Whisper-large-v3 performance on all experiments

**Supplementary Figure 8:**
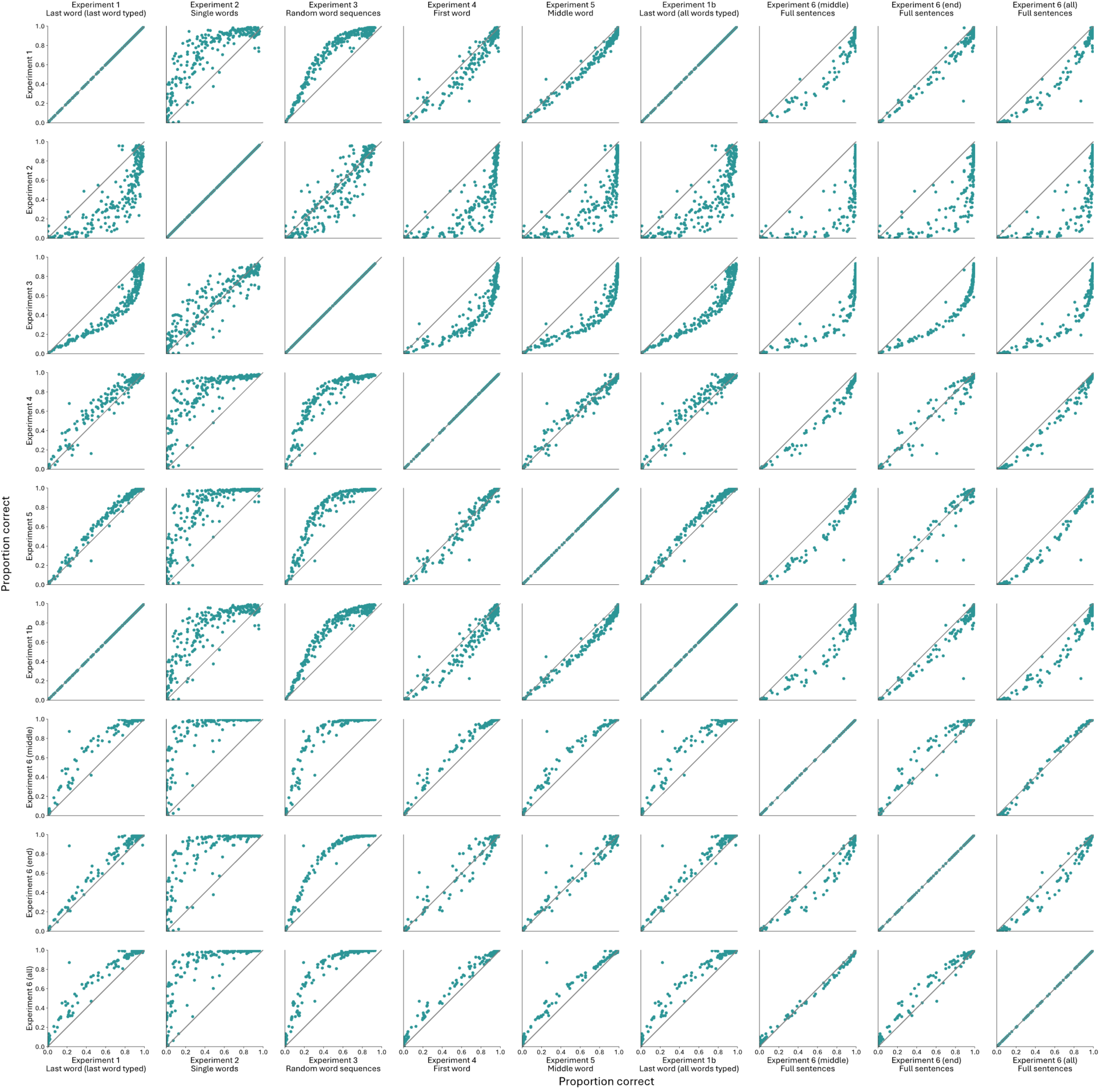
Parakeet vs. Parakeet performance on all experiments

**Supplementary Figure 9:**
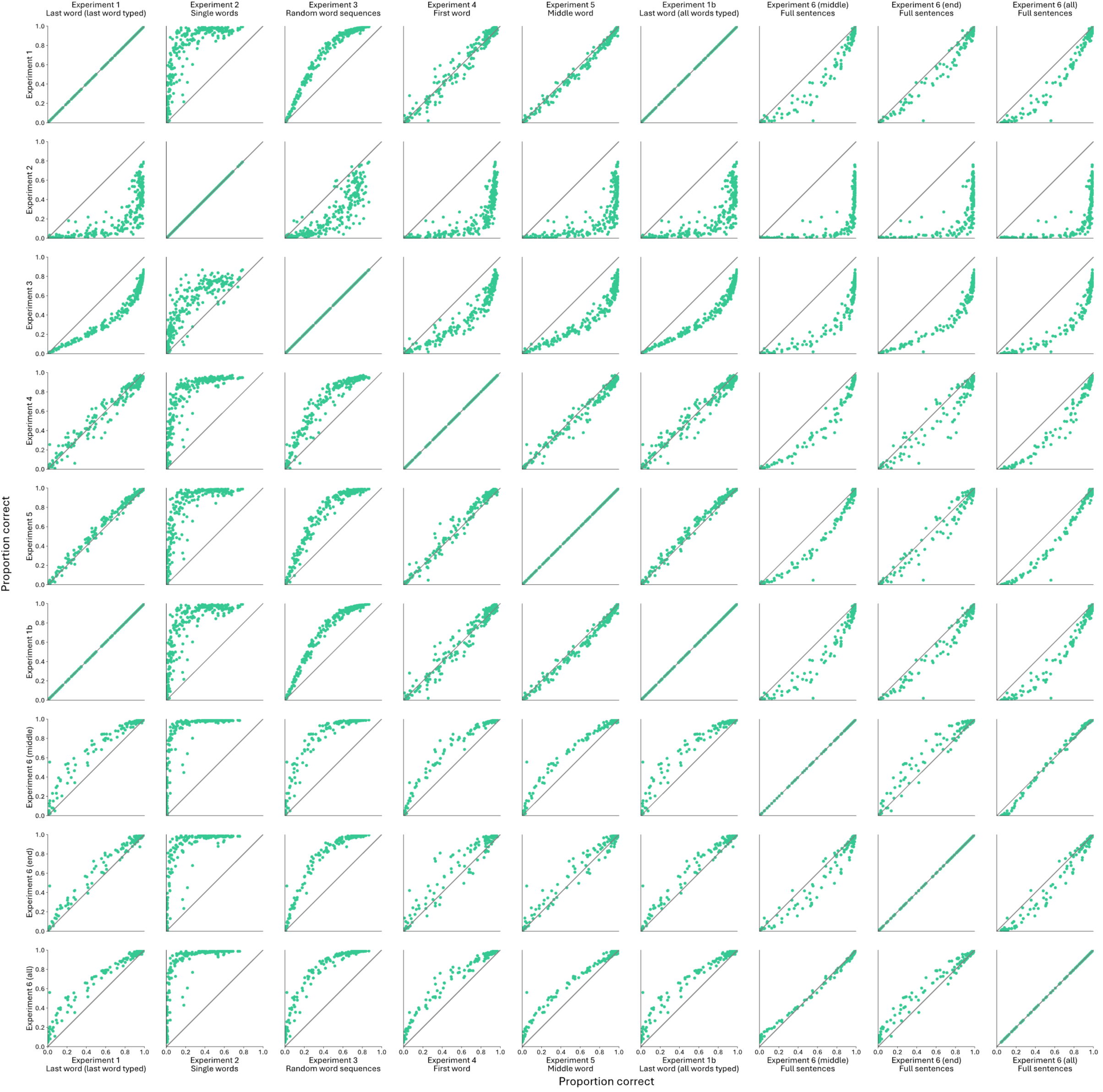
Seamless vs. Seamless performance on all experiments

**Supplementary Figure 10:**
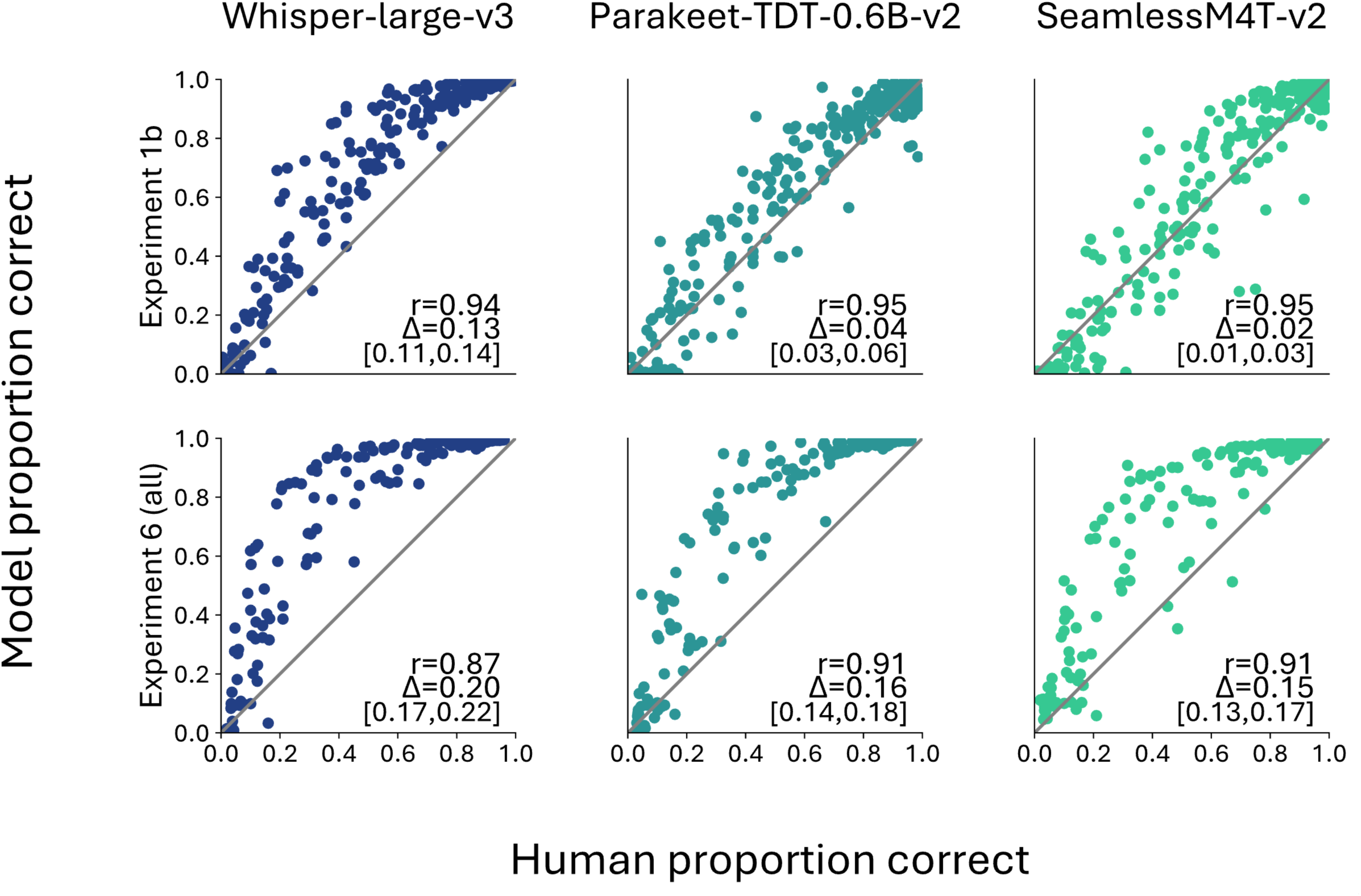
Scatter plots of performance for Experiment 1.5 and Experiment 1.6 (all)

**Supplementary Figure 11:**
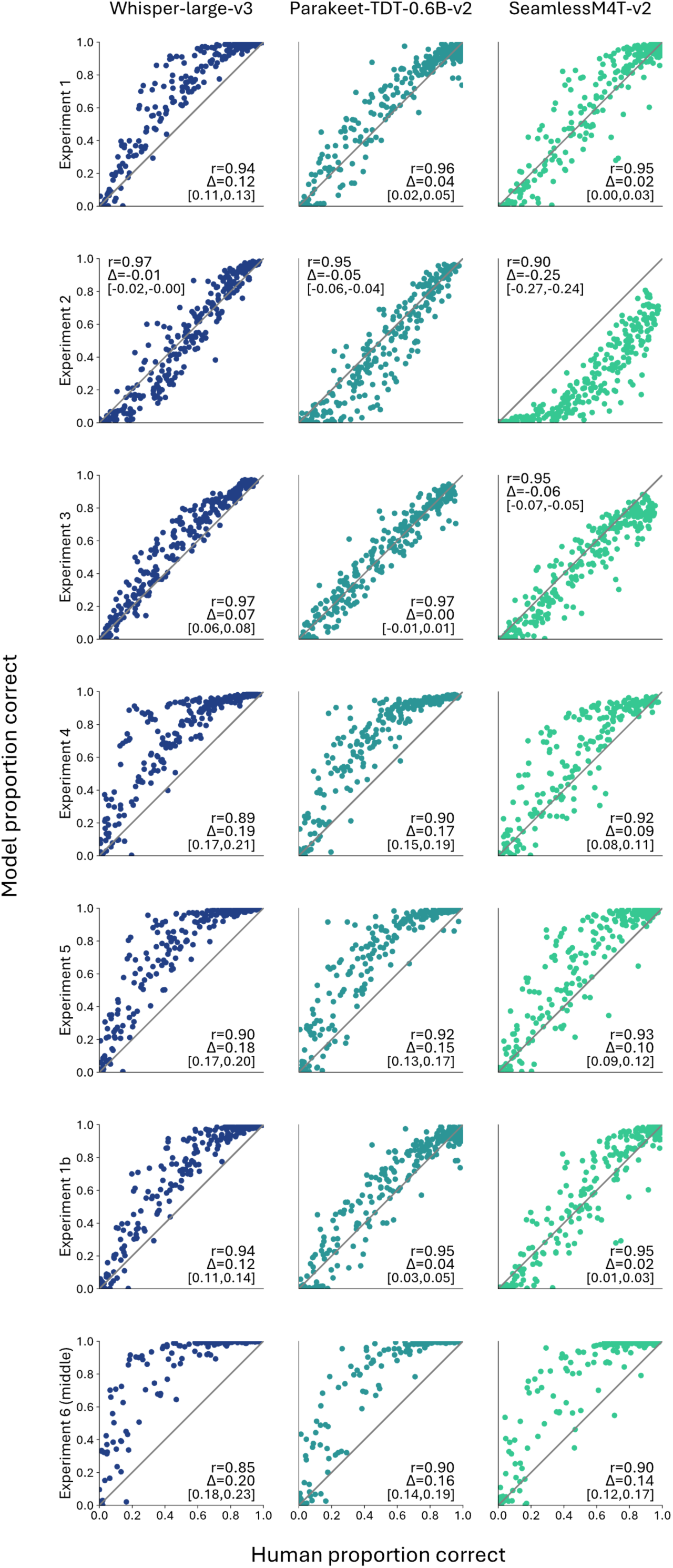
Scatter plots of performance for only the words common to all experiments

**Supplementary Figure 12:**
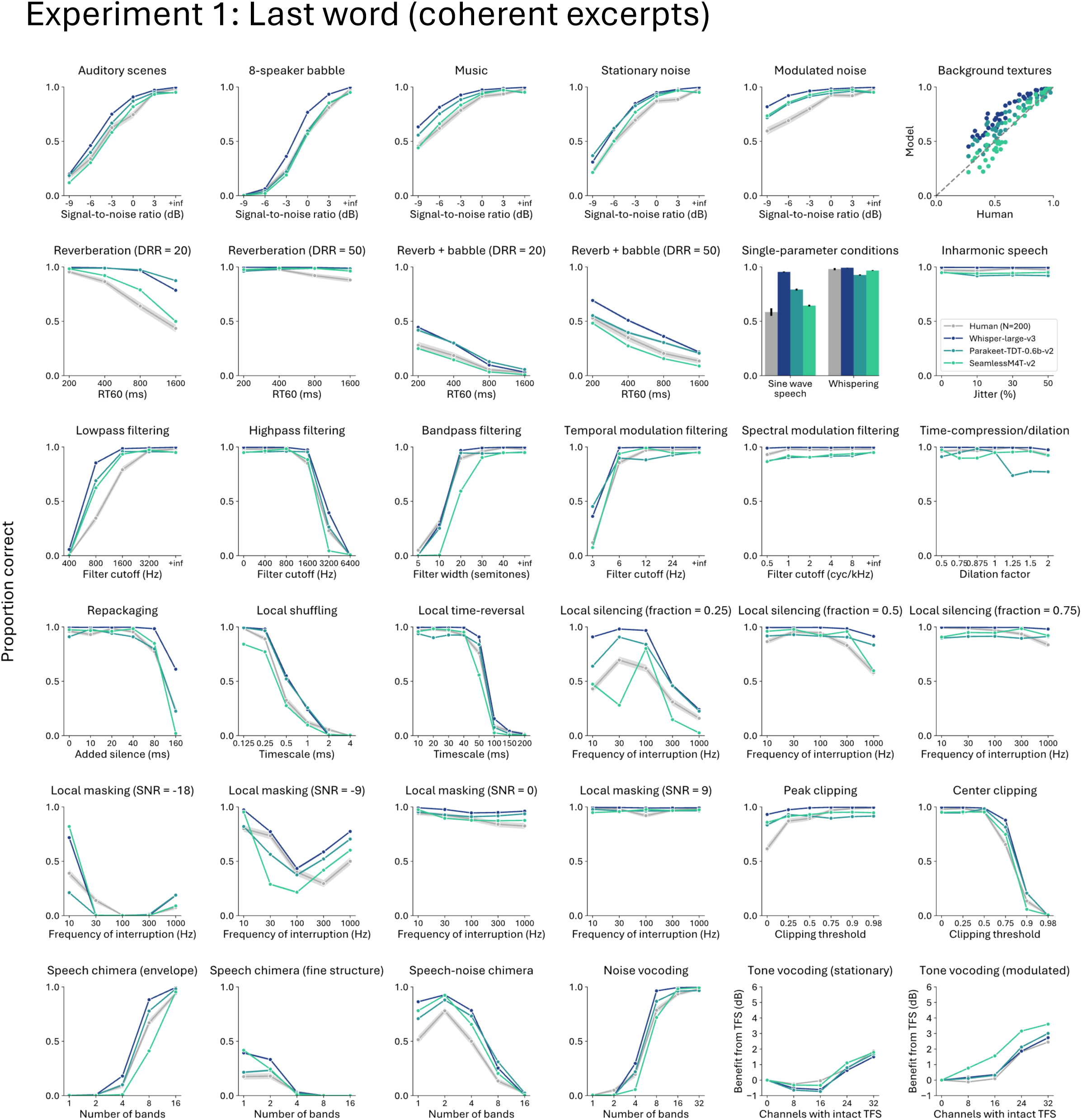
Industry model and human results on each distortion type for Experiment 1

**Supplementary Figure 13:**
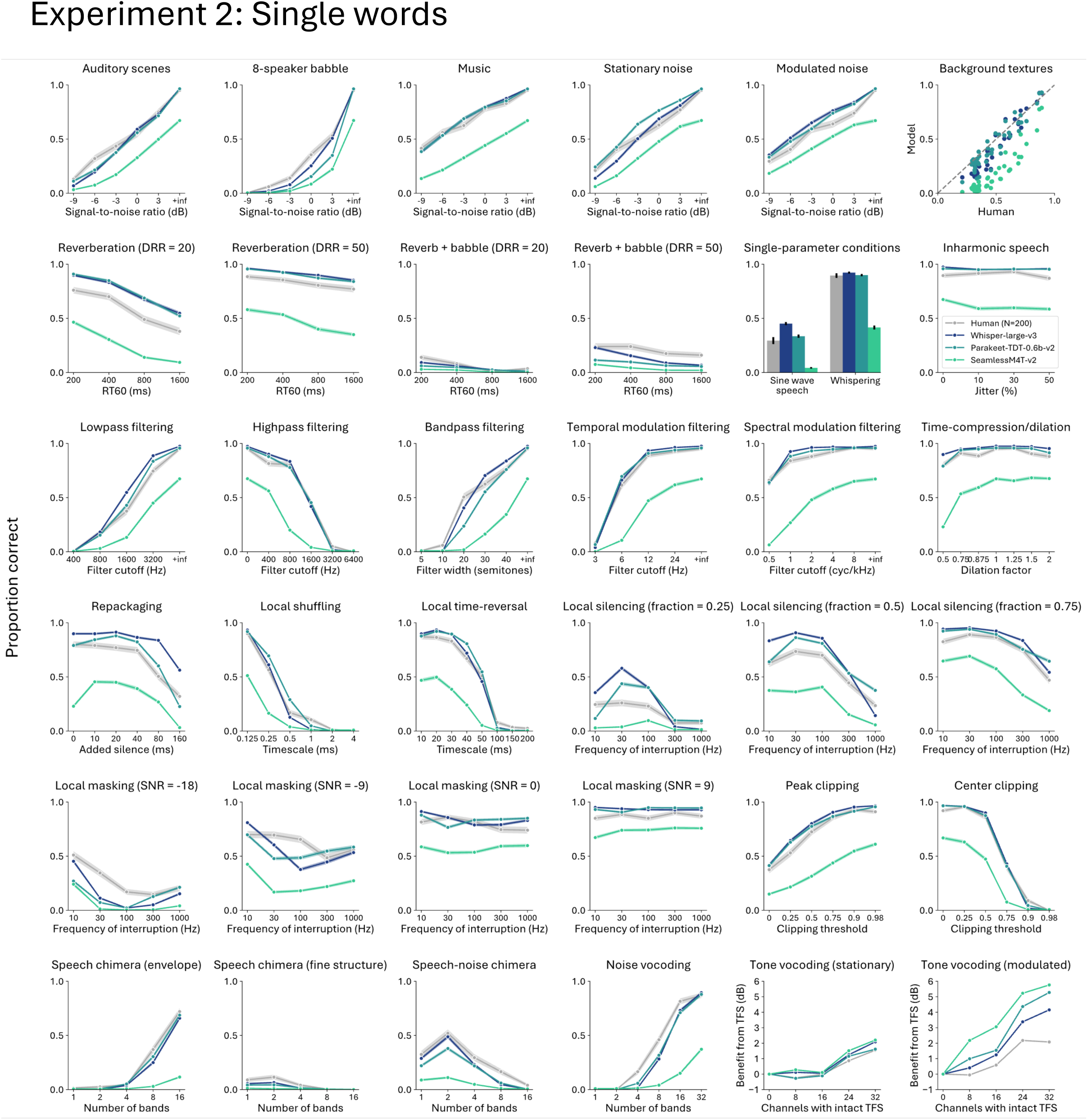
Industry model and human results on each distortion type for Experiment 2

**Supplementary Figure 14:**
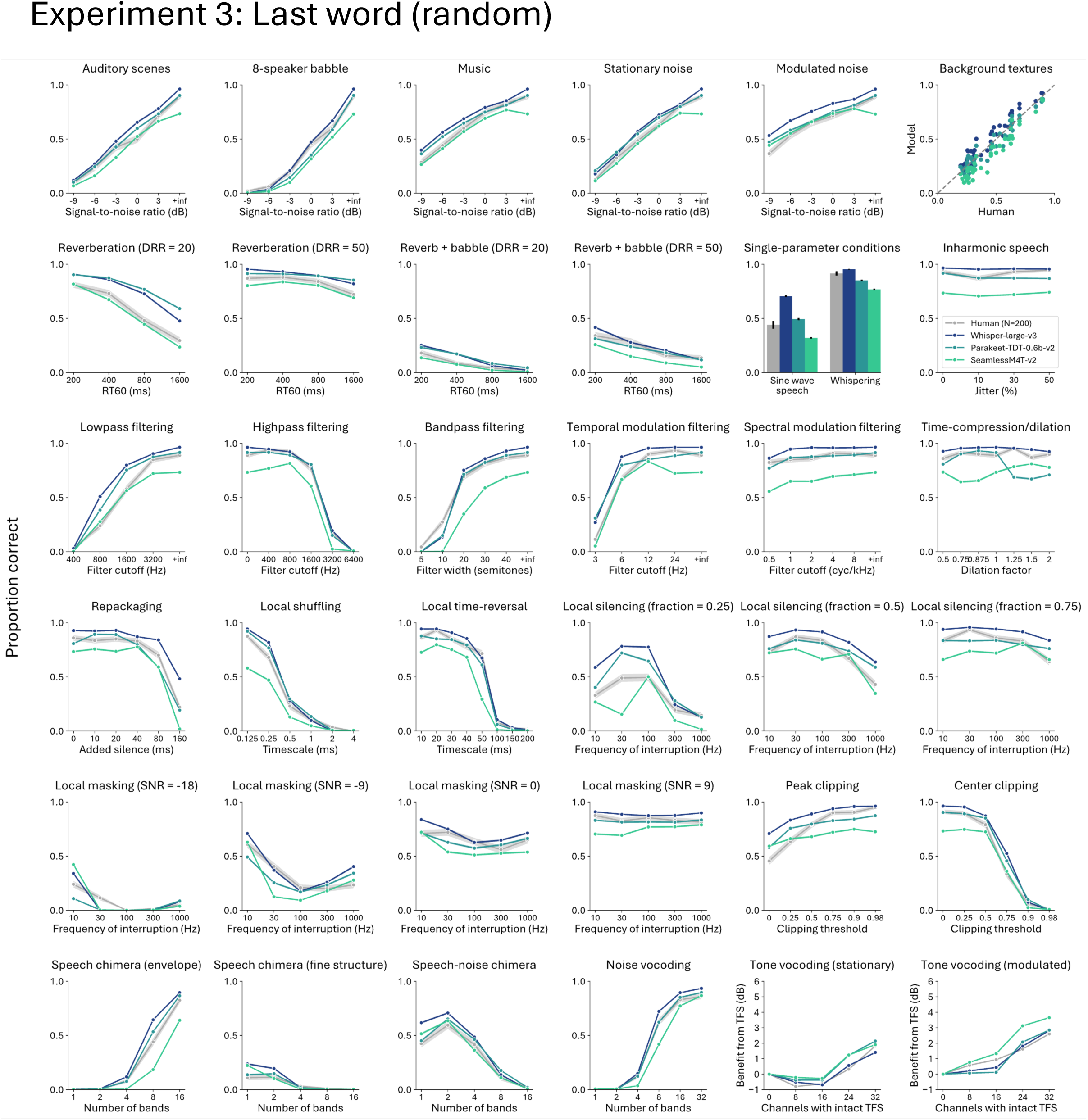
Industry model and human results on each distortion type for Experiment 3

**Supplementary Figure 15:**
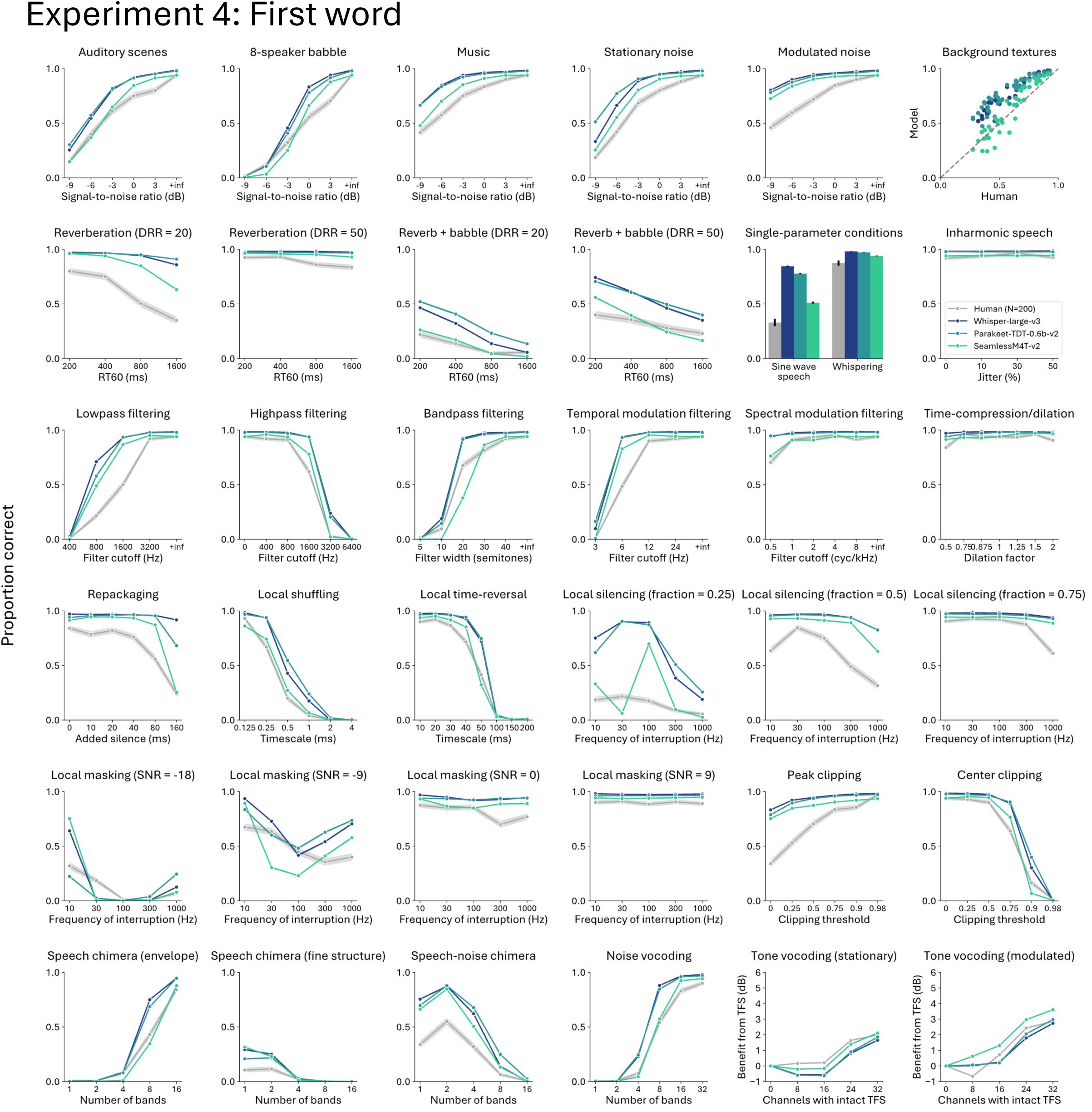
Industry model and human results on each distortion type for Experiment 4

**Supplementary Figure 16:**
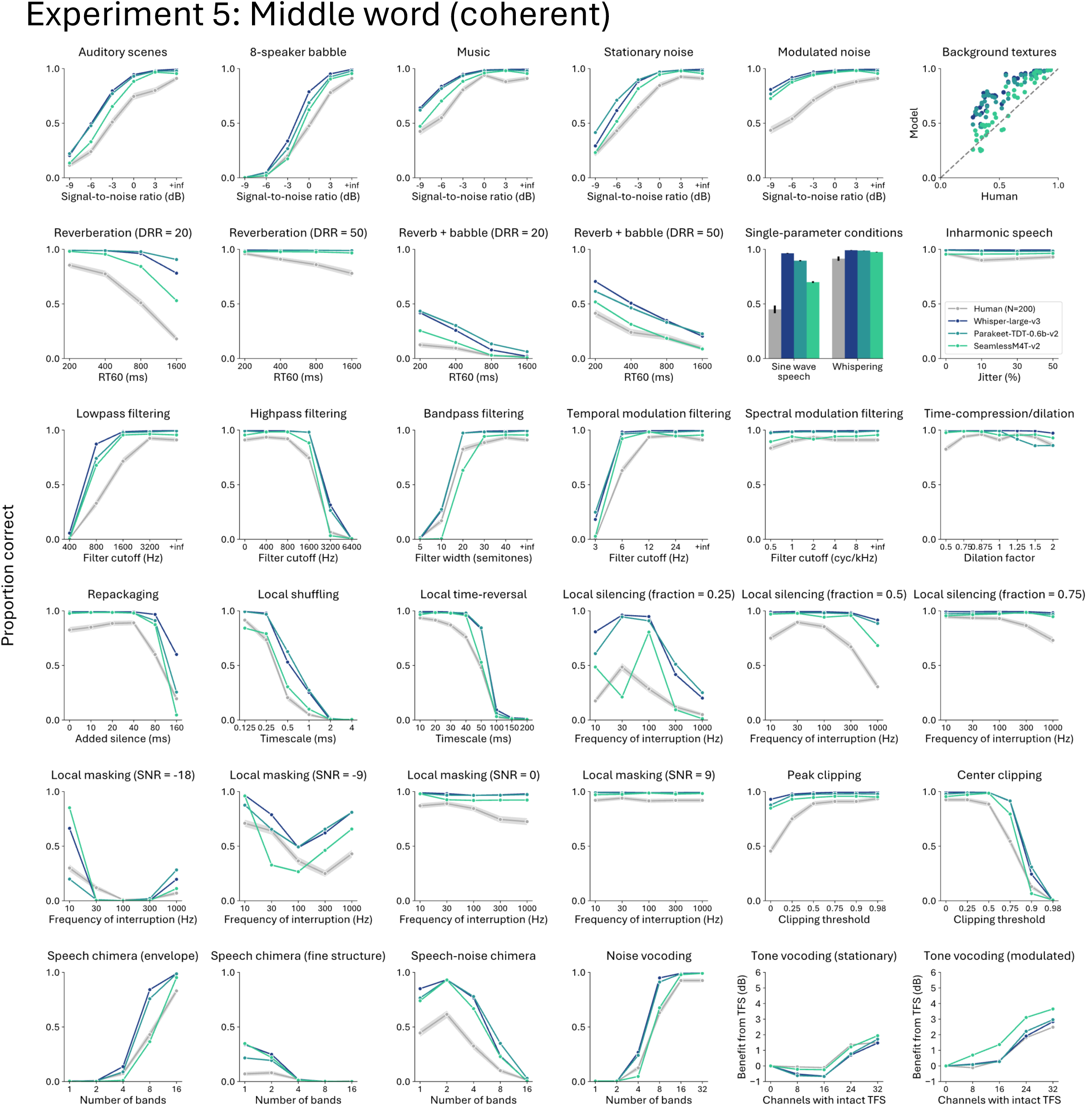
Industry model and human results on each distortion type for Experiment 5

**Supplementary Figure 17:**
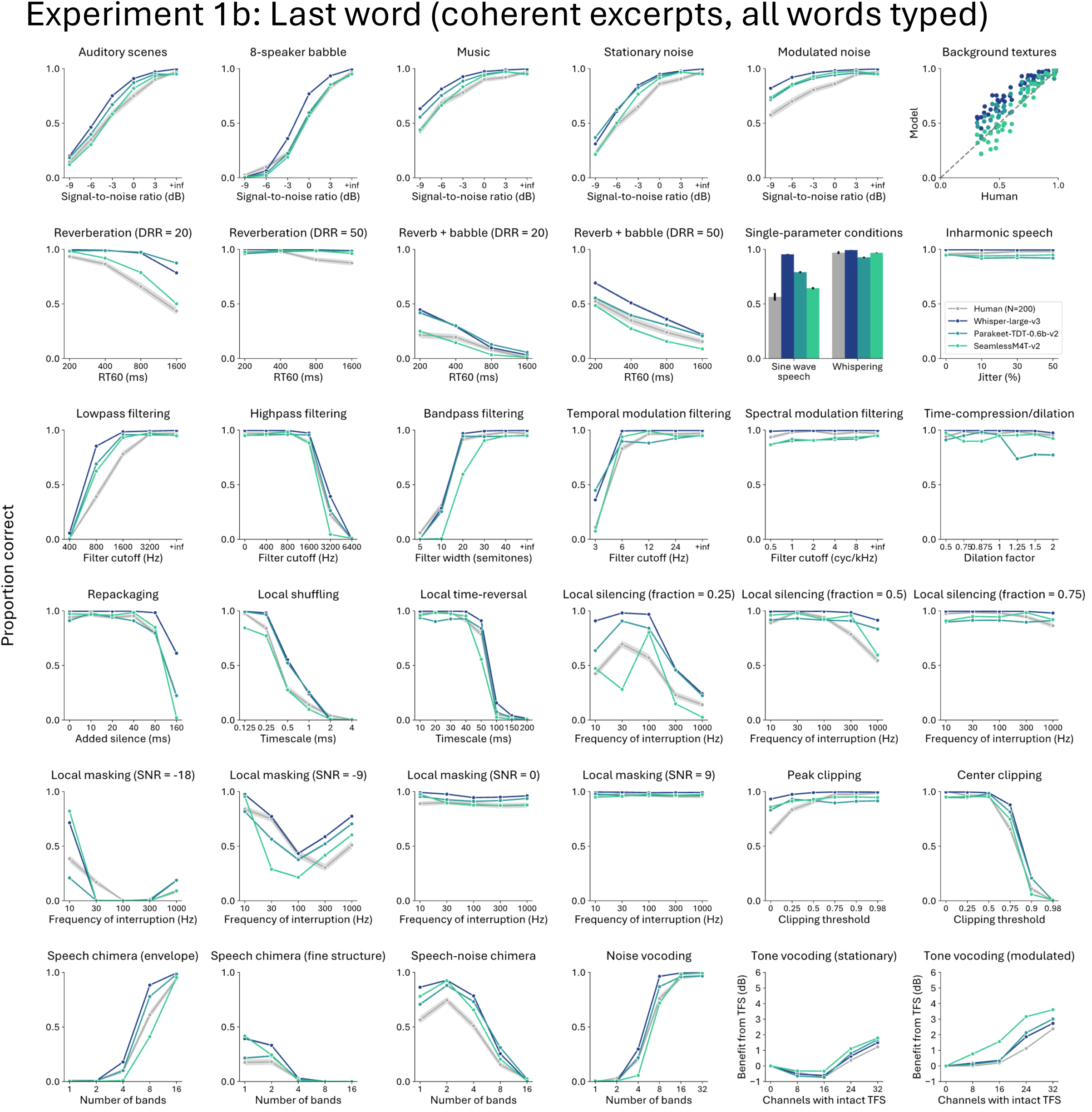
Industry model and human results on each distortion type for Experiment 1b

**Supplementary Figure 18:**
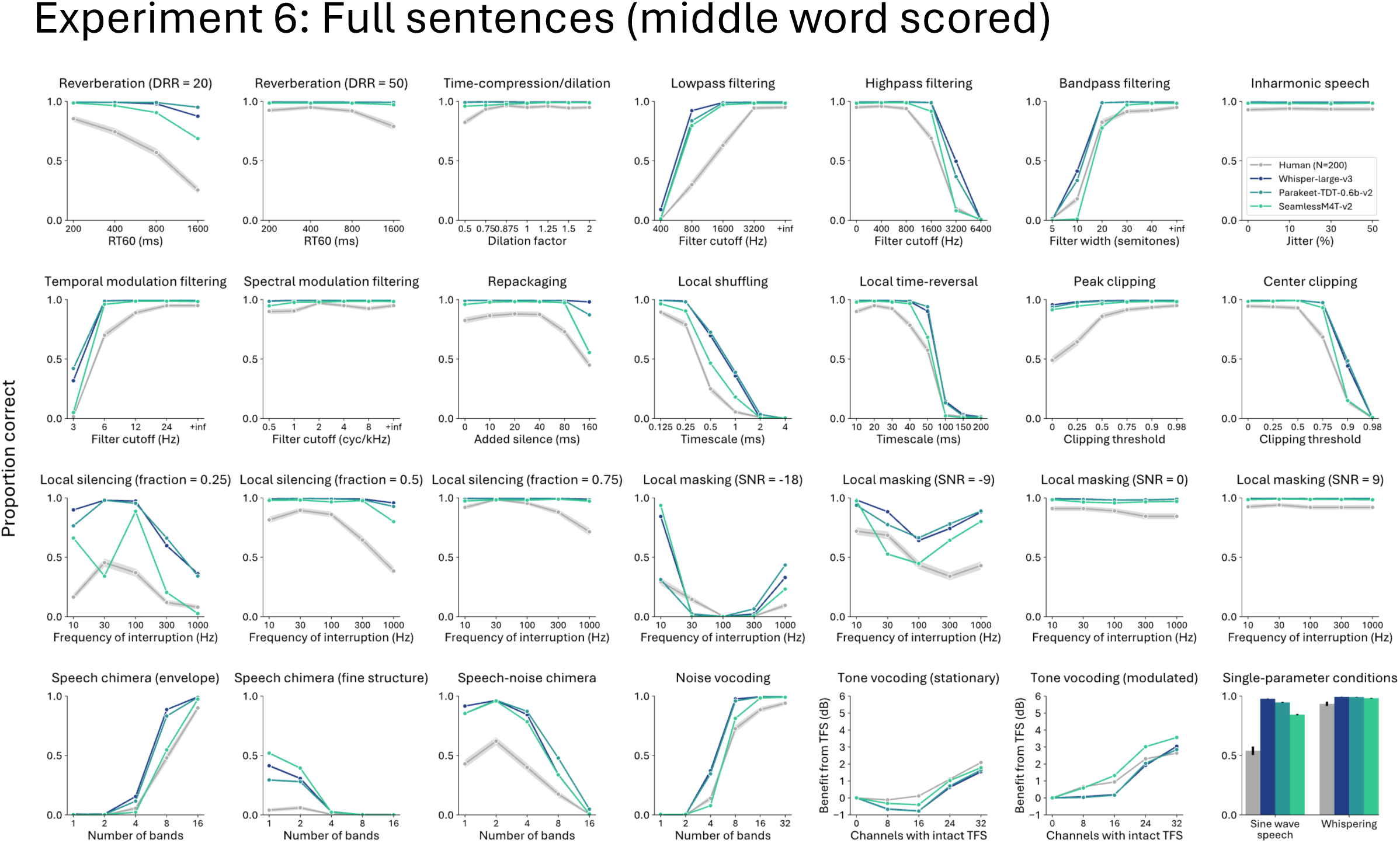
Industry model and human results on each distortion type for Experiment 6 (middle word scored)

**Supplementary Figure 19:**
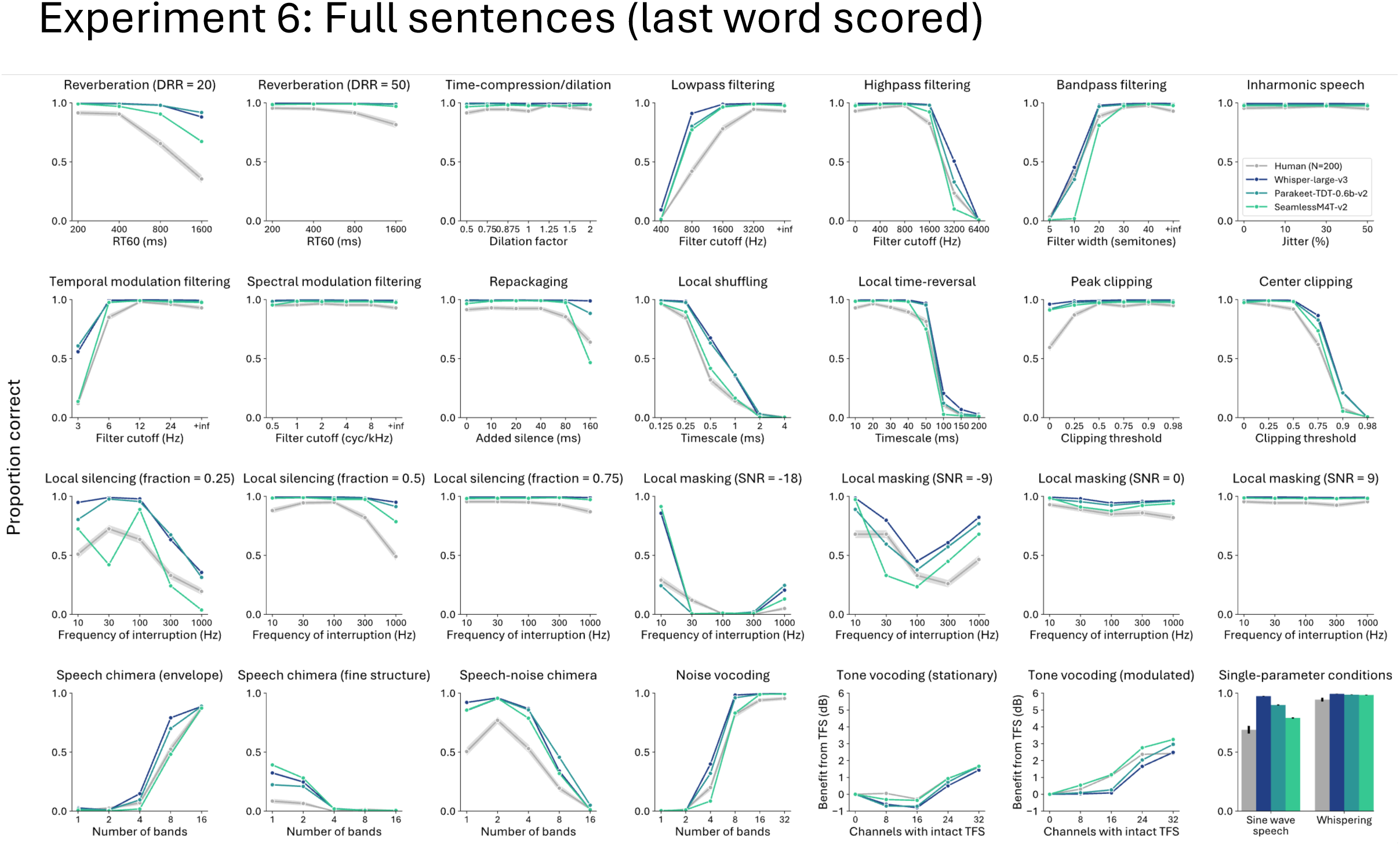
Industry model and human results on each distortion type for Experiment 6 (last word scored)

**Supplementary Figure 20:**
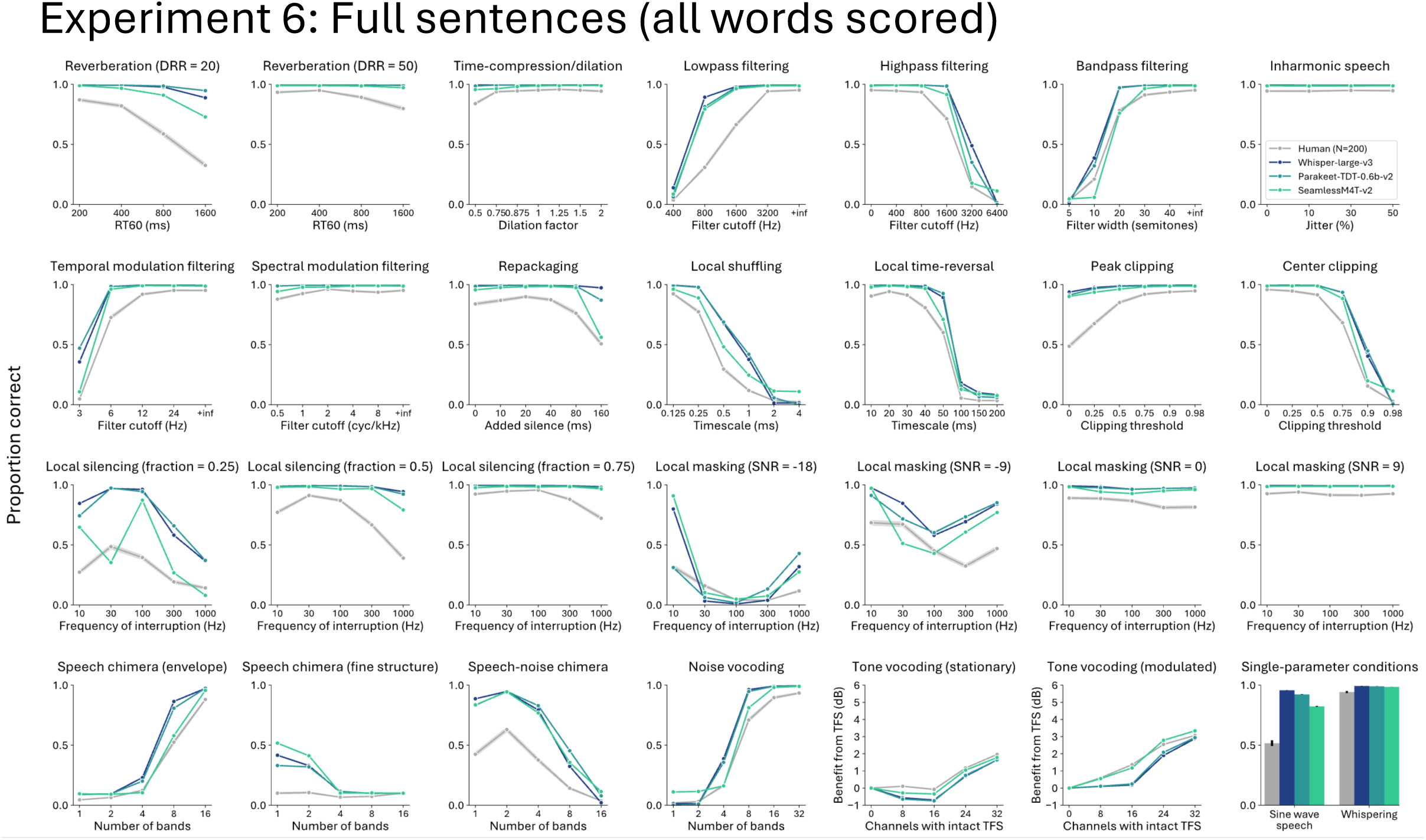
Industry model and human results on each distortion type for Experiment 6 (all words scored)

**Supplementary Figure 21:**
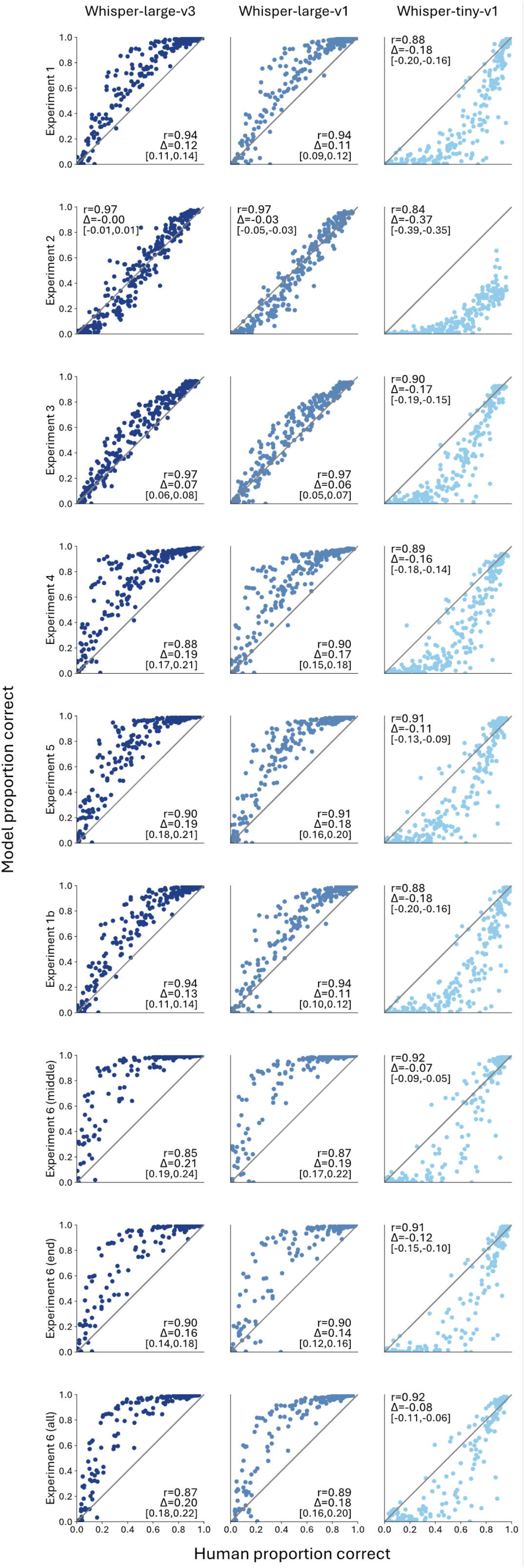
Scatter plots of performance for Whisper variants on all experiments

**Supplementary Figure 22:**
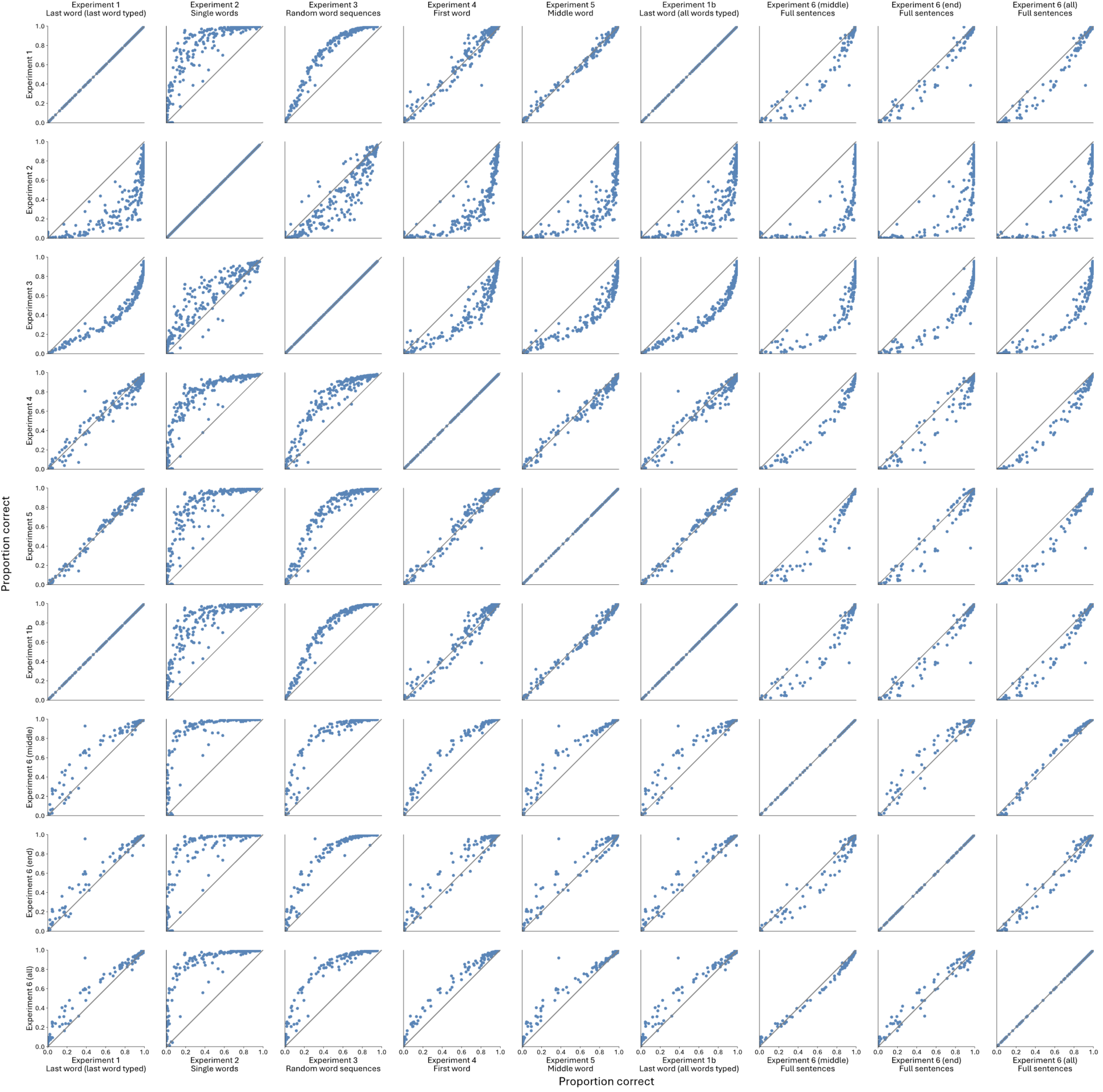
Whisper-large-v1 vs. Whisper-large-v1 performance on all experiment

**Supplementary Figure 23:**
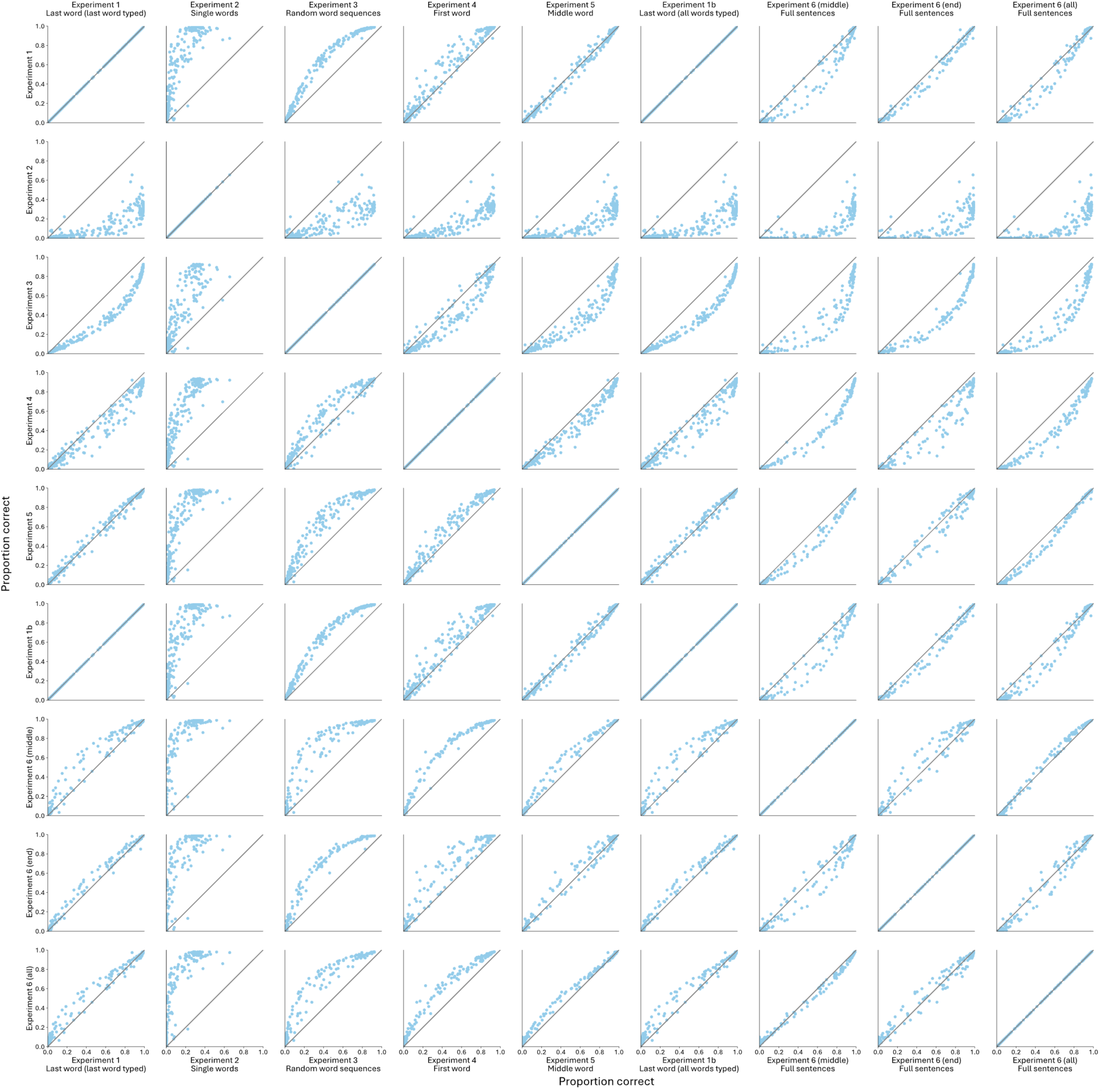
Whisper-tiny-v1 vs. Whisper-tiny-v1 performance on all experiments

**Supplementary Figure 24:**
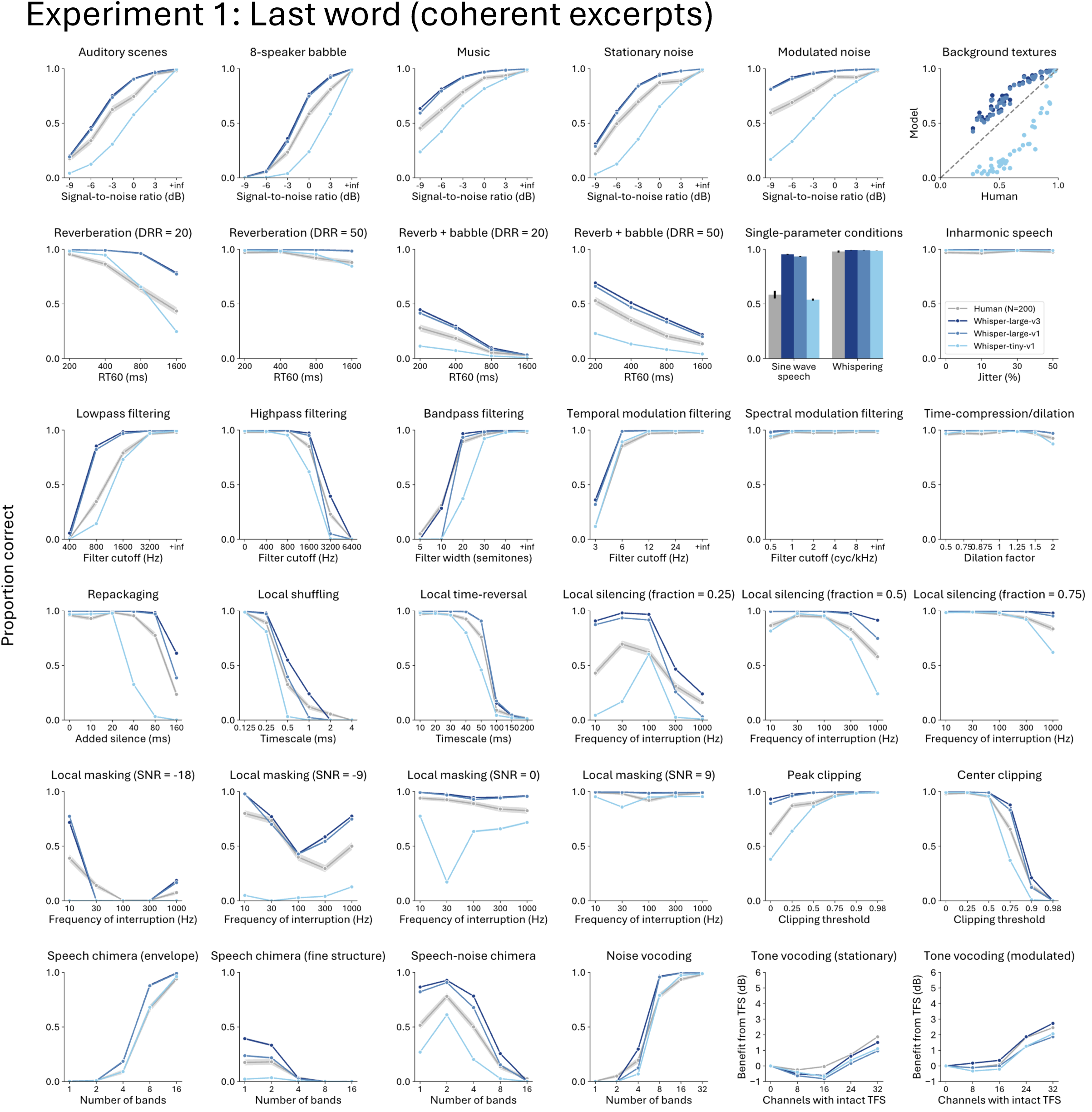
Whisper variants and human results on each distortion type for Experiment 1

**Supplementary Figure 25:**
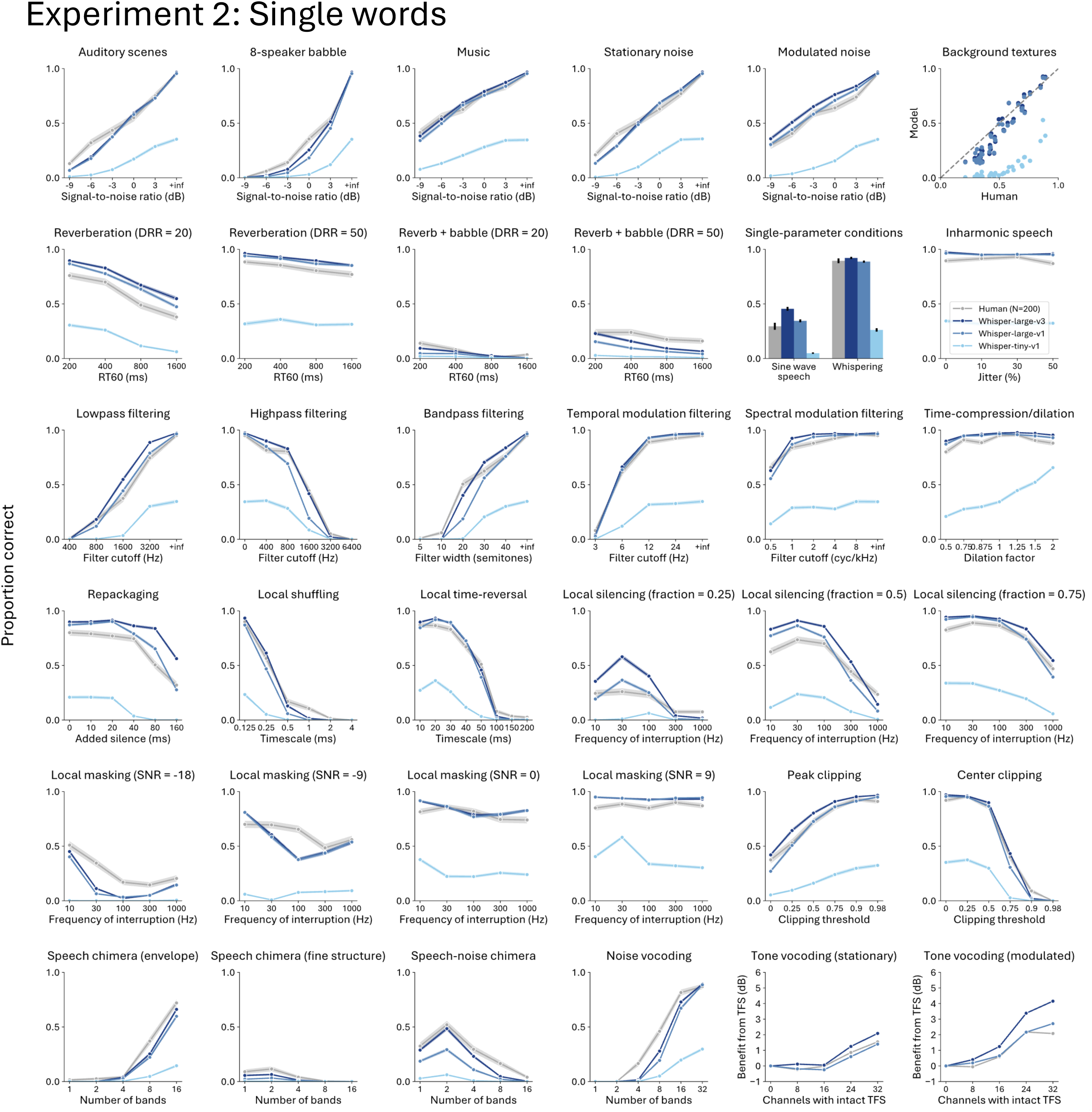
Whisper variants and human results on each distortion type for Experiment 2

**Supplementary Figure 26:**
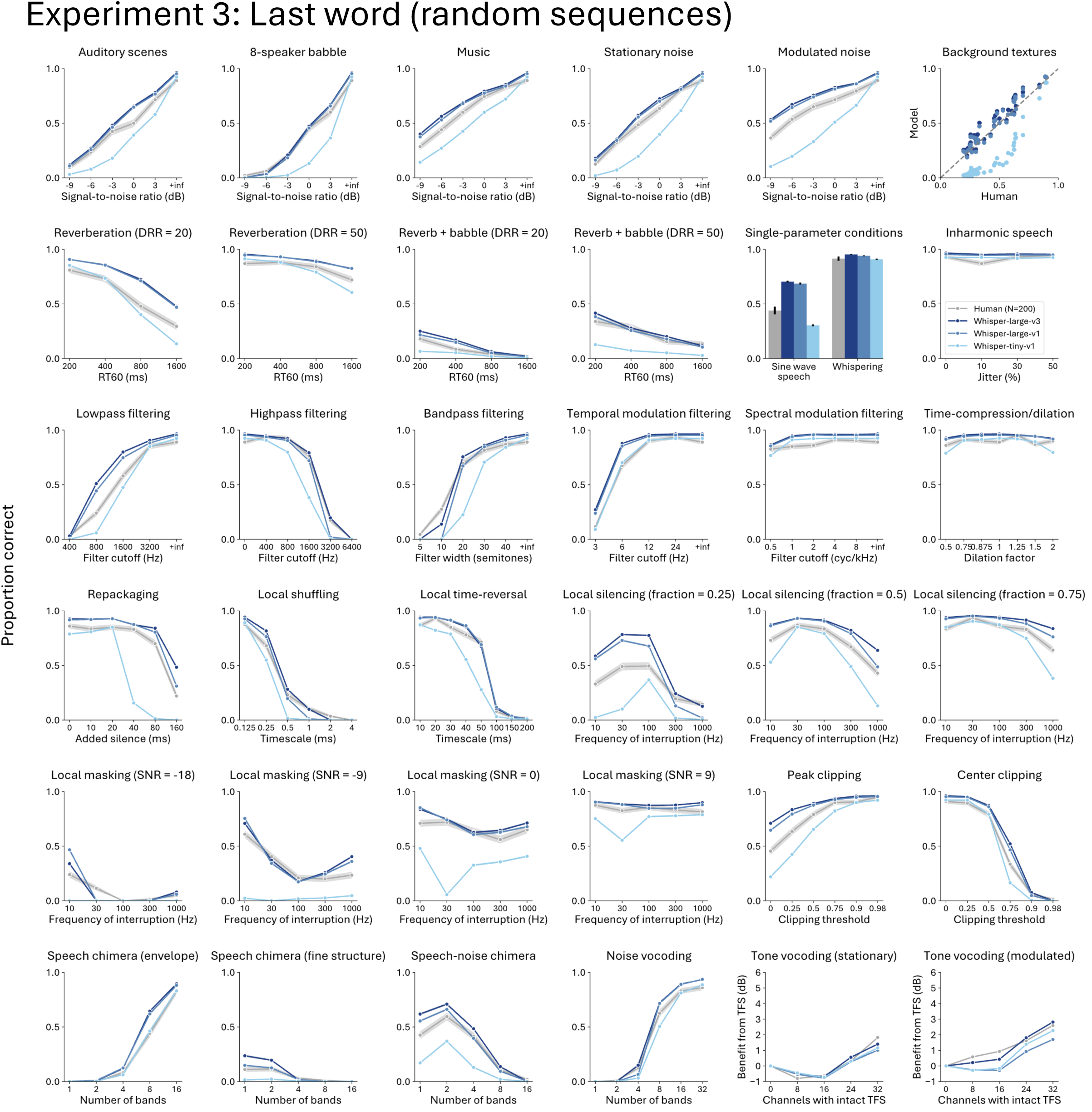
Whisper variants and human results on each distortion type for Experiment 3

**Supplementary Figure 27:**
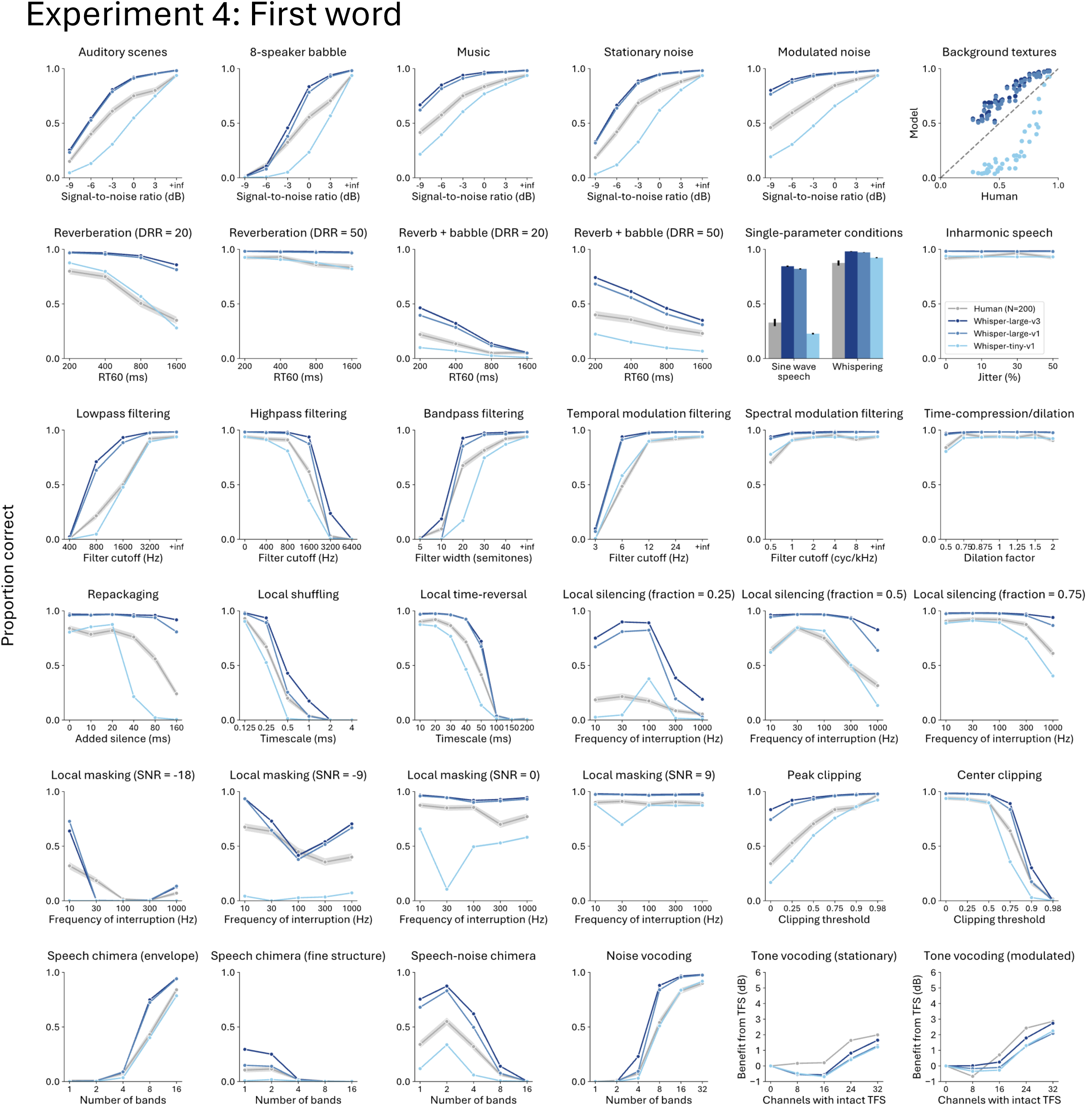
Whisper variants and human results on each distortion type for Experiment 4

**Supplementary Figure 28:**
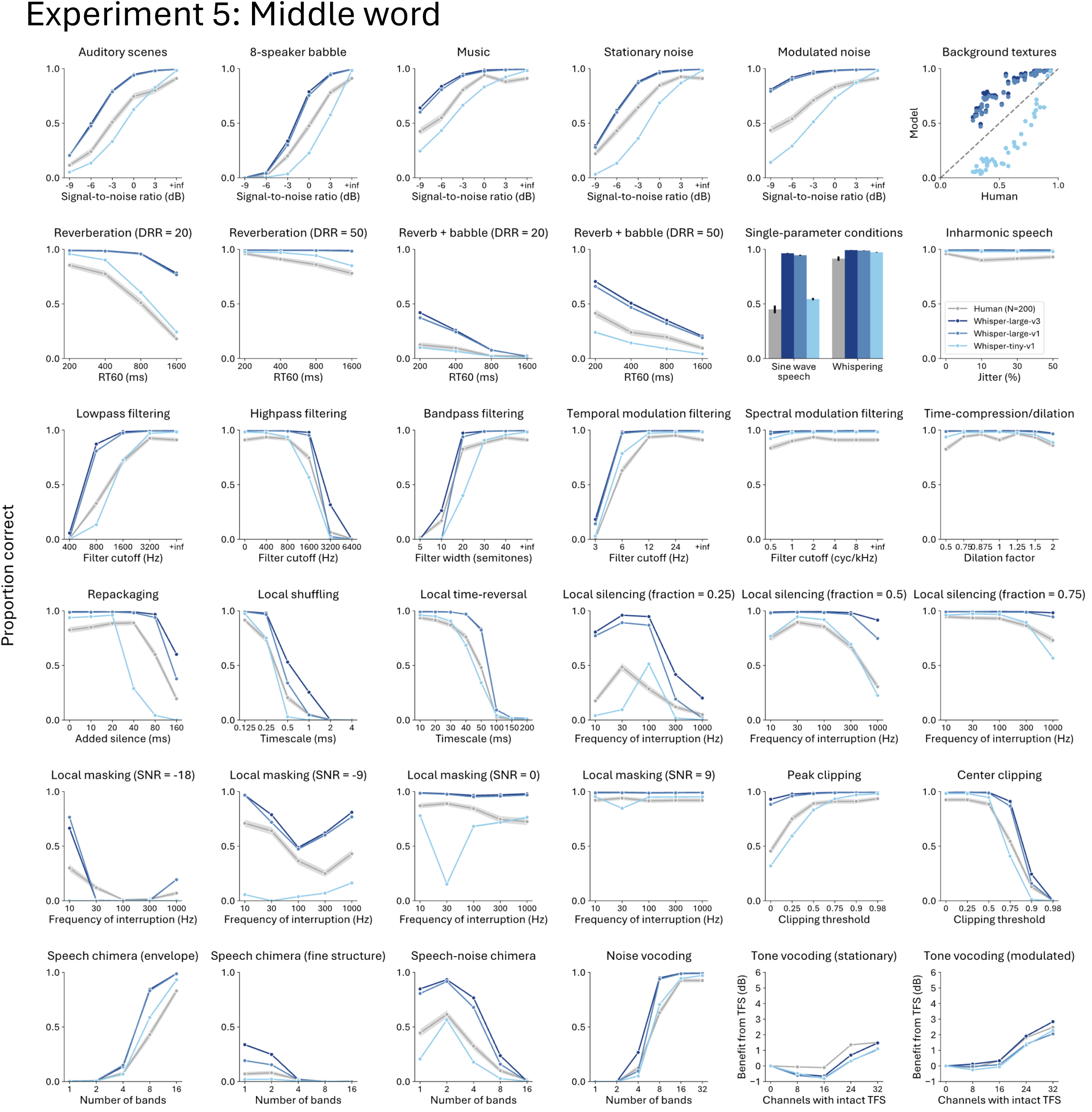
Whisper variants and human results on each distortion type for Experiment 5

**Supplementary Figure 29:**
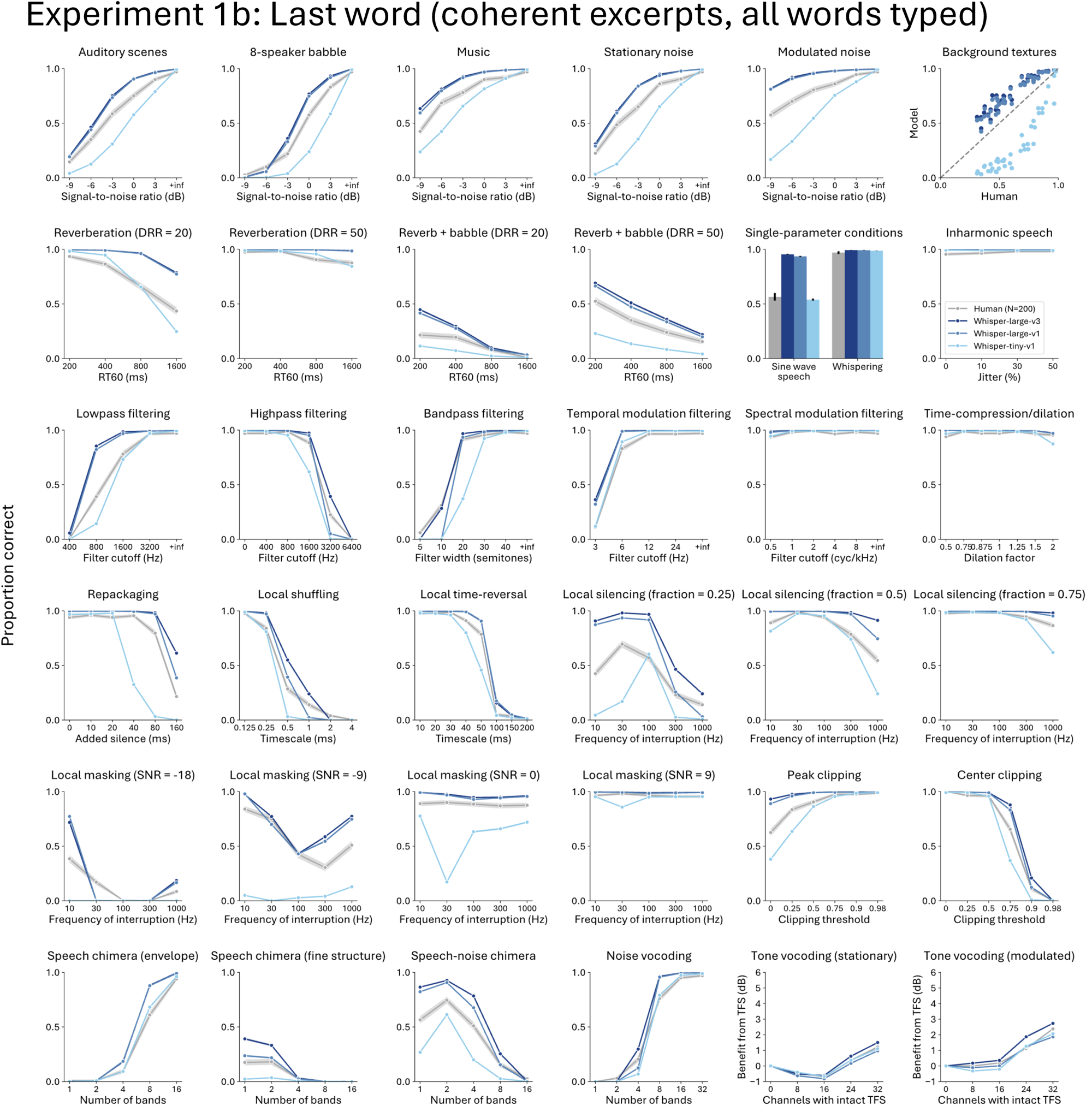
Whisper variants and human results on each distortion type for Experiment 1b

**Supplementary Figure 30:**
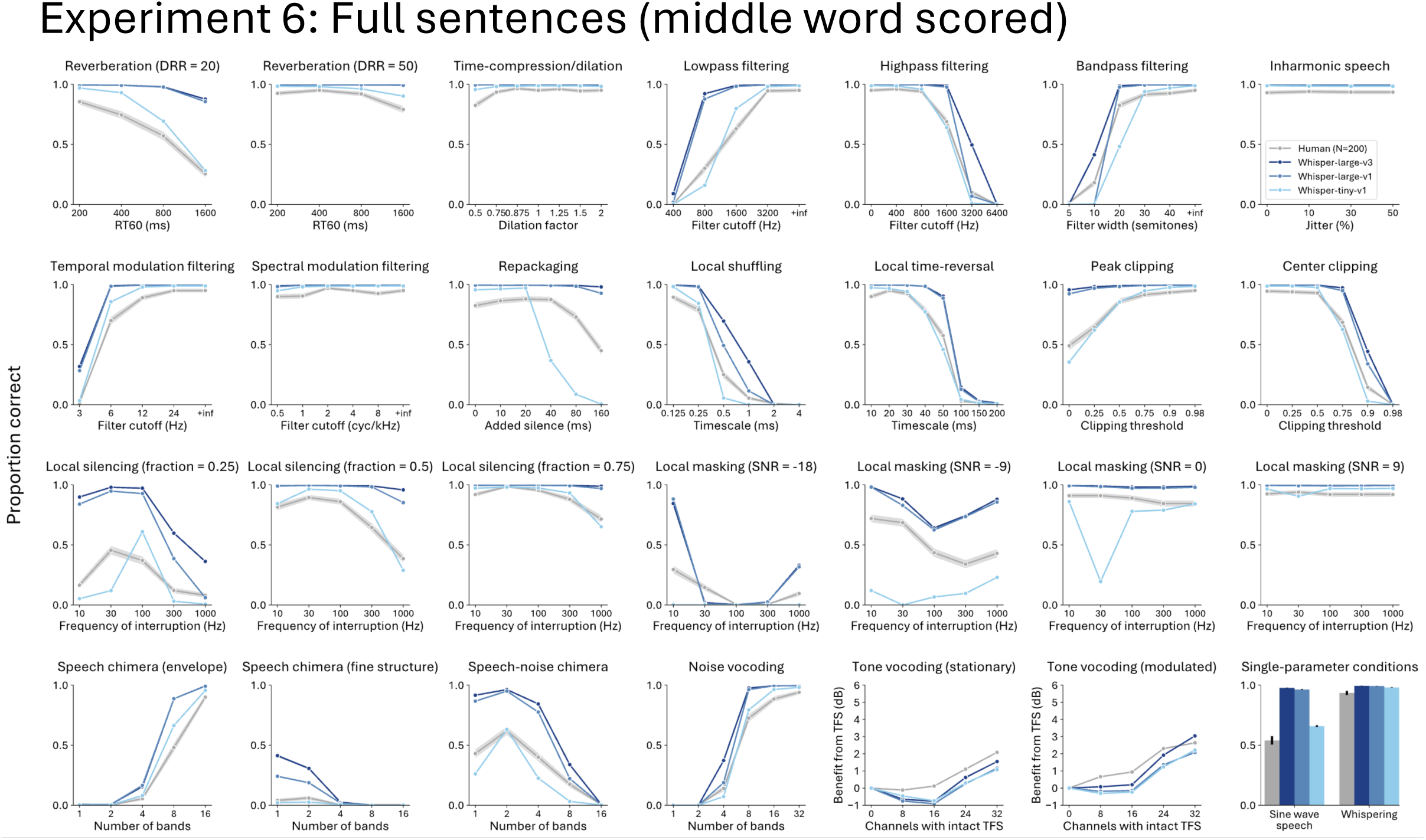
Whisper variants and human results on each distortion type for Experiment 6 (middle word scored)

**Supplementary Figure 31:**
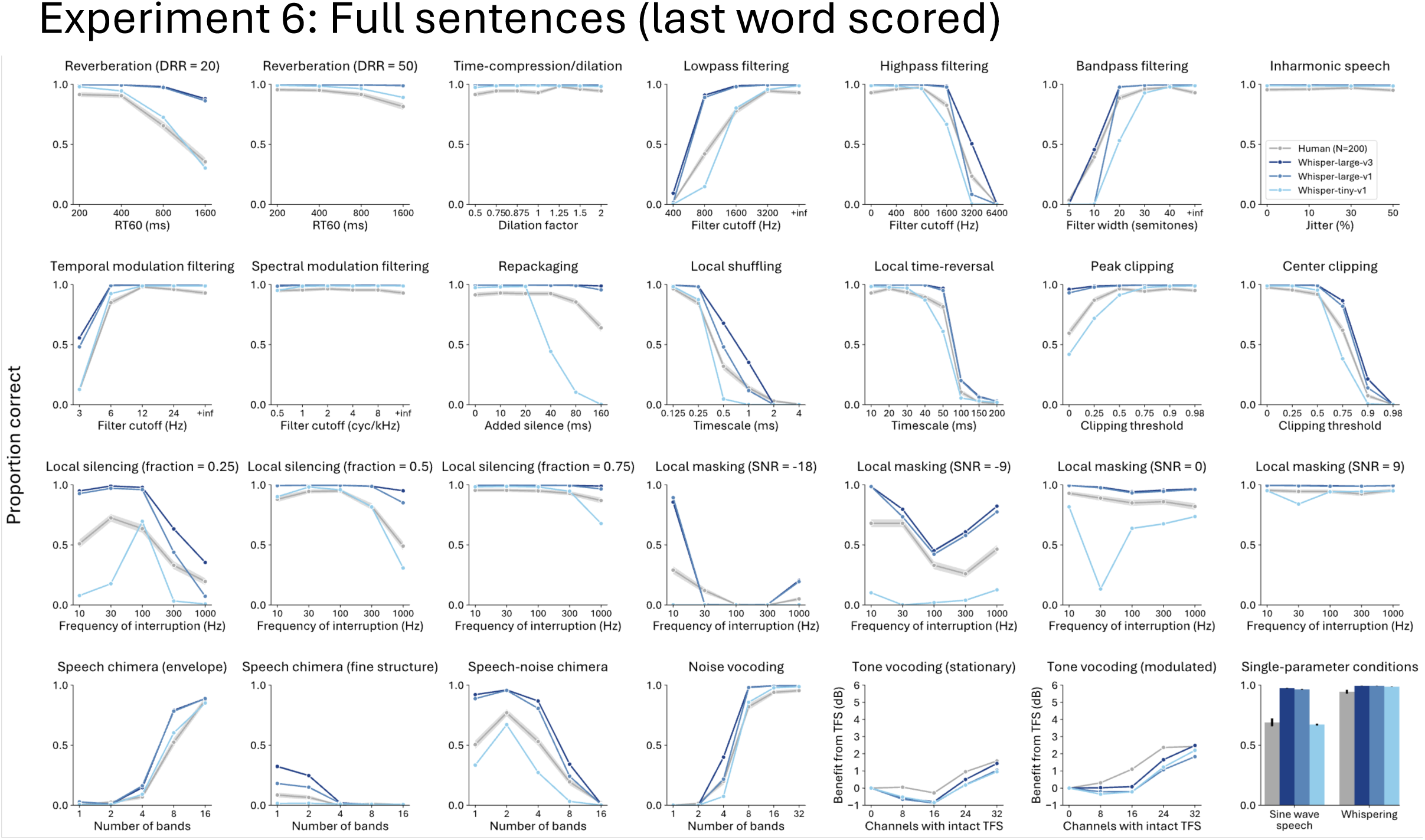
Whisper variants and human results on each distortion type for Experiment 6 (last word scored)

**Supplementary Figure 32:**
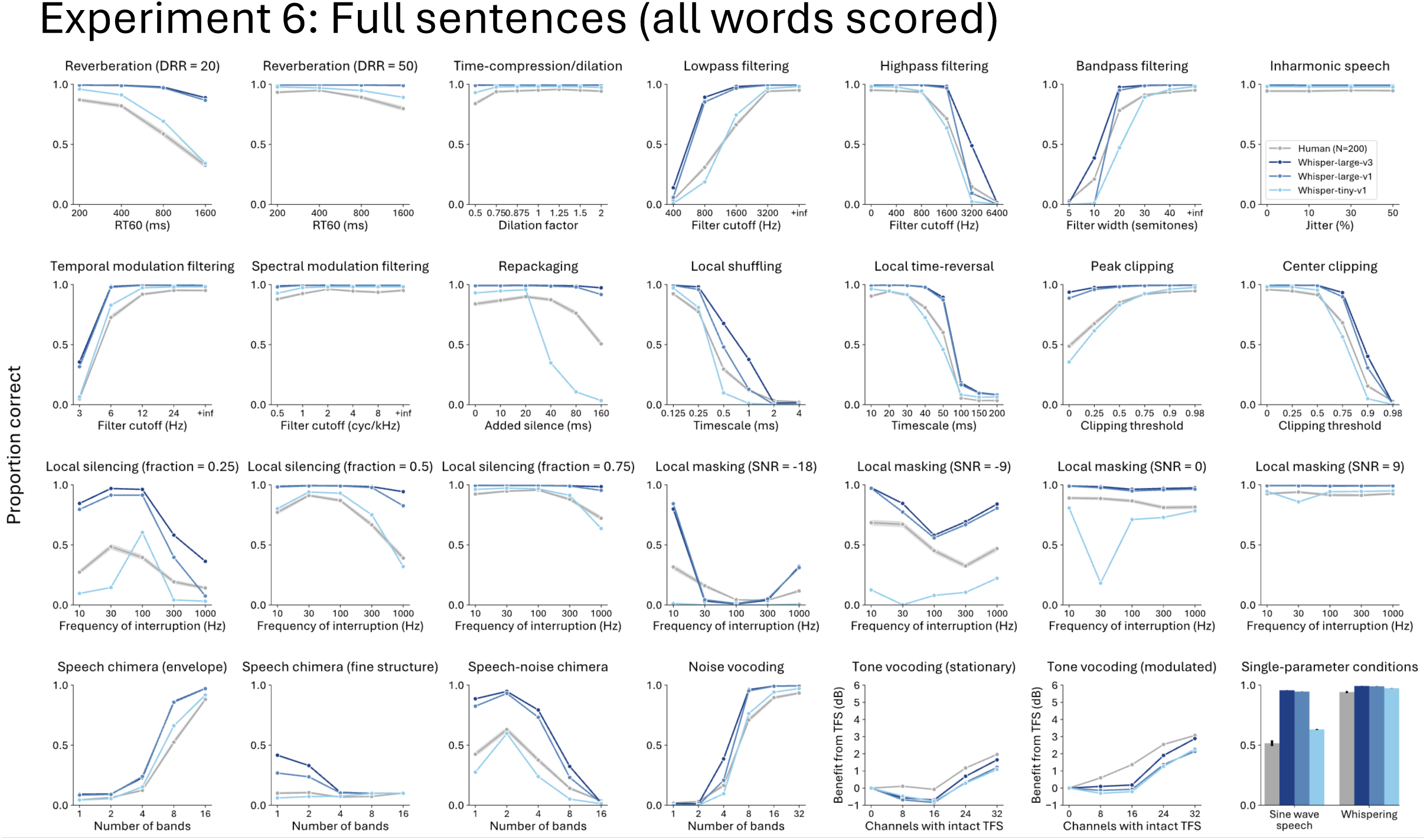
Whisper variants and human results on each distortion type for Experiment 6 (all words scored)

**Supplementary Figure 33:**
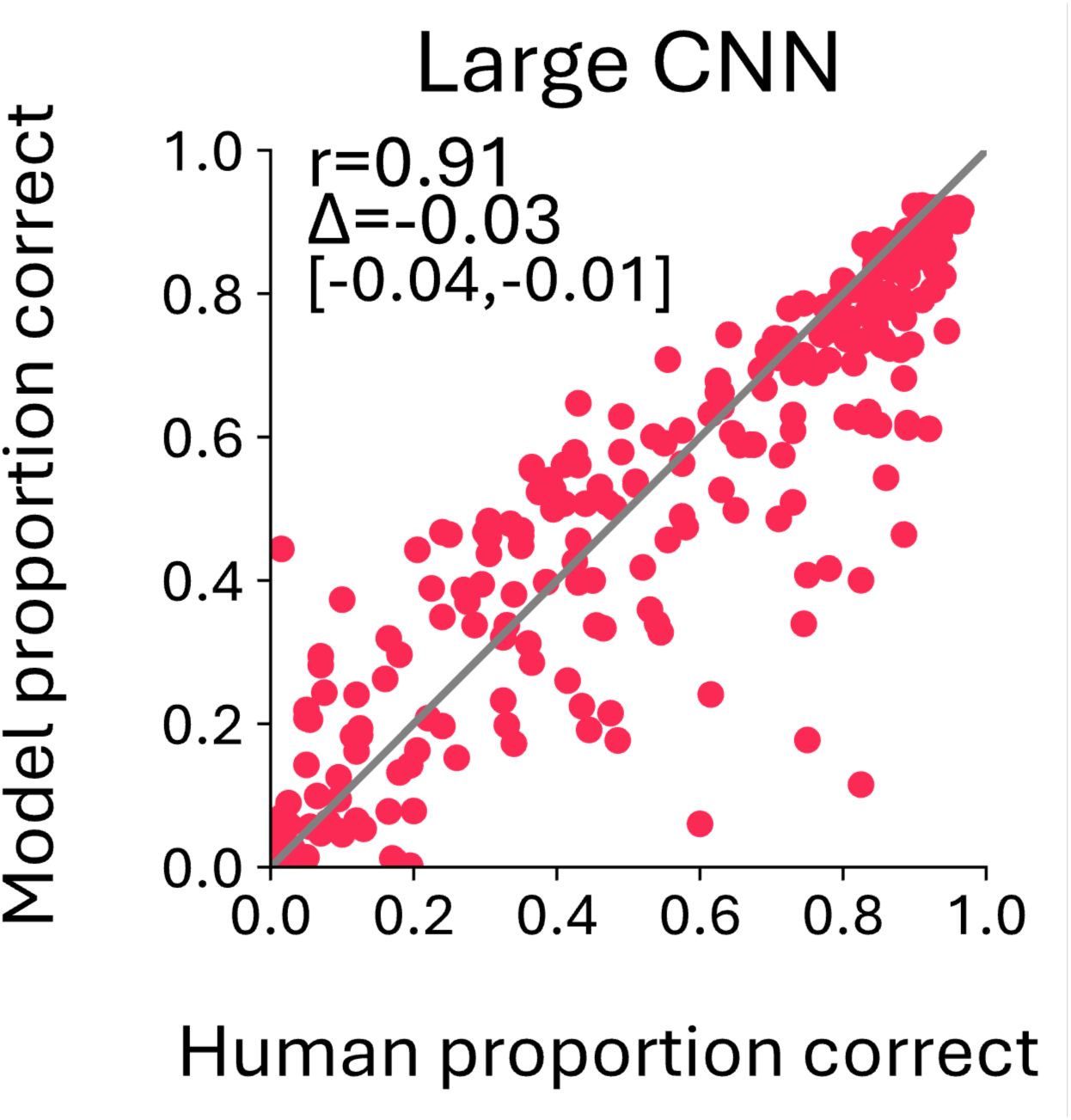
Scatter plot of performance for the Large CNN on Experiment 5

**Supplementary Figure 34:**
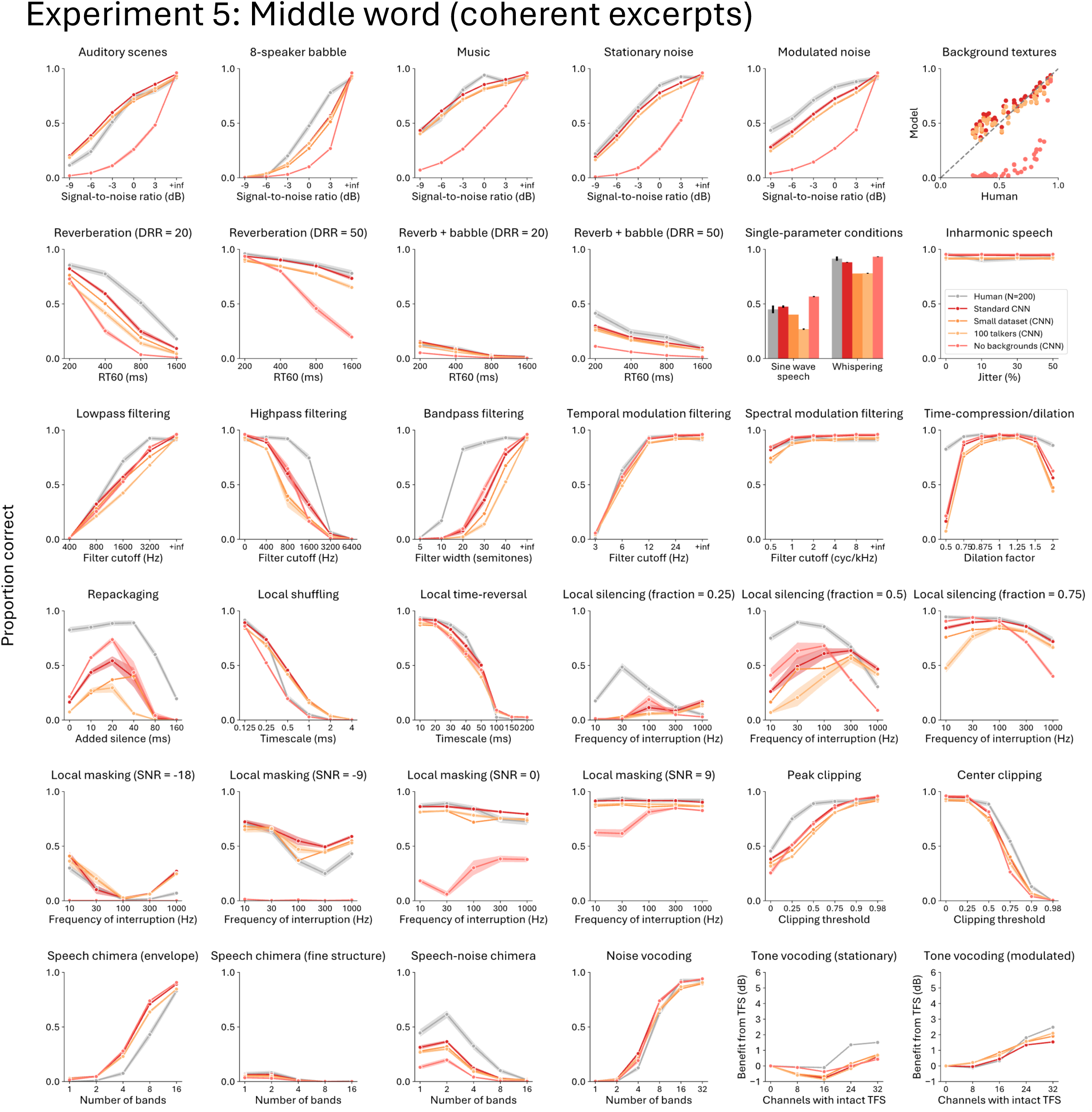
In-house CNNs and human results on each distortion type for Experiment 5

**Supplementary Figure 35:**
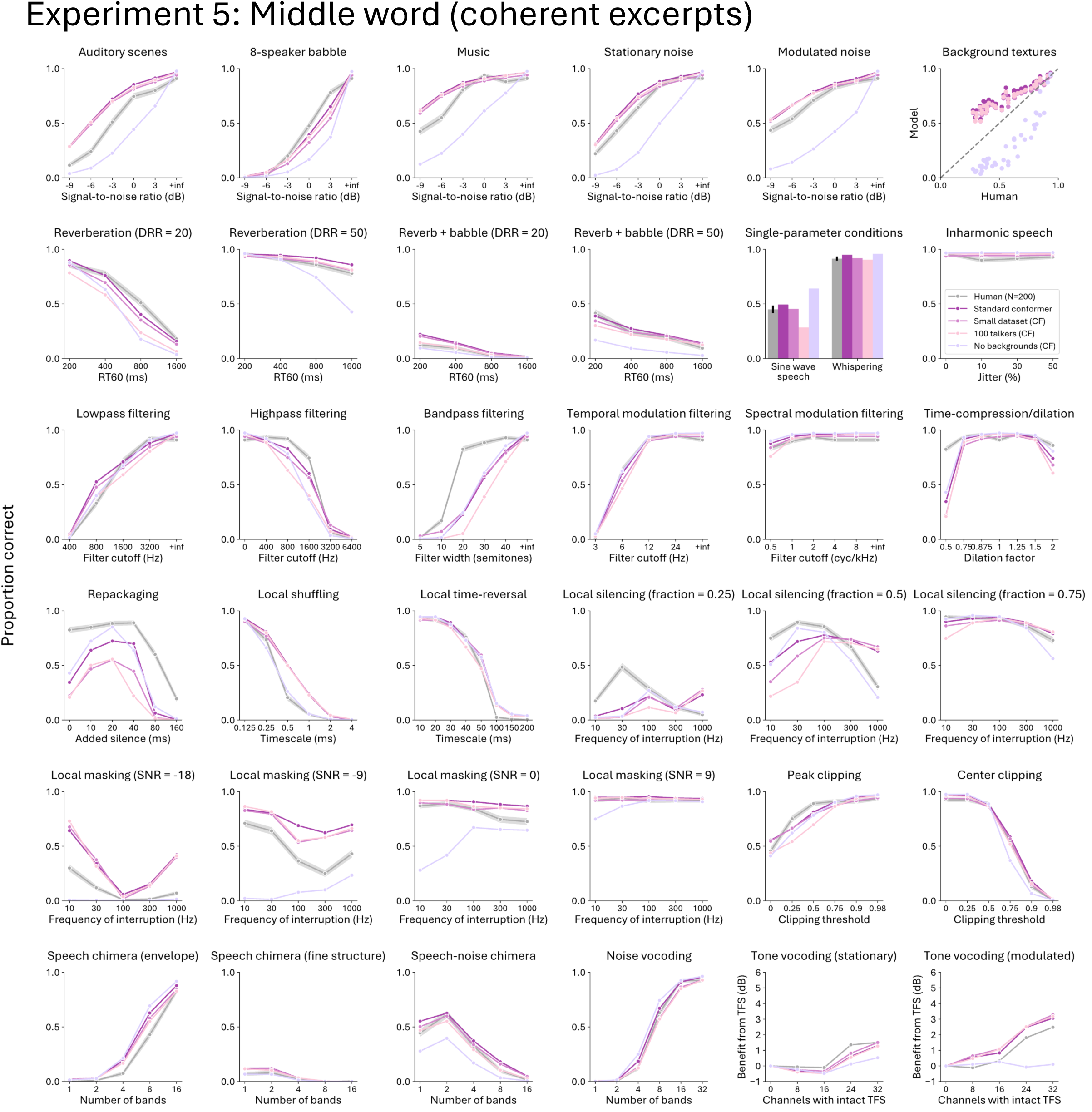
In-house Conformers and human results on each distortion type for Experiment 5

**Supplementary Figure 36:**
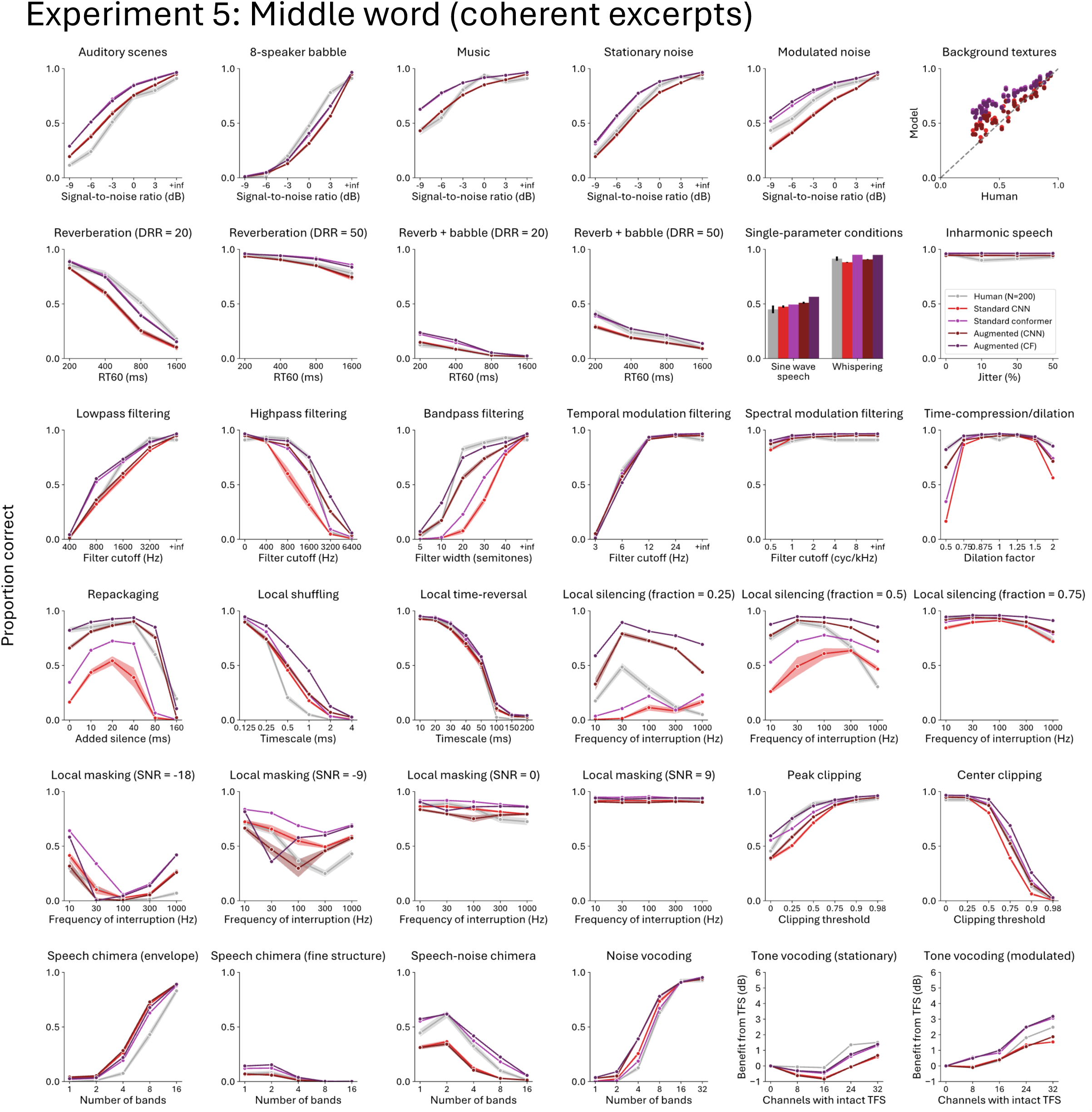
In-house Standard models and Augmented models and human results on each distortion type for Experiment 5

**Supplementary Figure 37:**
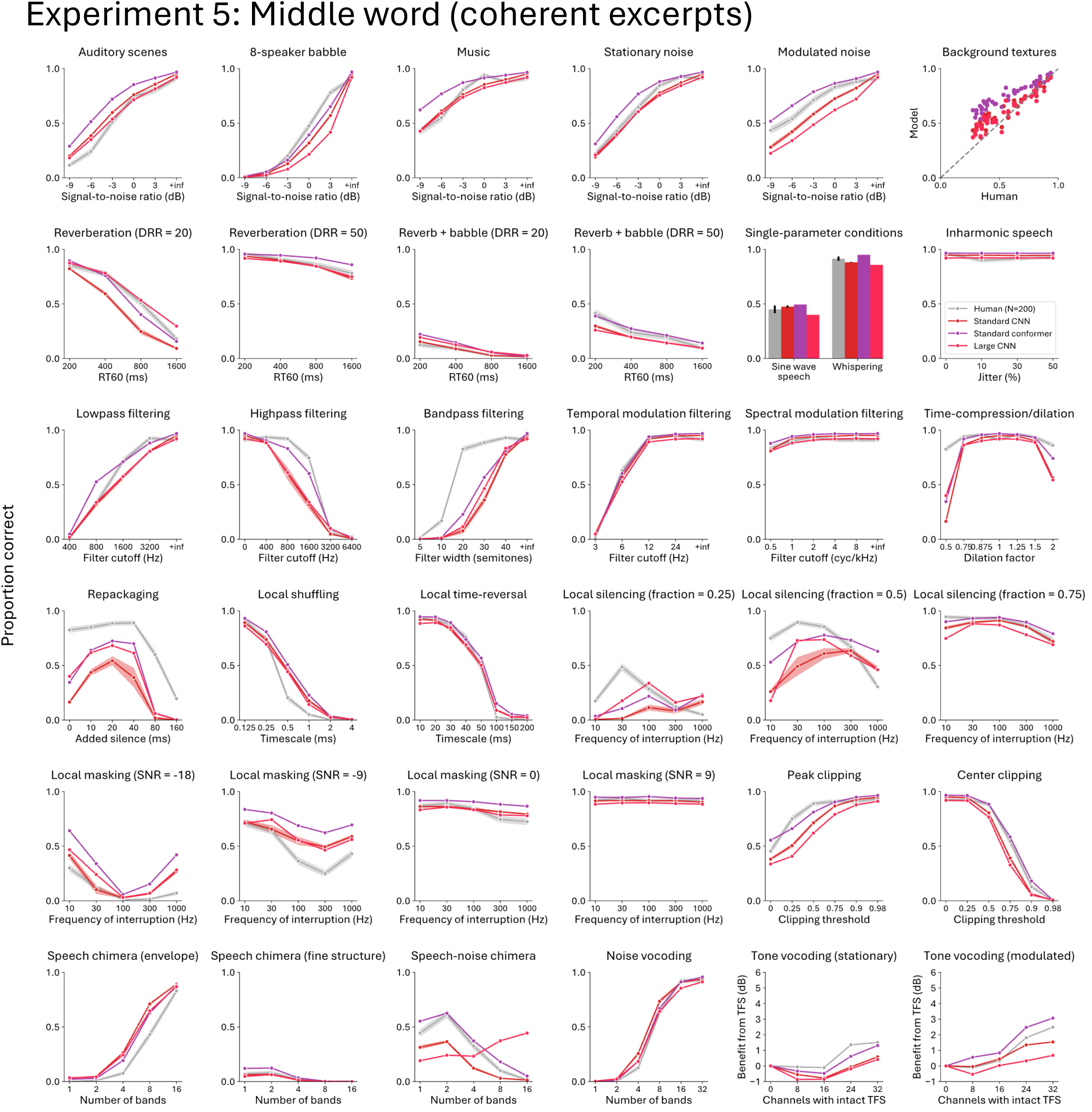
In-house Standard CNN, Standard Conformer, Large CNN and human results on each distortion type for Experiment 5

**Supplementary Figure 38:**
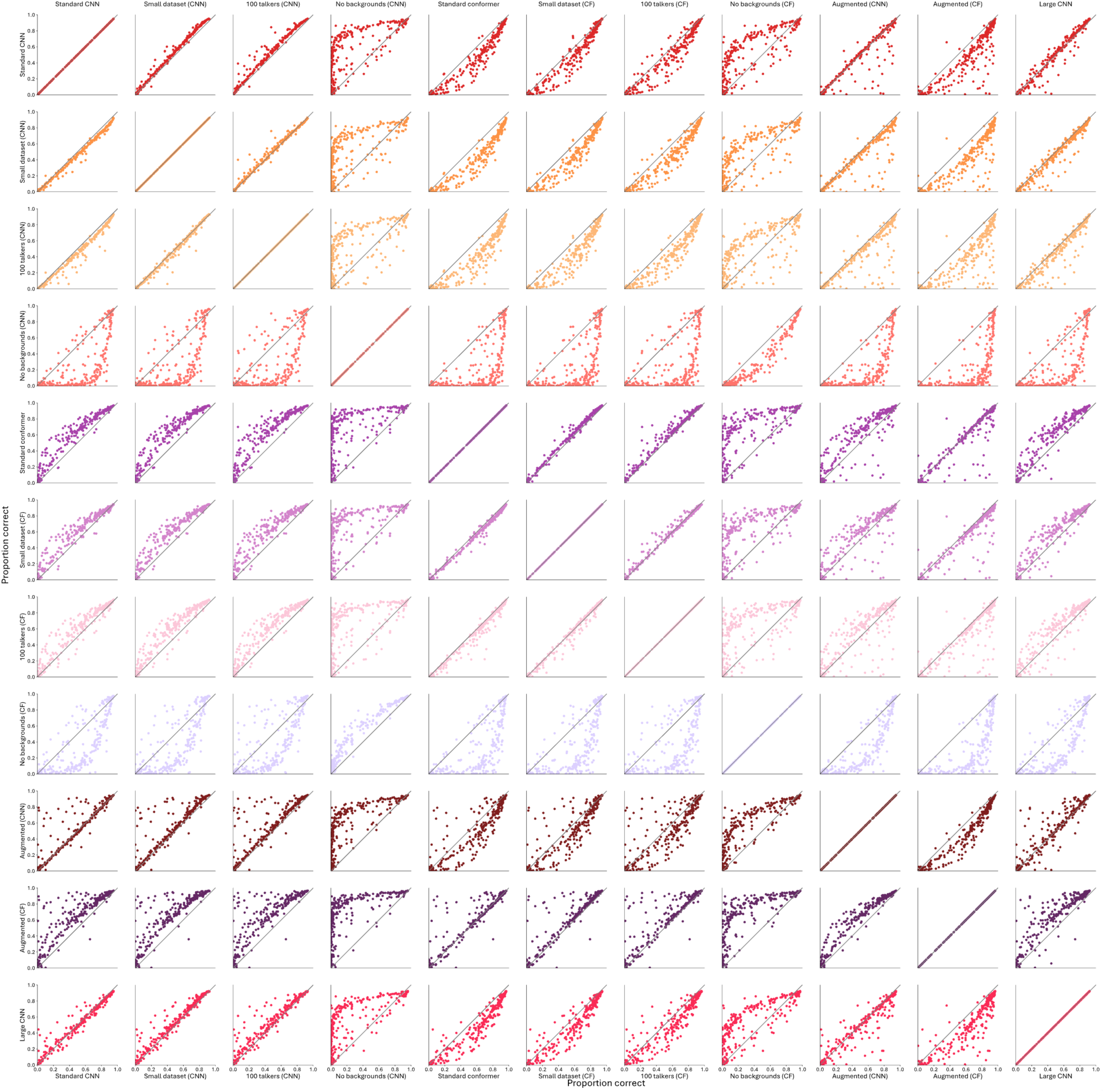
In-house model vs. in-house model comparison scatter plots on Experiment 5

**Supplementary Figure 39:**
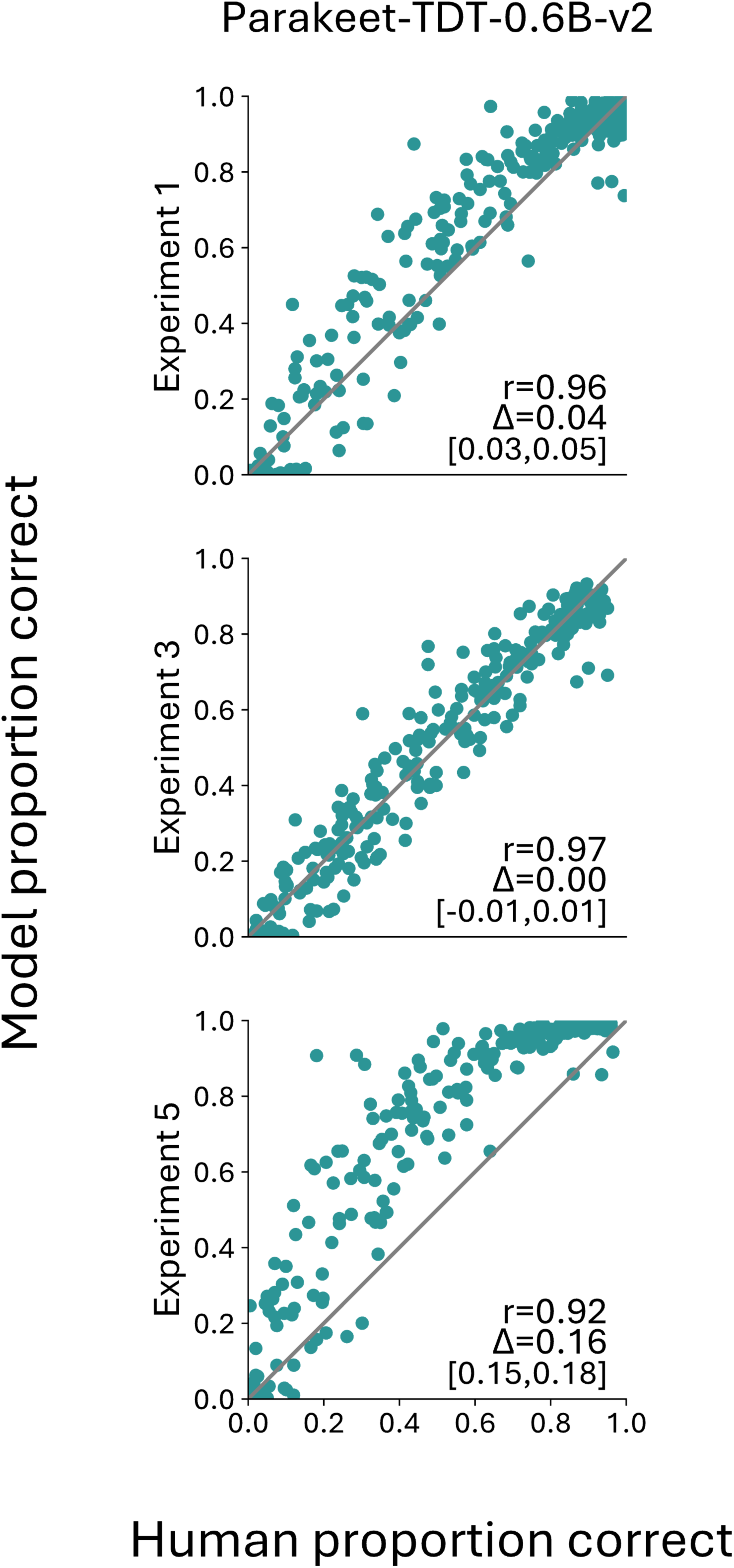
Scatter plots for Parakeet vs. humans restricted to the source clips where Parakeet doesn’t output an empty string

**Supplementary Figure 40:**
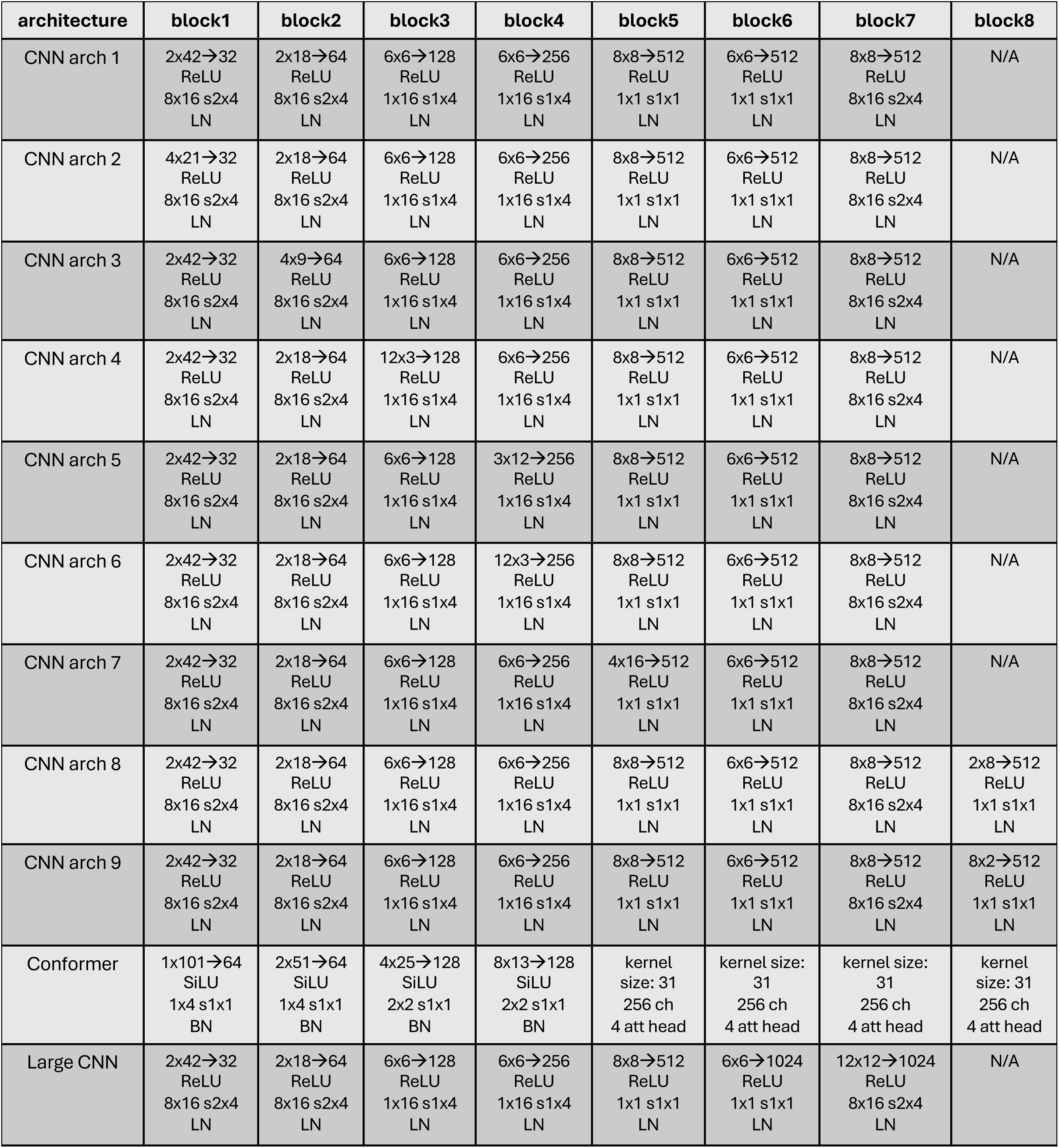
Full specification of in-house model architectures

## Notes

### Competing Interest Statement

The authors have declared no competing interest.

